# Kinetic Proofreading can Enhance Specificity in a Non-enzymatic DNA Strand Displacement Network

**DOI:** 10.1101/2023.08.25.554917

**Authors:** Rakesh Mukherjee, Aditya Sengar, Javier Cabello-García, Thomas E. Ouldridge

## Abstract

Kinetic proofreading is used throughout natural systems to enhance the specificity of molecular recognition. At its most basic level, kinetic proofreading uses a supply of chemical fuel to drive a recognition interaction out of equilibrium, allowing a single free-energy difference between correct and incorrect targets to be exploited two or more times. Despite its importance in biology, there has been little effort to incorporate kinetic proofreading into synthetic systems in which molecular recognition is important, such as nucleic acid nanotechnology. In this article, we introduce a DNA strand displacement-based kinetic proofreading motif, showing that the consumption of a DNA-based fuel can be used to enhance molecular recognition during a templated dimerization reaction. We then show that kinetic proofreading can enhance the specificity with which a probe discriminates single nucleotide mutations, both in terms of the initial rate with which the probe reacts and the long-time behaviour.

## Introduction

Specificity of molecular interactions is at the heart of biology. In particular, selective nucleotide base pairing drives information propagation in DNA replication, RNA transcription and protein translation. Most simply, molecular recognition can be driven by differences in binding free energy between two candidate molecules and a substrate, resulting in an equilibrium with a bias toward one candidate. However, in many biosynthetic processes, the specificity of correct product formation is orders of magnitude higher than that suggested by free-energy differences, despite additional complications that make discrimination more challenging than implied by the equilibrium picture^1^. Translation of protein from mRNA operates with an error rate^2,3^ of 10^−4^, and DNA replication^4^ with an astonishingly low error rate of 10^−9^. To describe such unusually high accuracy, Hopfield^5^ and Ninio^6^ independently proposed “kinetic proofreading” (KP) (Fig. 1a), in which small free-energy differences are utilised repeatedly over multiple steps in a reaction cycle to achieve a significant overall discrimination. The idea has been widely applied, and the same basic principle has been identified in antigen recognition by T-cells^7^, aminoacylation of tRNA^8^, disentanglement of DNA by topoisomerases^9^, specific protein degradation via ubiquitination^10^, and chromatin remodelling^11^.

**Figure 1.**
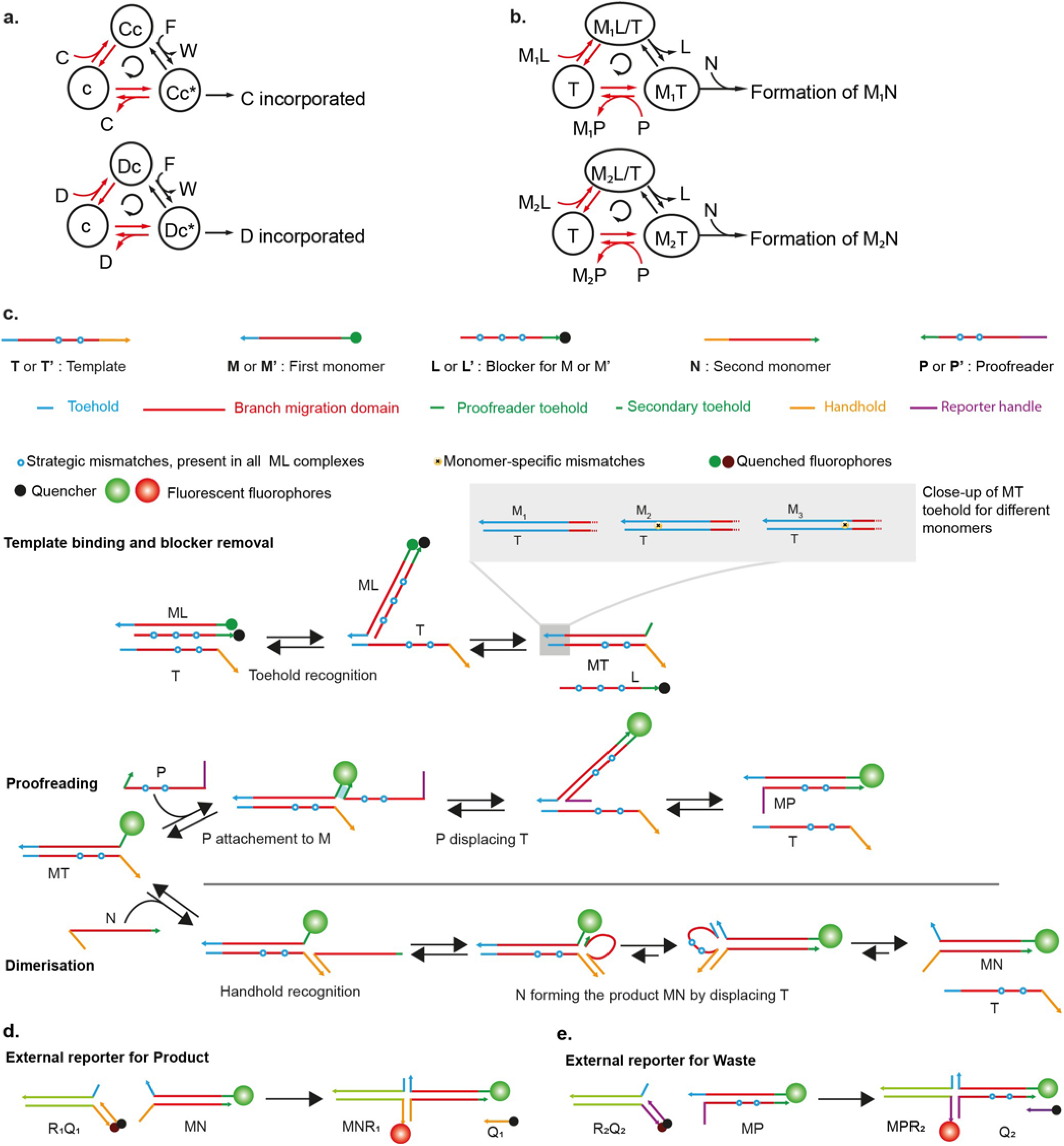
Kinetic proofreading networks. (a) Hopfield’s KP model. Recognition site c can bind to competing molecules C or D. Chemical free energy from the conversion of F(uel) into W(aste) drives the state of the recognition site clockwise around a cycle. The red-coloured arrows show two stages at which discrimination between C and the weaker binding D is expected to occur, leading to a double enrichment of Cc* relative to Dc* and hence an enhanced relative rate at which C is incorporated into a product. (b) Proposed synthetic KP motif, highlighting its topological similarity to (a). The proposed system uses a template *T* to recognize and dimerize monomers *M* and *N*, selecting *M*_*1*_ from the competing variants. Monomer *M* is initially sequestered in a complex *ML*; conversion of this complex into *MP* releases the free energy to drive the template clockwise around the proofreading cycle, allowing discrimination at two stages (red arrows) to be manifest. (c) Domain-level design of a DNA-based kinetic proofreading network. The blocked monomer *ML* binds to the template *T* in two steps via TMSD to form the intermediate *MT* complex. The intermediate can be converted into *MP* waste complex by the proofreader, or dimerized via an HMSD reaction with the second monomer *N* to form the dimeric product *MN*. After either process, template *T* is released to act as a substrate in another reaction cycle. Variants *M*_*2*_ and *M*_*3*_ have a single nucleotide mismatch with the template toehold, which is used at two stages to provide discrimination (red arrows in (b)). Blue circles with white core indicate strategic mismatches that provide thermodynamic drive for all the monomers. The number of such mismatches varies between different competing sets of monomer strands, as noted in the following sections, but remains the same within a set of competing monomers. The invasion of *ML* by *T* is reported by the increase in fluorescence of the monomer strands. Product and waste formation are reported by two external reporters (d-e). Further design details are given in Supplementary Fig. S1-2.

Hopfield’s KP is illustrated in Fig. 1a. Hopfield envisioned a single recognition site c discriminating between two candidate molecules C (correct) and D (incorrect) for incorporation into a product. Either molecule can bind to c, forming Cc or Dc. Then the molecule can either unbind or the complex undergoes a chemical transition to another bound state, Cc* or Dc*. This moment is the first discrimination point: D will have less affinity for c than C and will unbind faster; D is thus less likely to proceed to the second bound state. From the second bound state, there is another opportunity to detach prior to incorporation into the final product, giving a second opportunity to discriminate. The two stages of discrimination can doubly enrich the correct product over the incorrect product.

Central to Hopfield’s KP is the driving of a single recognition site c through a cycle of states (Fig. 1a)^12^. Chemical free energy must be consumed to ensure systematic motion around the cycle. The result is that the Cc*/Dc* ratio can be maintained out of equilibrium, which allows it to be doubly sensitive to the difference in binding energy between the two candidates.

Discriminating similar molecules is also a ubiquitous motif in synthetic nanotechnology, whether in computational strand displacement cascades^13^, tile assembly systems^14^ or diagnostics-based applications^15^. The ability to discriminate between perfectly matching and almost matching sequences is particularly relevant to single nucleotide polymorphism (SNP) detection. SNPs are single nucleotide mutations in specific gene sequences and are markers for various diseases^16^.

Synthetic nanotechnologists have designed elegant systems in which discrimination between intended and unintended complexes is enhanced relative to a baseline^17–21^. Typically, the approach is to modify a design to increase the number of mismatched base pairs that must be formed to make the unintended product. Although such an approach, which may not always be possible in complex networks, implements a form of proofreading, it does not constitute non-equilibrium KP, where a single free-energy difference is exploited multiple times. Indeed, to the best of our knowledge, *de novo* KP has not previously been demonstrated in synthetic systems, despite its importance in nature. This fact suggests a fundamental limitation in both our understanding of, and ability to design, biochemical networks.

In this article, we demonstrate that KP can be implemented in non-enzymatic, DNA-based systems to improve discrimination between very similar recognition domains through a fuel-consuming cycle. Specifically, we apply KP to enhance the specificity with which a molecule is incorporated into a two-stranded DNA dimer by a catalytic template, a synthetic analogue of tRNA charging. We first characterise the individual discrimination steps, then demonstrate that the proofreading motif enhances the overall specificity of dimerization. Finally, we demonstrate the utility of KP by showing that it can enhance a simple network for the detection of generic SNPs in ssDNA.

In so doing, we: a) explore a fundamental biochemical motif by engineering a minimal example *de novo*; and b) demonstrate that it is possible to implement KP in the specific context of DNA strand displacement networks. For the former motivation, constructing a minimal synthetic KP motif tests our understanding of the fundamental biochemistry, potentially revealing subtleties that are absent in idealised models. For the latter, we open the door to the application of KP as a general mechanism to a range of DNA strand displacement-based systems^22^ that need to distinguish between correct and incorrect products with similar free energies. Since KP is a way of *improving* the specificity of a recognition process, it can be applied even to systems that already discriminate quite well, allowing the intended targets to be distinguished from even larger pools of competing molecules.

## Materials and Methods

### Sequence design and nomenclature

All sequences were generated using the NUPACK web server to minimise undesired secondary structures. Some strand designs were manually altered to incorporate strategic features such as mismatches. All the strands were purchased from Integrated DNA Technologies as HPLC purified at 100 µM concentrations in IDTE buffer, pH 8. All the sequences used in the paper are listed in Tables S15-17. Detailed design considerations are shown in Supplementary Fig. S1-S2.

The first type of monomers named *M*_*1*_, *M*_*2*_, and *M*_*3*_ are used for characterising monomer binding, sequence-specific proofreading, and complete discard mechanism. *M*_*1*_ is the “correct” monomer, and *M*_*2*_ and *M*_*3*_ are the “incorrect” strands. Blocker strand *L* is used for the monomer-blocker duplex. *P*_*6*_, *P*_*7*_, and *P*_*8*_ are the proofreader strands used for the set of monomers. An altered design of the first type of monomers, *M’*_*1*_, *M’*_*2*_, and *M’*_*3*_ was used for the final templated dimer formation along with the second type of monomer *N*. Similarly, altered template *T’*, blocker *L’*, proofreaders *P’*_*6*_, *P’*_*7*_, and *P’*_*8*_ were used with the second set of monomers. Generic terms *M, P, T, R* were used to denote all strands belonging to one of these categories.

External fluorescence reporters, denoted generically as *RQ*, as shown in Fig. 1d-e and Fig.5a, are formed by three strands. The strand bearing the quencher species is denotes as *Q*; other two partially complimentary strands are collectively denoted as *R. R*_*1*_*Q*_*1*_ is the porter for the dimeric product *M’N* and *R*_*2*_*Q*_*2*_ for the *MP* wastes.

### General annealing protocol

The *ML* and *M’L’* duplexes were prepared by annealing 100 nM *M* or *M’* with 20% excess *L* or *L’* to ensure that all *M* or *M’* strands were bound to *L* strands. To prepare *MT* and *MP* complexes, 100 nM *M* strands were annealed with 20% excess *T* or *P* strands. Reporter complexes *RQ* were comprised of three strands, *Rcomp, r*_*i-j*_ (collectively called *R*) and *Q* (details in Supplementary Fig. S2, Supplementary Tables S15-17). For this, 200 nM *Rcomp* strands were annealed with 20% excess of *r*_*i*-*j*_ and *Q* strands. The volumes of the annealing mixtures were 100 or 200 µL depending on the experimental requirements. The required strands were mixed to have the final concentrations in 1x Tris-Acetate-EDTA buffer containing 1M NaCl. This mixture was first heated to 95°C and held there for 5 minutes, then gradually cooled down to 20C at 1°C per minute and stored at 4°C until used.

### Fluorescence Kinetics measurements and calibration

All fluorescence measurements were performed in a BMG Labtech CLARIOstar fluorescence plate reader in Greiner µClear flat-bottomed 96-well plates. Reactions were initiated, unless otherwise stated, by injecting 50 µL of trigger (usually a combination of single stranded or duplexed DNA oligonucleotides with buffer) into 150 µL of the other reactants using the instrument’s in-built injectors (pump speed 430 µL/ sec). The final mixtures were then shaken for 10-30 seconds at 400-500 rpm before measuring the fluorescence.

Monomers *M*_*1*_, *M*_*2*_, *M*_*3*_, *M’*_*1*_, *M’*_*2*_, *M’*_*3*_, and *Rsnp* were labelled with fluorophore Cy3. *L, L’*, and *Qsnp* were labelled with IowaBlack FQ quencher. *Rcomp-AF* and *RcompP* were labelled with AlexaFluor-647. *Q*_*1*_ and *Q*_*2*_ were labelled with IowaBlack RQ quencher.

For the Cy3 and AlexaFluor-647 measurements, presets from the instruments were used for excitation and emission wavelengths. Specifically, for Cy3, excitation: 530-20 nm, emission: 580-30 nm, gain: 2100; and for AlexaFluor-647, excitation: 625-30 nm, emission: 680-30 nm, gain 2700). For each well, each measurement was taken as 20 flashes in a spiral scan with a maximum diameter of 4 mm. Step-by-step experimental methods used in each experiment are given in Supplementary Notes 1-8.

Fluorescence calibrations were performed for fluorophore-labelled complexes to quantify the fluorescence signals obtained from unit concentration of such complexes. Cy3 (for *ML, MP, MT* complexes) and AlexaFluor-647 (for *RQ* and *MPR* complexes) signals were monitored in a volume of 200 µL with different candidates’ concentrations ranging from 0-20 nM in 1x TAE buffer with 1 M NaCl. Only *M*_*1*_ and *P*_*6*_ were used rather than testing all possible permutations of monomers and proofreaders because the local environments of the fluorophores in all complexes were same.

### Data Processing and analysis

This section describes the methods used for the estimation of the reaction rate constants for the various intermediary reactions. All rate constants are obtained from the experimental reaction dataset as obtained from the Cy3 and AlexaFluor-647 measurements shown in Supplementary Fig. S16, S19, S22, S26. The AlexaFluor-647 channel gives a fluorescent signal in the presence of *RQ, MPR*, and *PR* whereas Cy3 gives signal for *ML, MT, MP* and *MPR*.

### Trajectory analysis and rate constant estimation

We used the ParametricNDSolve built-in function in Mathematica 12.3 to fit mass action models of reaction kinetics to the processed data. The ODE models used to describe each experiment are given in Supplementary Notes 10 – 13. In general, the procedure involved fitting a single set of rate constants to a number of kinetic traces under distinct initial conditions. In each case, we defined an array *K*_*n*_ of known rates (determined by earlier fits to different experiments) and an array *K*_*un*_ of unknown rates. The system of ODEs was constructed within the ParametricNDSolve function, and all the starting conditions, initial concentration values and *K*_*n*_, were provided. ParametricNDSolve was then used to minimize the mean squared error (MSE)

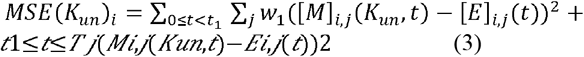

over possible values of *K*_*un*_. Here, [*M*]_*i,j*_(*K*_*un*_, *t*) is the predicted concentration of the chosen species *i* in kinetic trace *j* at time T, given *K*_*un*_ as the values of the unknown rate constants. Concurrently, [*E*]_*i,j*_ (*t*) corresponds to the data observed in experiment. To improve the accuracy with which fits captured the behaviour at early time points, weights *w*_*1*_=1 and *w*_*2*_=0.2 were assigned to data points with *t* < *t*_l_ and *t*_l_ 5: *t*, respectively. *t*_*1*_ is approximately equal to the time instance when the species concentration has plateaued and was manually defined for each reaction. Each element in the set *K*_*un*_ is sampled from a variable range of values that can be found in Supplementary Notes 10-13.

The data on the binding of blocked monomers to templates was used for identifying rate constants governing the displacement of blockers from monomers and vice versa. The procedure is outlined in Supplementary Note 10. The fitted parameters can be found in Table S9 and the comparison between experimental observations and simulation predictions is shown in Supplementary Fig. S32. Subsequently, data pertaining to the triggering of reporter complexes by pre-prepared MP duplexes was utilized to determine rate constants for the reporter, as explained in Supplementary Note 11. Due to challenges in unequivocally determining the total monomer concentration in these reactions, eight distinct sets of rate constants for each reporter were generated, based on plausible estimates of the total monomer concentration. These eight sets of reporter rate constants, in conjunction with the rate constants for template binding, were employed in a simultaneous fit of the template recovery and full discard pathway experiments. The set of rate constants that minimized the total error for template recovery and discard pathway was then selected. The procedure is outlined in more detail in Supplementary Note 11. The resulting fits for reporter characterization are shown in Supplementary Fig. S34-36; the fits for template recovery are shown in Supplementary Fig. S37-39; and the fits for discard pathway are shown in Supplementary Fig. S40-S42. The fitted rate constants are given in Tables S10-11.

### Model-based predictions for intermediates and dimerization

The values of rate constants obtained from fits were used to predict the time-varying concentration of *MT* complexes during the full discard pathway experiments, Supplementary Fig. S43. The predictions were made by numerically integrating Eqs. S35-S39 using the rate constants in Tables S9-S11 and the experimental initial conditions as parameters.

Although the dimerization system used re-designed monomers and templates, we nonetheless used the rate constants obtained from initial fits to predict the yield of dimers over time. The predictions were made by integrating Eqs. S51-S53 using the rate constants in Tables S9-S11 and the experimental initial conditions as parameters. A plausible dimerization rate constant of 3.50E+05 M^-1^ s^-1^ was also used. The results are plotted in Supplementary Fig. S53. At variance with the experimental results, the model predicts a high dimerization yield for all monomers and proofreaders in the long-time limit. We suspect that this prediction is due to an excessive reverse proofreading (rebinding of proofread monomers to the template) rate constant for mismatched monomers; these constants are not well-constrained by the data.

## RESULTS

### Network Design for Kinetic Proofreading

We construct an enzyme-free, DNA-based synthetic KP system. DNA is used purely as an engineerable molecule that predictably assembles into well-defined structures driven by Watson-Crick-Franklin base pairing. Our KP system exploits the widely-used motif of toehold mediated strand displacement^22–25^ (TMSD). In TMSD, a single invader strand displaces an incumbent from an incumbent-target duplex (see Fig. 1c). The process is accelerated and pushed thermodynamically downhill by a toehold of available bases in the target to which only the invader can bind. Alongside TMSD, we also exploit the handhold mediated strand displacement^26–28^ (HMSD) reaction (also shown in Fig. 1c). HMSD is analogous to TMSD, but the binding to the target is accelerated by a handhold in the incumbent rather than the target.

The network is illustrated at a domain level in Fig. 1c. Domains are sections of single-stranded (ss)DNA that are designed to bind as a collective unit. In this network, the monomers *M* and *N* that form *MN* are each single strands of DNA (rather than individual nucleotides or amino acids). Dimerization in solution is inhibited by the presence of a blocker strand *L* bound to *M*, but can occur rapidly via a template T, which acts as the substrate in our system. First, using a toehold (t_T_), *M* can bind to *T* via TMSD and remove *L*, revealing a second short toehold domain (t_P_)^29^. The newly available toehold t_P_ can be used by the proofreader strand *P* to initiate a second TMSD reaction to form a waste duplex *W*, or a second monomer *N* can bind to the handhold h and then complete HMSD to form a dimeric product *MN*. Effectively, the template *T* catalyses two competing non-covalent processes: a two-step toehold exchange^30–32^ in which *ML + P* is converted into *MP + L*, and an HMSD-mediated dimerization^26–28^ with *ML + N* being converted to *MN + L*.

From the perspective of the template, the process is a direct analogue of Hopfield’s KP (compare Fig. 1a to Fig. 1b). The free energy released by converting *ML + P* into *MP + L* drives the template round the sequence of states *T*→*ML/T*→*MT*→*T*, with conversion to the final product MN happening from the MT state. This nonequilibrium cycling allows for the same difference in binding free energy between three versions of the first monomer: *M*_*1*_, *M*_*2*_ and *M*_*3*_, to be exploited at two stages. These monomers differ by a single base in the toehold recognition domain t_T_; *M*_*1*_ has a perfectly matching toehold for template recognition, whereas *M*_*2*_ and *M*_*3*_ have one mismatch at different positions in the toehold. First, after the initial binding of *M* to *T*, the strength of the toehold t_T_ determines the probability that the monomer will remain bound for long enough for blocker displacement

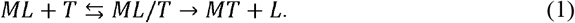

Second, a stronger toehold interaction can also reduce the probability of successful displacement by the proofreader^33^,

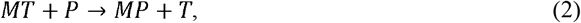

allowing for the concentration of *M*_*1*_*T* to be further enriched relative to *M*_*2*_*T or M*_*3*_*T* and thereby enhancing the relative rate at which *M*_*1*_*N* is formed.

Although the synthetic KP motif is equivalent to Hopfield’s KP motif from the perspective of the template, there is an important difference. In our design, the thermodynamic drive pushing *T* around the proofreading loop comes from converting the monomers themselves into waste, rather than ancillary fuel molecules. We will return to the consequences of this difference in the conclusion.

Complete design details, including sequences, are provided in Supplementary Fig. S1 and Table S15-16. Long binding domains (red domain, Fig. 1c) contain mismatched base pairs (blue circles) in the initial *ML* complexes; these mismatches are gradually eliminated during subsequent reactions. The mismatches are common to all monomers and provide a “hidden thermodynamic drive”^33^ towards formation of the final product *MN*.

We initially consider a fixed set of monomer, template, and blocker strands. We explore the performance of the individual substeps and overall network for different designs of the proofreader strand.

### Monomers binding to the template

We consider the rates of invasion of blocked monomer complexes *ML* by the template *T*. The monomers were labelled with Cy3 fluorophore, and the blocker strand *L* was labelled with quencher IowaBlack-FQ. The progress of the reaction was monitored by the increase in Cy3 fluorescence when *L* was displaced by *T* (Fig. 2a). We performed the reaction with a range of *T* concentrations for 8nM of *M*_*1*_, *M*_*2*_ and *M*_*3*_; the results for [*T*]=10nM are plotted in Fig. 2b, showing the correct monomer *M*_*1*_ forming a duplex with *T* faster and with a higher equilibrium yield^15^ than the mismatched monomers (Supplementary Fig. S32). Solid lines in Fig. 2b show fits to an ODE model for each system (Supplementary Note 11); these fits used a single set of parameters for each system to fit all concentrations of *T*.

**Figure 2.**
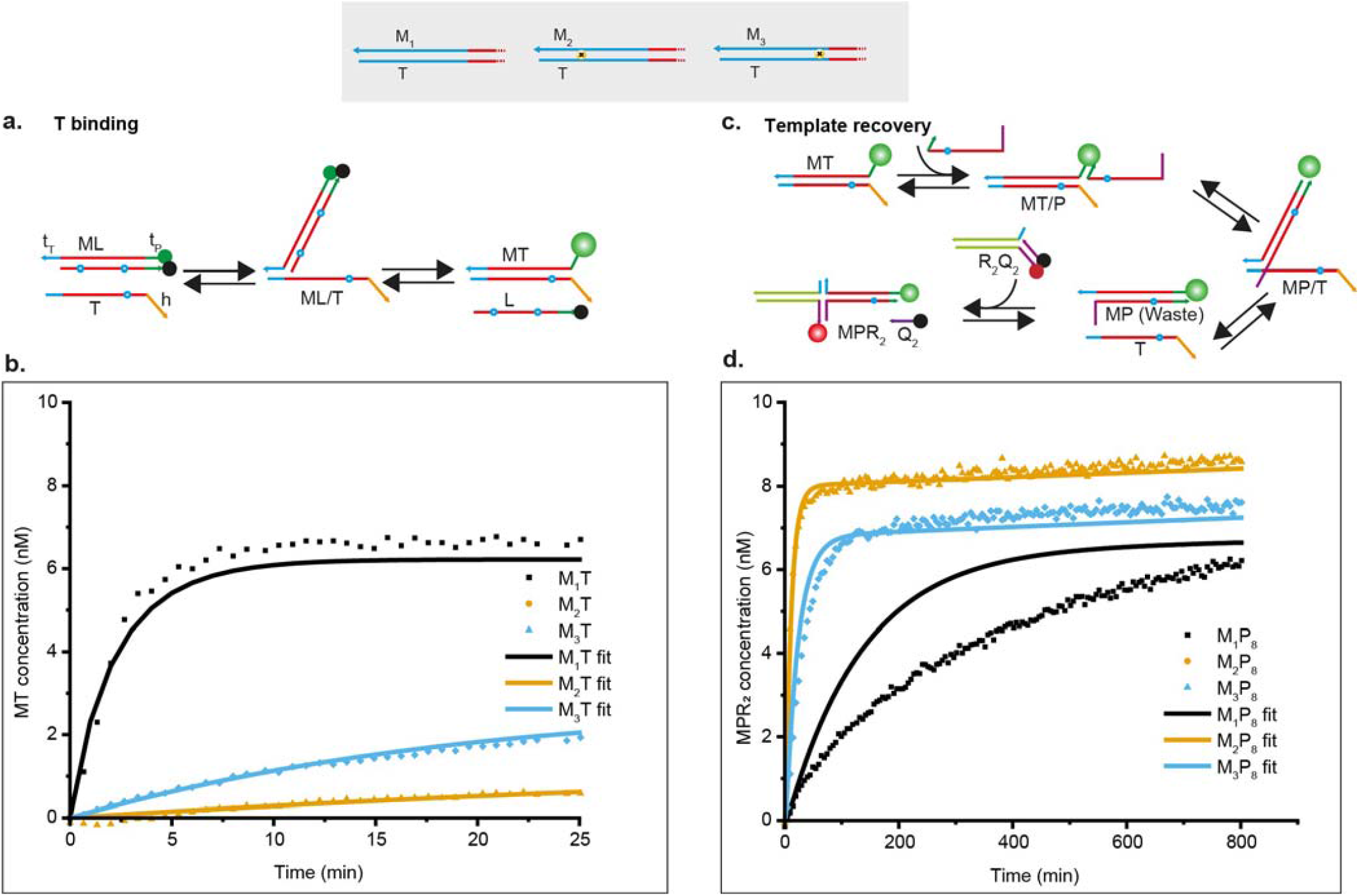
Two stages of discrimination in the KP experiment. (a) Schematic of template binding showing that template *T* invades blocked monomer *ML* by TMSD to form *MT*. (b) Kinetics of the template binding process obtained from the Cy3 signal of the *M* strand. 8 nM *M*_*1*_*L, M*_*2*_*L*, and *M*_*3*_*L* were added in separate wells and the reactions were initiated by injecting 10 nM *T* in each well. Obtained fluorescence signals are converted into concentrations using calibration curves of known concentrations (Supplementary Note 11). Solid lines are fits to an ODEmodel fitted to several experimental conditions simultaneously. (c) Template recovery is the removal of *M* from *T* by the proofreader molecule *P*. Formed waste *MP* is reported by the external reporter R_2_Q_2_, which is labelled with AlexaFluor-647. (d) Kinetics of the template recovery step were obtained from the AlexaFluor-647 signals. 8 nM *M*_*1*_*T, M*_*2*_*T*, and *M*_*3*_*T* and 20 nM of their respective reporter complex were added in separate wells. 50 nM *P*_*8*_ was added in each well to trigger the reaction. Solid lines are fits to an ODE model (Supplementary Note 12) fitted to several experimental conditions simultaneously (Supplementary Fig. S37-39).

### Sequence-specific proofreading

As a second step, we looked at the efficacy and sequence-specificity with which proofreader strands *P* remove monomers from the template, allowing “template recovery” to participate in another reaction. We explored three different designs for *P*; each had the same 5-base toehold t_P_, but we modified the thermodynamic strength of the *MP* waste product by truncating the displacement domain of *P* (shown in red in Fig. 2c). We label the designs *P*_*6*_, *P*_*7*_ and *P*_*8*_, with the 6, 7, and 8 being the number of bases in the toehold and displacement domain of *M* that are left unpaired after the proofreading reaction. *P*_*6*_ thus represents the most thermodynamically favoured proofreader, and *P*_*8*_ the least. In all cases, reactions were driven by adding large excess of *P*, which constitute a fuel reservoir. For all combinations of proofreaders and monomers, a range of concentrations of *MT* complexes were added to a large excess of proofreaders; the formation of *M*_*1*_*P* waste was reported by an external reporter *R*_*2*_*Q*_*2*_, which was labelled with AlexaFluor-647-IowaBlack RQ fluorophore-quencher pair and was already present in solution (Fig. 2c, Supplementary Note 4, supplementary Fig. S34-36). The results for *MT*= 8nM and *P*_*8*_ are shown in Fig. 2d; full results are given in Supplementary Fig. S37-39.

Again, we also report fits to an ODE model for each system; these fits used a single set of parameters for each proofreader, fitted simultaneously to all template recovery experiments as well as subsequent experiments on the full proofreading cycle. As expected, *P*_*6*_, which had the strongest binding to *M*, was the fastest in forming the waste duplex *PM*. Other proofreaders were slower, but still demonstrated template recovery. Crucially, *P*_*7*_ and P_8_ showed clear discrimination between *M*_*1*_, *M*_*2*_ and *M*_*3*_, with *M*_*1*_ being removed from *T* much more slowly, as evident for *P*_*8*_ in Fig. 2d and borne out by the fitted rate constants (Supplementary Table S11). Alongside the discrimination present in the initial template binding, the existence of this second discrimination step demonstrates the potential for kinetic proofreading. For *P*_*6*_, removal by proofreader appears slightly slower for *M*_*1*_ than *M*_*2*_ and *M*_*3*_, but the distinction is smaller, and the relatively slow reporter rate complicates the analysis.

The action of the proofreaders is fundamentally different from the presence of a large reservoir of blocker strands. Adding a large concentration of blocker monomers to *MT* complexes does result in the removal of *M* and the recovery of T, as shown outlined in Supplementary Note 2 and Fig. S33. However, removal of *M*_*2*_ and *M*_*3*_ is not substantially faster than *M*_*1*._ The relative rates for template displacing blocker and blocker displacing template are fundamentally constrained by the freeenergy change of the process. The overall discrimination is equilibrium in nature, and if the full equilibrium discrimination is manifested in the rate of template binding, no discrimination will be observed in the reverse reaction^34^. In contrast, using a distinct proofreader strand allows, in principle, the initial template binding and subsequent template recovery steps to be thermodynamically decoupled by fuel consumption. In practice, we rely on a single additional mismatch between blocker and template to manifest this difference; previous work has shown that a well-placed single mismatch can have large kinetic and thermodynamic effects.

### Complete discard mechanism

We now analyse the entire catalytic discard mechanism in which the blocked monomers first bind to the template to form *MT* duplexes, and are subsequently converted into waste complexes by *P* (Fig.3a). We added *T* to solutions of *M*_*1*_, *M*_*2*_ or *M*_*3*_, *P*_*6*_, *P*_*7*_ or *P*_*8*_, and the appropriate waste reporter, and tracked both the amount of monomers that had been liberated from blocker strands and the amount of waste in solution. Results for *P*_*8*_ and *T*= 4 nM and are shown in Fig. 3b, 3c, and full results in Supplementary Fig. S40-42. Fits of the data to an ODE model were performed simultaneously with the data on template recovery, using a single set of parameters for each proof-reader (Supplementary Note 13).

**Figure 3:**
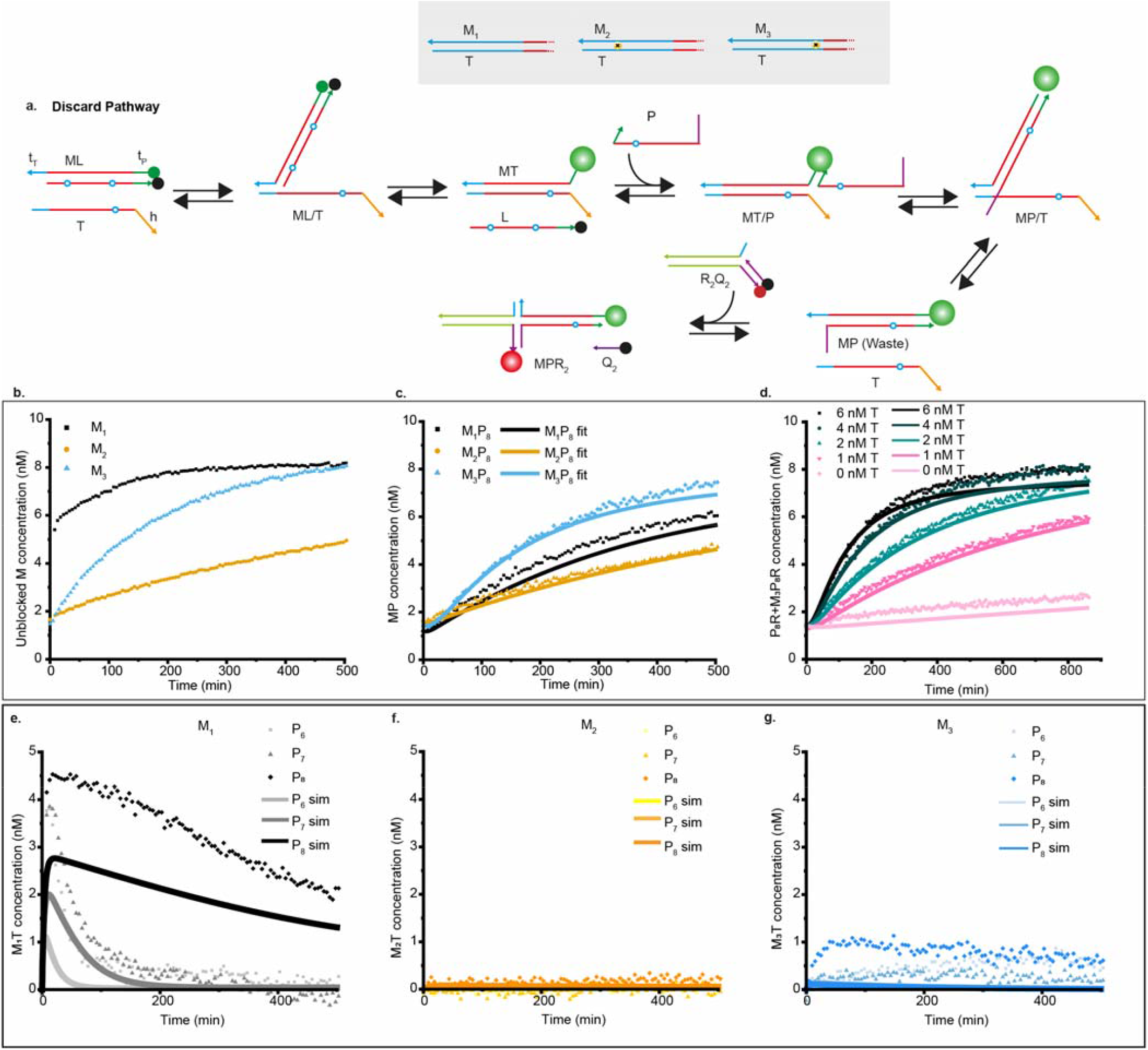
Kinetics of the full discard pathway. (a) The full discard pathway starts with blocked monomer, proofreader and reporter, and is triggered by the injection of *T*, which displaces the blocker from *M* and is subsequently displaced itself by the proofreader strand. (b) Concentration of *M* liberated from its blocker after injection of 4 nM *T*, as inferred from Cy3 fluorescence, for 8 nM *M*_*1*_*L, M*_*2*_*L* and *M*_*3*_*L* in different wells, each with 50 nM *P*_*8*_ and 20 nM of their corresponding reporters. (c) Formation of *MP* waste in the same reactions, as inferred from reporter fluorescence. Solid lines are fits to an ODE model fitted to several experimental conditions simultaneously (Supplementary Note 13). (d) Fluorescence of reporter for *M*_*3*_*P*_*8*_ waste after triggering a mixture of *M*_*3*_*L* and *P*_*8*_ with different concentrations of T. Rates of waste formation increased with increased concentrations of *T*. For all cases, formed waste concentrations are higher than added template which indicates repeated catalytic activity. (e) Concentration of intermediate *M*_*1*_*T* over time for three different proofreaders, *P*_*6*_, *P*_*7*_, and *P*_*8*_. Concentration of *M*_*1*_*T* was estimated by subtracting AlexaFluor647 signals from Cy3 signals converted to concentration. (f, g) Concentration of *M*_*2*_*T* and *M*_*3*_*T* over time in equivalent experiments. Solid lines in (e, f, g) are estimates of *MT* from the fitted ODE model.

The speed of *MP* complex formation depends on both the rate of initial template binding and proofreader-driven template recovery. The first step is faster for *M*_*1*_, and the second step is faster for *M*_*2*_ and *M*_*3*_. More importantly, the system exhibits clear catalytic turnover as multiple monomers can be converted into waste for each template, with systems at low template concentrations producing output signals several times the initial template concentration (Fig. 3d and Supplementary Fig. S40-42). Proofreading thus returns a functional template as required.

The purpose of KP is to enrich the population of template-bound *M*_*1*_ monomers relative to template-bound *M*_*2*_ and *M*_*3*_. The concentrations of each template-bound monomer can be estimated by subtracting the waste concentration form the unblocked monomer concentration (subtracting Fig. 3c from Fig. 3b for *P*_*8*_ and *T*= 4 nM). Although this approach is crude, it provides clear evidence for a spike in in *M*_*1*_*T* at short times, with *M*_*2*_*T* and *M*_*3*_*T* highly suppressed at all time points (Fig. 3d-f and Supplementary Fig. S43). This behaviour is qualitatively consistent with predictions of an ODE model, parameterized by fits to earlier experiments (Supplementary Fig. S43). We suspect that the apparent ∼1 nM yield of *M*_*3*_*T* at long times is likely an artefact of the crude methodology. This interpretation is supported by the fact that the apparent yield of *M*_*3*_*T* is almost as high for *P*_*6*_ as *P*_*8*_, despite *P*_*6*_ having much higher affinity for *M*_*3*_.

### Templated dimer formation with kinetic proofreading

Having discriminated between the correct and incorrect monomers at both the template binding and template recovery steps, we redesigned the DNA strands to allow for dimerization through HMSD, using a handhold domain in a new template, *T’*. We also redesigned the monomers, now called *M’*_*1*_, *M’*_*2*_, and *M’*_*3*_, to accommodate three mismatches between the blocker and monomers (Fig.1c), rather than two. One of those mismatches is eliminated in template binding and remaining two during HMSD. In this redesign, we retained the domain lengths from the previous characterisation process but changed the sequences (Supplementary Table S15-17).

The handhold recognises the second monomer *N*, allowing dimerization, via HMSD (Fig. 1c). The intermediate *M’T’* can also be invaded by the redesigned proofreader *P’* to form waste *M’P’*, allowing kinetic proofreading of *M’*. We explored this process by adding various template concentrations to mixtures of monomers and proofreaders, with *M’*_*1*_, *M’*_*2*_ and *M’*_*3*_ probed separately. Dimer concentration was reported by an external reporter similar to that used for waste characterisation (Fig. 1d). We observed that the reporter had a small but significant tendency to react directly with the monomers (Supplementary Fig. S44). We therefore monitored lock strand removal by the increase in Cy3 signal intensity, and added external reporter only after the Cy3 signal reached a plateau for all template-containing systems.

We tested the three variants *P’*_*6*_, *P’*_*7*_ and *P’*_*8*_, with the results for *P’*_*6*_ and 2nM *T* shown in Fig. 4a-b, and the others in Supplementary Fig. S44-52. All systems showed discrimination at the first step, which is not proofreader-dependent; formation of *M’*_*1*_*T’* is substantially faster than *M’*_*2*_*T’* or *M’*_*3*_*T’*, as evidenced by the Cy3 signal. In the absence of proofreader, however, systems with *M’*_*1*_, *M’*_*2*_ or *M’*_*3*_ all reach high concentrations of *M’N* eventually, as evidenced by the dimer reporter signal. By contrast, in the presence of proofreaders *P’*_*6*_ or *P’*_*7*_, the eventual production of *M’*_*2*_*T’* and *M’*_*3*_ *T’* is strongly supressed; the level of incorrect product in these cases is comparable to the inherent leak of the reporters (Supplementary Fig. S44-52). Quantitatively, one may consider the discrimination factor *D*_x:y_, which in this case is given by the yield of *M’*_*x*_*T’* relative to *M’*_*y*_*T’* at the end of the experiments, since the total input concentration of each monomer is equal. In the absence of any proofreader, the discrimination factor is approximately one, because all monomers obtain essentially the same final yield. Specifically, for the data plotted in Fig. 4b, *D*_l:2_ = 0.87 and *D*_l:3_ = 0.92 with no proofreader present. In the presence of *P’*_*6*_ (Fig. 4a) we obtain *D*_l:2_ = 3.00 and *D*_l:3_ = 4.20, a 4-5 fold improvement. For the equivalent data for *P’*_*7*_, shown in Supplementary note 18, Supplementary Table S12, we obtain *D*_l:2_ = 2.71 and *D*_l:3_ = 3.91.

**Figure 4.**
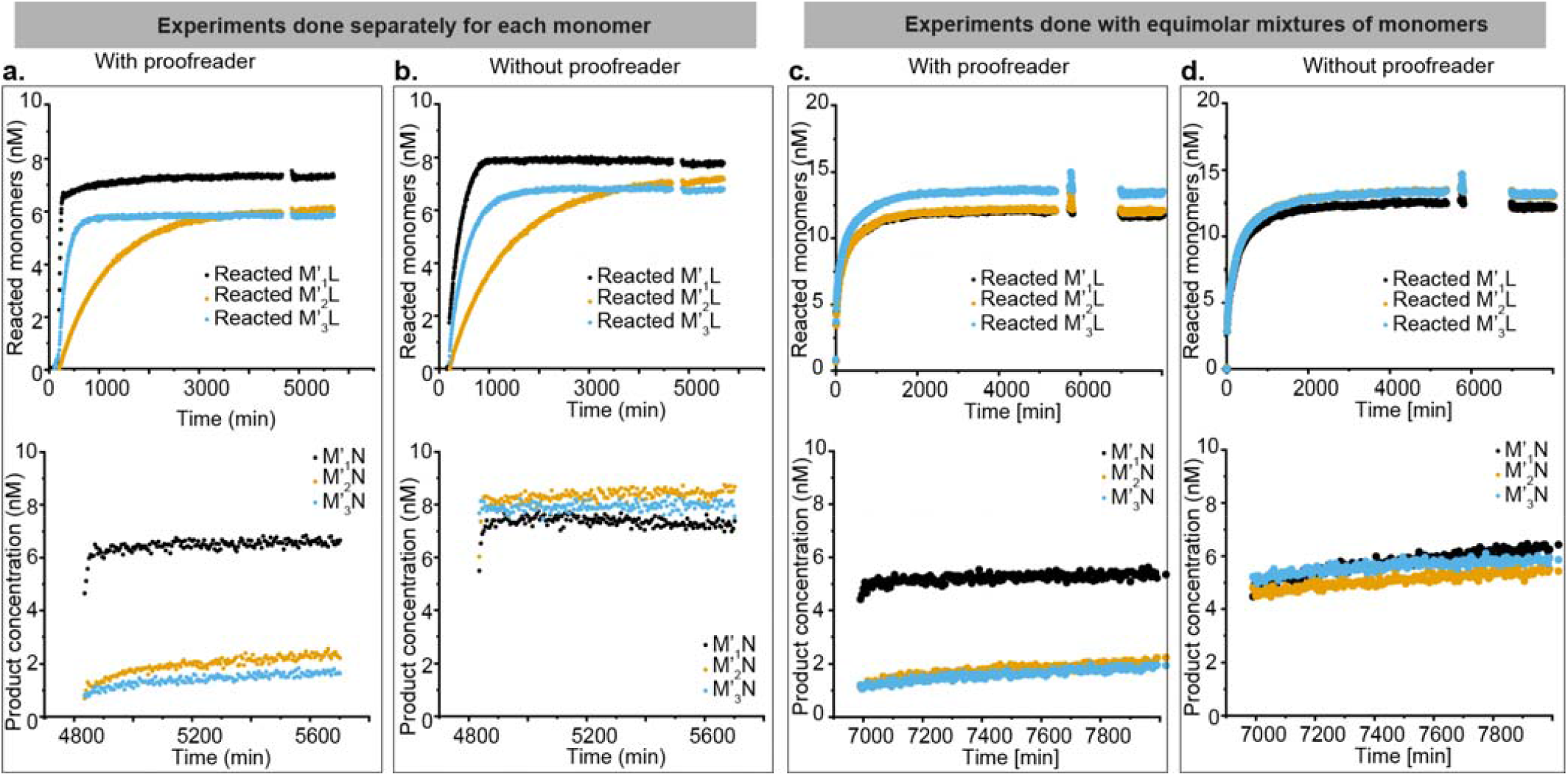
Kinetic proofreading in a dimerization process. (a) Top: Concentration of reacted monomers in a dimerization reaction with proofreaders. 8nM blocked monomers *M’*_*1*_, *M’*_*2*_ and *M’*_*3*_ in separate wells were mixed with 10 nM *N*, 50 nM *P’*_*6*_ and the reactions were triggered by injecting 2 nM *T’*. Unblocked monomer concentrations were inferred from Cy3 fluorescence. Bottom: Dimeric product formation by HMSD in the same reaction. After three days, the extent of product formation was assayed by adding 20 nM of the corresponding reporters; product concentration is inferred from reporter fluorescence, as outlined in Supplementary Note 15 and Fig. S44-49. We observe a much higher yield of *M’*_*1*_*N* than for the alternatives. (b) Top: Blocker strand removal from identical experiments in absence of the proofreader. Bottom: Dimeric product formation in reactions without a proofreader are almost equal for all monomers. (c) Top: Total concentration of monomers liberated in a competitive system initiated with 5nM each of *M’*_*1*_*L’, M’*_*2*_*L’*, and *M’*_*3*_*L’*, 15nM *N* and 50nM *P’*_*6*_ triggered with 2nM T’. Three replicas were analysed using the same approach as in (a), one to track each of *M’*_1_, *M’*_2_, and *M’*_3_ separately. Bottom: Product formation in these experiments. As in (a), we observe a much higher yield of *M’*_*1*_*N* than for the alternatives. (d) Top: Total concentration of monomers liberated in identical experiments to (c) but without *P’*_*6*_. Bottom: All products *M’*_*1*_*N, M’*_*2*_*N* or *M’*_*3*_*N* have a high yield.

This performance indicates successful proofreading, in which the alternative proofreading discard pathway competes effectively with incorporation into the dimerised state – although we note that the apparent mechanism for *P’*_*6*_ was a little unexpected (Supplementary Note 9). By contrast, *P’*_*8*_ does not show effective proofreading. We believe that, in this case, the proofreading pathway is too slow, allowing *M’*_*2*_*T’* and *M’*_*3*_*T’* to form even when *P’*_*8*_ is present. Model predictions based on the experiments in Fig. 2 (Supplementary Fig. S53) show high yields of *M’*_*2*_*T’* for *P’*_*7*_ and *P’*_*8*_; in the case of *P’*_*8*_, slower proofreading contributes to this response.

As with template recovery, the reservoir of proofreading molecules acts in a fundamentally different manner to an excess of blocker strands. In Supplementary Fig. S54, we show that an excess of blockers does not prevent the eventual formation of *M’*_*2*_*T’* and *M’*_*3*_*T’*, and only has a weak effect on kinetics. To demonstrate that our KP network enhances discrimination from a mixed pool of distinct monomers, we prepared an equimolar mixture of *M’*_*1*_*L, M’*_*2*_*L* and *M’*_*3*_*L* with an equal concentration of each duplex and an excess of *N*. As in the individual experiments, *P’*_*6*_ and *P’*_*7*_ again significantly suppressed the formation of incorrect dimers *M’*_*2*_*N* and *M’*_*3*_*N* while allowing *M’*_*1*_*N* to be formed at a high level (Fig. 4c and d, Supplementary Fig. S55-60).

### SNP identification via kinetic proofreading

We next demonstrate how KP can be incorporated into a simple functional network designed to detect SNPs in a DNA strand where the mutations are anywhere in the displacement domains (Fig. 5a). It is imperative to state that we do not propose this particular network as a competitive alternative method to the existing approaches in isolation. Our emphasis is not on the absolute performance of the system before or after proofreading, but exploring the relative improvement in specificity of a reaction network due to the inclusion of KP. Several DNA strand displacement-based detection methods involving TMSD or toehold exchange have been proposed to differentiate between the wild type and the SNPs^15,35–38^. Such systems could potentially benefit from adding some form of KP to the detection process.

**Figure 5.**
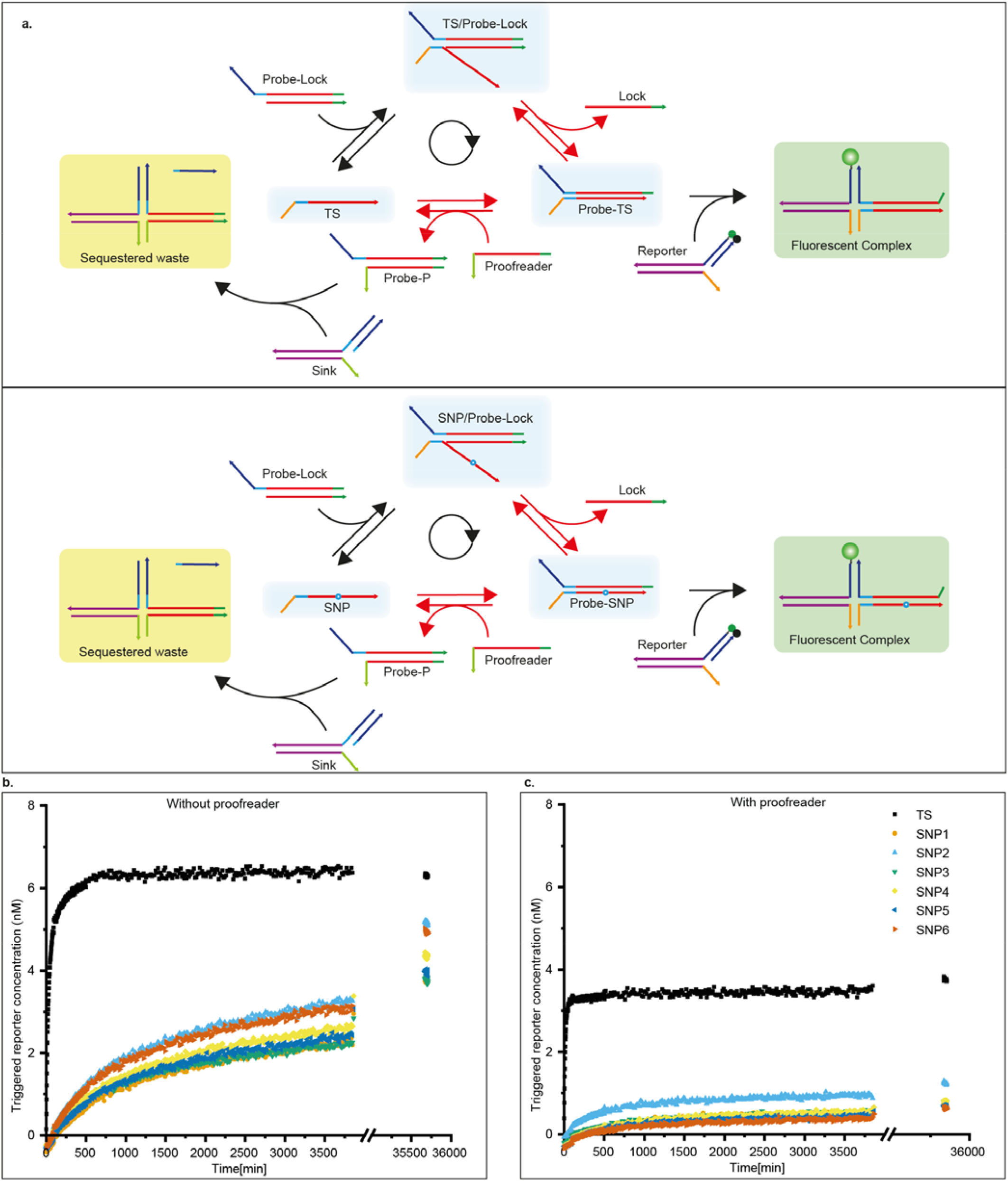
SNP identification scheme utilizing KP to suppress output of mismatched target. (a) Reaction scheme: *TS* and *SNP* strands bind to the blocked probe to form the intermediate *TS-Probe* or *SNP-Probe*, which trigger the fluorescence by displacing the quencher strands from the external reporters. Proofreader molecules can also invade these intermediates to convert the *Probe* into *Waste*, which is in turn sequestered into an inert complex by the *Sink*. The red-coloured arrows denote two stages of discrimination between *TS* and *SNP*. (b) Fluorescence signal triggered by the *TS* and 6 different *SNPs* when 10 nM of *Probe-Lock* is added to a mixture containing 15 nM of candidate strands, 20 nM of corresponding reporters, and 20 nM of *Sink* complex. (c) The same reactions are performed in presence of 20 nM Proofreader *P*_*snp*_. While the *TS* still shows rapid growth up to about 4 nM in the reported strand concentration, the *SNPs* show limited increase the signal even after several days. At least some of the residual activation apparent in (Fig. 5c) is due to a leak reaction between *Probe-lock* and reporter (Supplementary Fig. S61).

In the context of SNP detection, it is natural for the strand being tested – either the Target Strand (*TS*) or SNP-bearing strand (*SNP*) – to be single stranded. We therefore consider a different reaction network structure relative to the dimerization example, whilst maintaining the central KP functionality. By embedding KP within another functional network, we demonstrate its versatility.

A *Probe* strand is fully complementary to *TS* whereas the *SNP*s have a mismatch with the *Probe* at the point of mutation. The candidates (*TS* or *SNP*s) bind to the initially blocked *Probe* via TMSD, providing initial discrimination. This TMSD reveals a second toehold that facilitates the slow formation of a stable complex with fluorescent reporters. Alternatively, proofreader molecules *P*_snp_ can react with intermediates via the second toehold to form a *Probe-P*_snp_ complex that is sequestered by a sink *S*. This proofreading discard pathway provides a second opportunity to discriminate between *TS* and *SNP*. We generated a random *TS* conforming to our design and introduced a single mismatch at different positions of the displacement domain to give six *SNP* strands (Supplementary Table S18). We used different nucleotide mismatches to vary the free energy of the mismatched pair, and varied the location across the branch migration domain because the effect of mismatches on strand displacement is known to be strongly location-dependent. By design, the same proofreader and probe strands can be used regardless of the specific competing *SNP*. We triggered the system by adding a blocked *Probe* to a solution containing the candidate strand, reporter complex and sink complex (Supplementary Notes 8, 16). Fluorescent traces with and without 20nM proofreader are shown in Fig. 5; results for other concentrations of proofreader are given in Supplementary Fig. S61-62.

The proofreader improves accuracy in two ways. Firstly, by selectively reacting with the intermediate, the initial rate of product formation for *SNP*s is reduced by an additional factor of 2-3 relative to *TS* when proofreaders are used (Supplementary Fig. S63-64, Table S18). Secondly, the existence of a fuel-consuming discard pathway allows *SNP*-triggered probes to be consumed (Fig. 5c), preventing them from eventually reacting with the reporter, as is typical in the absence of a proofreader strand (Fig. 5b). We may once again calculate a discrimination factor *D*_TS:SNPx_ at the end of the experiment for each proofreader concentration (Supplementary Note 19, Supplementary Table S13). In the absence of proofreader, typical values of *D*_TS:SNPx_ ≈ 1 - 1.7 are obtained, relative to *D*_TS:SNPx_ ≈ 3- 7 for 20nM of proofreader.

## Conclusion

We have demonstrated that kinetic proofreading cycles, a ubiquitous biological motif, can be incorporated non-enzymatic DNA strand displacement networks. To the best of our knowledge, this work represents the first time a kinetic proofreading system has been engineered *de novo*. This successful demonstration of KP in a synthetic context sets the groundwork for applying the motif more broadly and opens pathways to explore the fundamental principles of KP in engineerable systems.

We have shown how KP can enhance the discrimination between perfectly-matching DNA strands and single nucleotide mutants, both in toeholds and branch migration domains. In this context, the relatively small difference between targets is unavoidable, making it challenging to redesign a system to increase the discrimination at equilibrium between the correct strand and many distinct variants simultaneously. Reexploiting a single free-energy difference through KP is therefore especially valuable. We note that even if discrimination exists without KP, enhancing that discrimination will always improve detection of low concentrations of a target from within a pool of competitors. KP relies on pushing the template through fuel-driven non-equilibrium cycles. In our design, the large excess of proofreader tends to drive the template through the cycle, although backwards steps that reverse the proofreading reaction are not impossible. The presence of downstream reactions, such as the sequestration of waste by reporter molecules, can provide an additional contribution to the thermodynamic drive. Although such a design feature may enhance KP, it is not required for our system – as evidenced by the successful KP in the dimeric production experiments.

Sensitivity to downstream reactions can be a significant factor in responsive circuits. One established approach to effectively discriminate between SNPs is to use toehold exchange probes that are carefully designed so as to render the overall freeenergy change of probe-target binding close to zero^15^. This method ensures that a destabilizing mismatch will have a substantial effect on the yield of product in steady state. These motifs work well in isolation, but can struggle to give large discrimination factors in steady state when incorporated into a larger circuit when the downstream reactions will tend to perturb the thermodynamics, as mentioned above. The KP motifs considered here have successfully boosted specificity even when coupled to downstream reactions that bias the thermodynamics, and are thus potentially well suited to incorporation into larger functional networks.

In our designs, the free energy used to drive the recognition site through a proofreading cycle comes directly from converting the inputs into waste, rather than coupling to an ancillary fuel molecule as imagined by Hopfield. As a result, rejected monomers are sequestered, rather than being released to attempt incorporation again. In the context of a one-off detection process, as considered here, this absorption of monomers is potentially beneficial. The discard pathway provided by KP allows the incorrect species to be rendered inert, so that incorrect products are never made at all, rather than just delayed as in a scheme powered by ancillary fuel molecules. This suppression of an unwanted signal even on long time scales, as much as the rate advantage provided by KP, will likely aid in the design of more precise synthetic molecular networks. This behaviour is distinct from thresholding^29^, in which a rapid reaction with a thresholding molecule is used to consume inputs so that output signals are only produced when the input concentrations exceed the concentration of the threshold molecule. Mechanistically, KP does not rely on depletion of the proofreader to propagate a signal, and so doesn’t require a minimal input concentration to generate an output. Moreover, thresholding molecules will, if anything, tend to sequester correct species faster than their mutant counterparts.

## Supporting information

Experimental protocols and supporting data

## ASSOCIATED CONTENT SUPPORTING INFORMATION

All the fluorescence data as obtained from the plate reader, the Mathematica scripts used to process those data, convert to concentration, and fit them to the ODE models, and the figures are available at https://doi.org/10.5281/zenodo.8132461.

The Supporting Information is available free of charge on the ACS Publications website.

## Notes

A preprint version of this work is available at bioRxiv: Kinetic Proofreading can Enhance Single Nucleotide Discrimination in a Non-enzymatic DNA Strand Displacement Network, Rakesh Mukherjee, Aditya Sengar, Javier Cabello-García, Thomas E. Ouldridge; doi: https://doi.org/10.1101/2023.08.25.554917

## ACKNOWLEDGMENT

The authors are grateful to Dr Wooli Bae and Dr Francesca Smith for their helpful discussions.

---

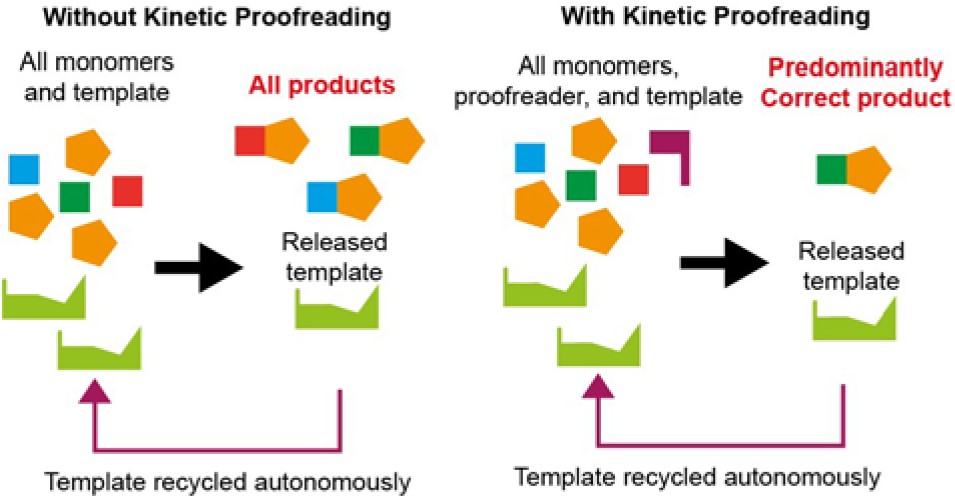

