## Supplementary material for "Kinetic Proofreading can Enhance Specificity in a Non-enzymatic DNA Strand Displacement Network": Experimental protocols and supporting data

### Table of Figures

Figure S1: Design details of the monomer toeholds.	8

Figure S2: Design strategy and reporting mechanism for the external reporters.	9

Figure S3: Example data of T binding to blocked monomers	12

Figure S4: Example data of L binding to M_2_T complex	14

Figure S5: Example data for measuring reporter rate for M_1_P_6_ complex	16

Figure S6: Example data for characterisation of template recovery from M_2_T complex by P_7_ in Cy3 channel	18

Figure S7: Example data for characterisation of template recovery from M_2_T complex by P_7_ in AlexaFluor-647 channel	19

Figure S8: Example data for characterisation of the discard pathway of the M_2_T complex tiggered by P_7_ in AlexaFluor-647 channel	21

Figure S9: Example data for characterisation of the discard pathway of the M_2_T complex triggered by P_6_ in Cy3 channel	22

Figure S10: Example data in the AlexaFluor-647 channel for product formation in a dimerisation reaction with kinetic proofreading	24

Figure S11: Example data in the Cy3 channel for the dimerisation process with kinetic proofreading	25

Figure S12: Example data of product formation in a dimerisation reaction with proofreading from an equimolar mixture of three different monomers monitored by AlexaFluor-647 fluorescence	27

Figure S13: Example data in the Cy3 channel for the dimerisation process with proofreading from an equimolar mixture of three different monomers	28

Figure S14: Example data of Cy3 signal from SNP detection scheme	30

Figure S15: Fluorescent signal from the Cy3 channel obtained for the template binding reaction of M_1_L for varying [T_total_].	32

Figure S16: Concentration of unlocked template M_1_T as a function of time.	33

Figure S17: Fluorescent signal from the Cy3 channel (a) and the Alexa channel (b) obtained for the reporter reaction of M_1_P_8_	35

Figure S18: Processed fluorescent signal from the Cy3 channel (a) and the Alexa channel (b) obtained for the reporter reaction of M_1_P_8_	37

Figure S19: Total monomer concentration from the Cy3 channel (a) and [MPR(t)] time signal from the Alexa channel (b) obtained for the reporter reaction of M_1_P_8_	39

Figure S20: Fluorescent signal from the Cy3 channel (a) and the Alexa channel (b) obtained for the template recovery reaction for monomer M_1_ and proofreader P_8_	41

Figure S21: Normalized fluorescent signal from the Cy3 channel (a) and the Alexa channel (b) obtained for the template recovery of monomer M_1_ by proof-reader P_8_	43

Figure S22: Total monomer concentration from the Cy3 channel (a) and [MPR]+[PR] time signal from the Alexa channel (b) obtained for the reporter reaction of M_1_P_8_	45

Figure S23: Fluorescent signal from the Cy3 channel (a) and the Alexa channel (b) for the template recovery reaction, where the monomer is M_1_ and proofreader is P_8_.	47

Figure S24: Normalized fluorescent signal from the Cy3 channel (a) and the Alexa channel (b) obtained for the discard pathway reaction with monomer M_1_ and proofreader P_8_	48

Figure S25: Normalized fluorescent signal from the Cy3 channel over a long time for the discard pathway reaction with monomer M_1_ and proof-reader P_8_	49

Figure S 26: [MPR]+[PR] time signal from the Alexa channel obtained for the discard pathway reaction with monomer M1 and Proofreader P8	50

Figure S27: Cy3 calibration curve for complexes containing M_1_	55

Figure S28: AlexaFluor-647 calibration curves for quenched and fluorescent reporters for M_1_P_6_	56

Figure S29: Cy3 calibration curves for complexes containing M_1_’	57

Figure S30: Calibration curves for quenched and fluorescent reporters for M_1_’N	58

Figure S31: Cy3 calibration curves for quenched and fluorescent reporter complexes for SNP detection	59

Figure S32: Template binding to ML complexes	60

Figure S33: L binding to MT complexes	62

Figure S34: Reporter characterisation for M_1_P complexes	64

Figure S35: Reporter characterisation for M2P complexes	66

Figure S36: Reporter characterisation for M_3_P complexes	68

Figure S37: Template displacement by the proofreaders from M_1_T complex	69

Figure S38: Template displacement by the proofreaders from M_2_T complex	71

Figure S39: Template displacement by the proofreaders from M_3_T complex	74

Figure S40: Conversion of blocked monomer M_1_L into waste M_1_P via template T	75

Figure S41: Conversion of blocked monomer M_2_L into waste M_2_P	77

Figure S42: Conversion of blocked monomer M_3_L into waste M_3_P	79

Figure S43: Experimental estimation and model prediction of MT concentrations in the discard pathway experiments	82

Figure S44: M’_1_N dimer formation in presence (top row) and absence (bottom row) of P’_6_.	84

Figure S45: M’_2_N dimer formation in presence (top row) and absence (bottom row) of P’_6_.	85

Figure S46: M’_3_N dimer formation in presence (top row) and absence (bottom row) of P’_6_.	86

Figure S47: M’_1_N dimer formation in presence (top row) and absence (bottom row) of P’_7_.	87

Figure S48: M’_2_N dimer formation in presence (top row) and absence (bottom row) of P’_7_	88

Figure S49: M’_3_N dimer formation in presence (top row) and absence (bottom row) of P’_7_.	89

Figure S 50: M’_1_N dimer formation in presence (top row) and absence (bottom row) of P’_8_.	90

Figure S 51: M’_2_N dimer formation in presence (top row) and absence (bottom row) of P’_8_.	91

Figure S 52: M’_3_N dimer formation in presence (top row) and absence (bottom row) of P’_8_.	92

Figure S53: Prediction of dimerization rates with and without different proofreaders	93

Figure S54: M’_3_N dimer formation in presence (left) and absence (right) of excess blocker	94

Figure S55: M’_1_N dimer formation from a mixture of M’ monomers in presence (top row) and absence (bottom row) of P’_6_	96

Figure S56: M’_2_N dimer formation from a mixture of M’ monomers in presence (top row) and absence (bottom row) of P’_6_	97

Figure S57: M’_3_N dimer formation in presence (top row) and absence of P’_6_ (bottom row) from a mixture of monomers	98

Figure S58: M’_1_N dimer formation from a mixture of M’ monomers in presence (top row) and absence (bottom row) of P’_7_	99

Figure S59: M’_2_N dimer formation from a mixture of M’ monomers in presence (top row) and absence (bottom row) of P’_7_	100

Figure S60: M’_3_N dimer formation from a mixture of M’ monomers in presence (top row) and absence (bottom row) of P’_7_	101

Figure S61: Concentration of candidate-probe complexes with different concentrations of proofreader.	102

Figure S62: Reporter characterisation for TS-Probe and SNP-Probe complexes	106

Figure S63: Estimates of initial reaction rates in SNP detection system with and without 20 nM proofreader.	107

Figure S 64: Comparison of TS and control SNP reactions with same toehold	109

### Table of tables

### Supplementary Figure 1: Monomer design

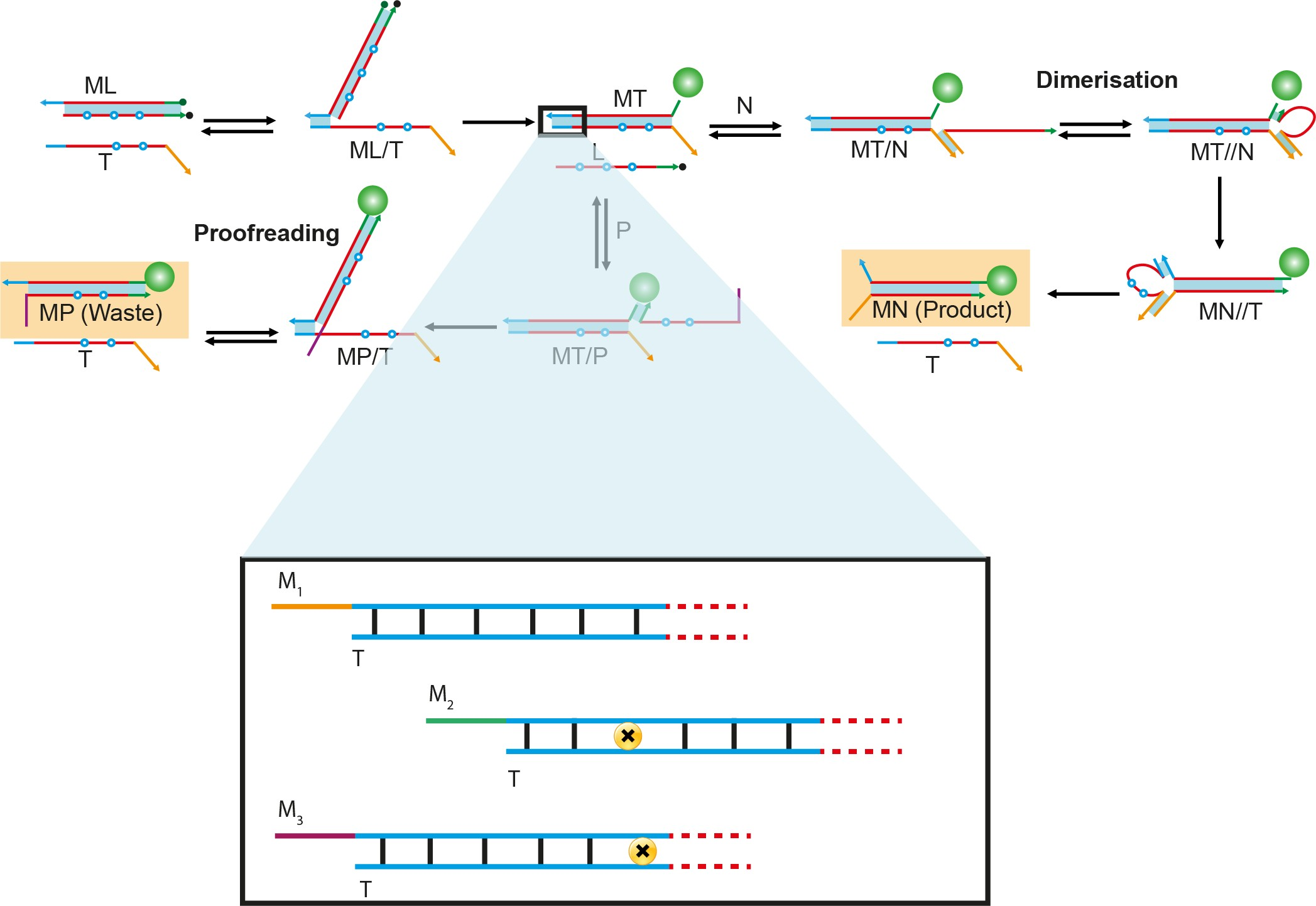

Figure S1: Design details of the monomer toeholds.

Monomer *M_1_* has a fully complementary toehold (blue domains) with the template. *M*_2_ and *T* have a Monomer *M_1_* and *M’_1_* has a fully complementary toehold (blue domains) with the template. *M*_2_ or *M’_2_* and *T* have a mismatch (yellow circle with a black cross) in the middle of the toehold. *M_3_* or *M’_3_* and *T* have a mismatch at the end of the toehold bofore the displacement domain (dashed red domains). Overhanging domains on the monomers (yellow for *M_1_* and *M’_1_*, green for *M_2_* and *M’_2_*, and purple for *M_3_* and *M’_3_*) are additional 2 nucleotide domains for reporter binding.

### Supplementary Figure 2: External reporter design

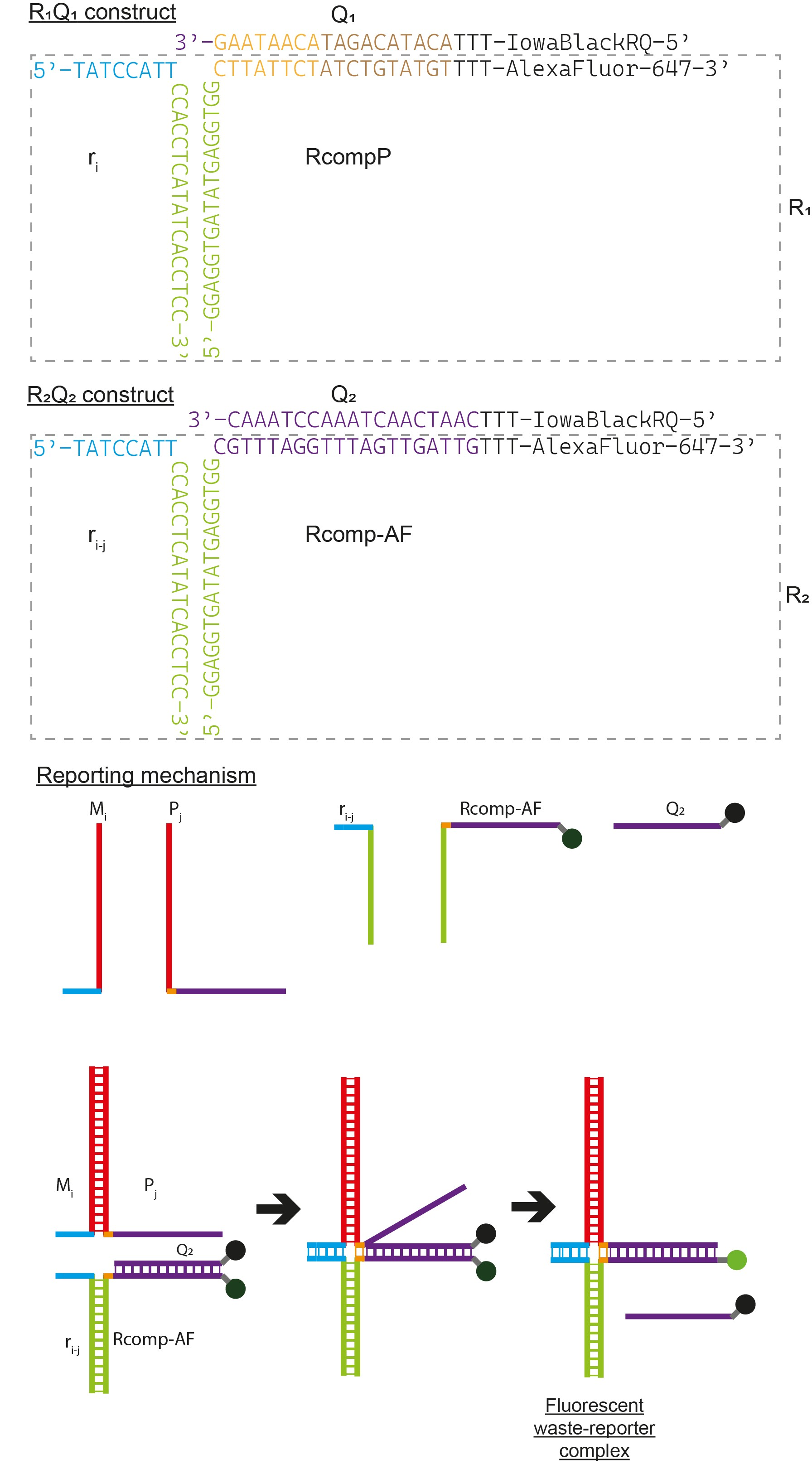

Figure S2: Design strategy and reporting mechanism for the external reporters.

The external reporters are used to monitor the formation of *MP* and *M’N* complexes. This method reduces the number of required fluorophore-labelled oligonucleotides, thereby reducing the cost and the complications of the experiments. The reporter-quencher complex is formed of three strands. *r_i-j_* (for *M_i_P_j_* waste complexes) or *r_i_* (for *M’_i_N* dimers) and *Rcomp-AF* (for *M_i_P_j_* waste complexes) *or RcompP* (for *M’_i_N* products) have the long complementary domain (green) to hold them together. *Rcomp-AF*/*RcompP* and *Q_1_/Q_2_* have another complementary domain (purple) at the end of which the two strands are labelled with a fluorophore (AlexaFluor-647) and a quencher (IowaBlack-RQ) respectively. Top two Figure show two examples of the reporter-quencher complexes along with their sequences. The three-strand reporter-quencher complex has the blue domains on *r_i-j_* and yellow domain on *Rcomp-AF or RcompP* to serve as a toehold domain for the *M_i_P_j_* or *M’_i_N* complexes to displace the *Q* strands and trigger fluorescence. Neither *M* nor *P* strands can complete this displacement efficiently unless bound into a duplex. For different monomers, one only needs to alter the unlabelled *r_i-j_* strand to form another reporter-quencher complex specific for that monomer-proofreader combination. Same strategies were employed for the reporter in SNP detection, and the sequestering sink complex. All the sequences are listed in Tables S14-16.

### Supplementary note 1: Protocol for template binding to blocked monomers

This protocol was used to gather data reported in Figure 2b of main text and Figure S32 of Supplementary information.

**Step 1:** 130 µL of experimental buffer was added to each well. The fluorescence values were measured over at least 10 data points.

**Step 2:** 20 µL of pre-annealed *M_1_L* complex (100 nM stock) was added to the buffer to get a final concentration (after step 3) of 10 nM. Fluorescence intensities were measured again over at least 10 data points to determine residual fluorescence of the quenched complexes.

**Step 3:** Different volumes of template strand (50 nM stock) and the experimental buffer were injected into the wells via the injector of the plate reader to make the final volume in each well 200 µL. The total injected volume of template and buffer was 50 µL for each well. Template injection volumes were varied to attain a final *T* concentration of 0-10 nM in the wells. The initial kinetics of template binding were followed by monitoring the change in fluorescence intensities for several minutes to several hours depending on the reaction rates.

Table S1: Experimental method for T binding

| **Steps** | **Components added** | **Purpose** | **Target concentration in 200 µL** | **Volume in µL** | **Measurement no.** |
| --- | --- | --- | --- | --- | --- |
| Step 1 | 130 µL buffer | Baseline fluorescence | n/a | 134 | - |
| Step 2 | 20 µL *M_1_L* (100 nM) | Residual fluorescence | 10 nM of *M_1_L* | 150 | 1 |
| Step 3 | 0-40 µL *T*, 50-10 µL buffer | Triggering the experiment | 0-10 nM of *T* | 200 | 2 |

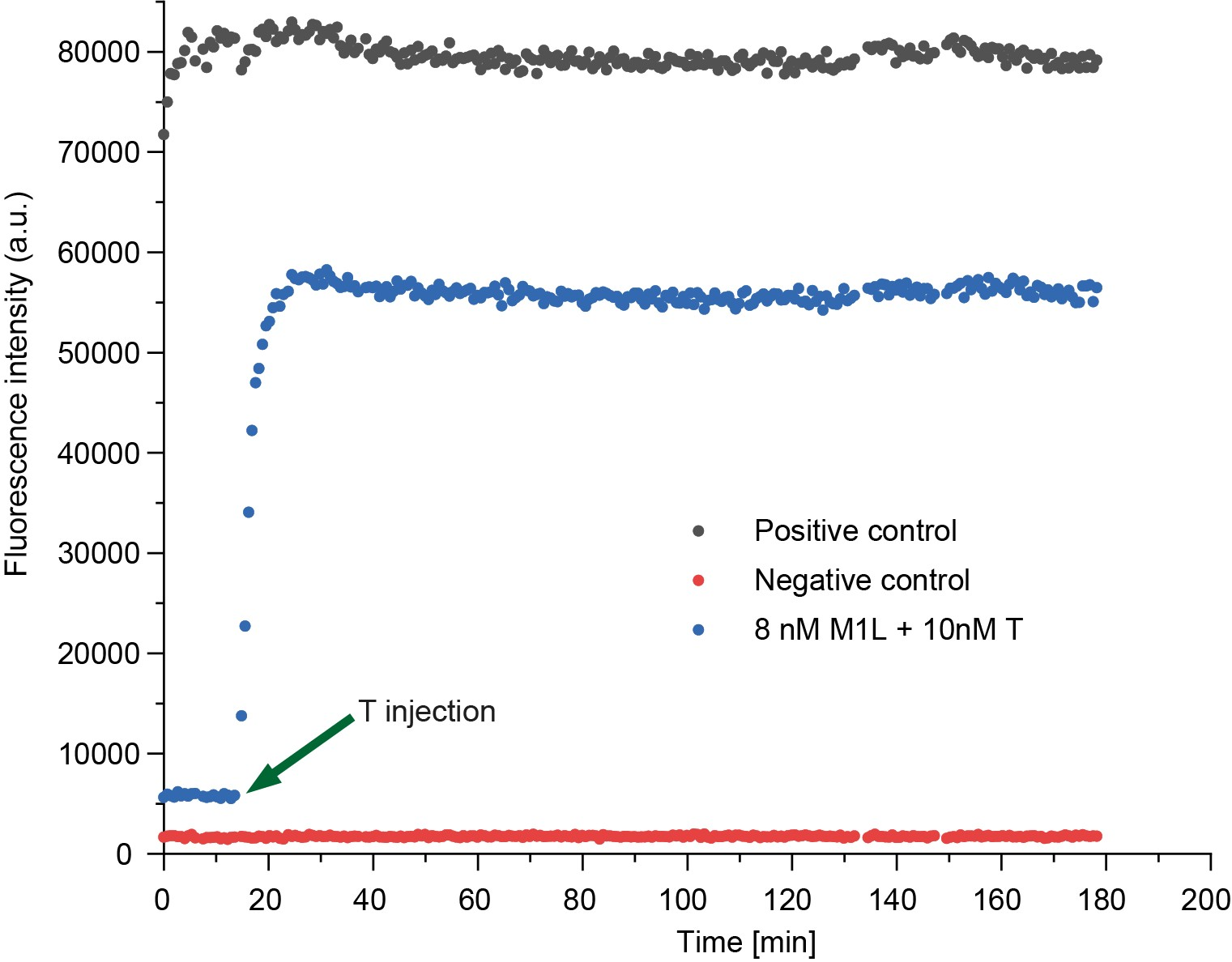

Figure S3: Example data of T binding to blocked monomers

For measurement 1, 10 nM of *M_1_L* complex in buffer was put into a plate well and fluorescence was measured. Then 0-10 nM of *T* was added to the well and the kinetics of the reaction was followed by monitoring Cy3 fluorescence (measurement 2). Only buffer was used as a negative control and pre-annealed *M_1_T* complex was used as positive control.

### Supplementary note 2: Protocol for *L* binding back to *MT* complexes

**Step 1:** 130 µL of experimental buffer was added to each well. The fluorescence values were measured over at least 10 data points.

**Step 2:** 20 µL of pre-annealed *M_1_T* complex (100 nM stock) were added to the buffer to get a final concentration (after step 3) of 10 nM. Fluorescence intensities were measured again over at least 10 data points to determine residual fluorescence of the quenched complexes.

**Step 3:** Different volumes of *L* strand (50 nM stock) and the experimental buffer were injected into the wells via the injector of the plate reader to make the final volume in each well 200 µL. The total injected volume of template and buffer was 50 µL for each well. Template injection volumes were varied to attain a final *L* concentration of 0-100 nM in the wells. The initial kinetics of template binding were followed by monitoring the change in fluorescence intensities for several minutes to several hours depending on the reaction rates.

Table S2: Experimental method for L binding back to MT complexes

| **Steps** | **Components added** | **Purpose** | **Target concentration in 200µL** | **Volume in µL** | **Measurement no.** |
| --- | --- | --- | --- | --- | --- |
| Step 1 | 130 µL buffer | Baseline fluorescence | n/a | 134 | - |
| Step 2 | 20 µL *M_1_T* (100 nM) | Residual fluorescence | 10 nM of *M_1_T* | 150 | 1 |
| Step 3 | 0-40 µL *L*, 50-10 µL buffer | Triggering the experiment | 0-100 nM of *L* | 200 | 2 |

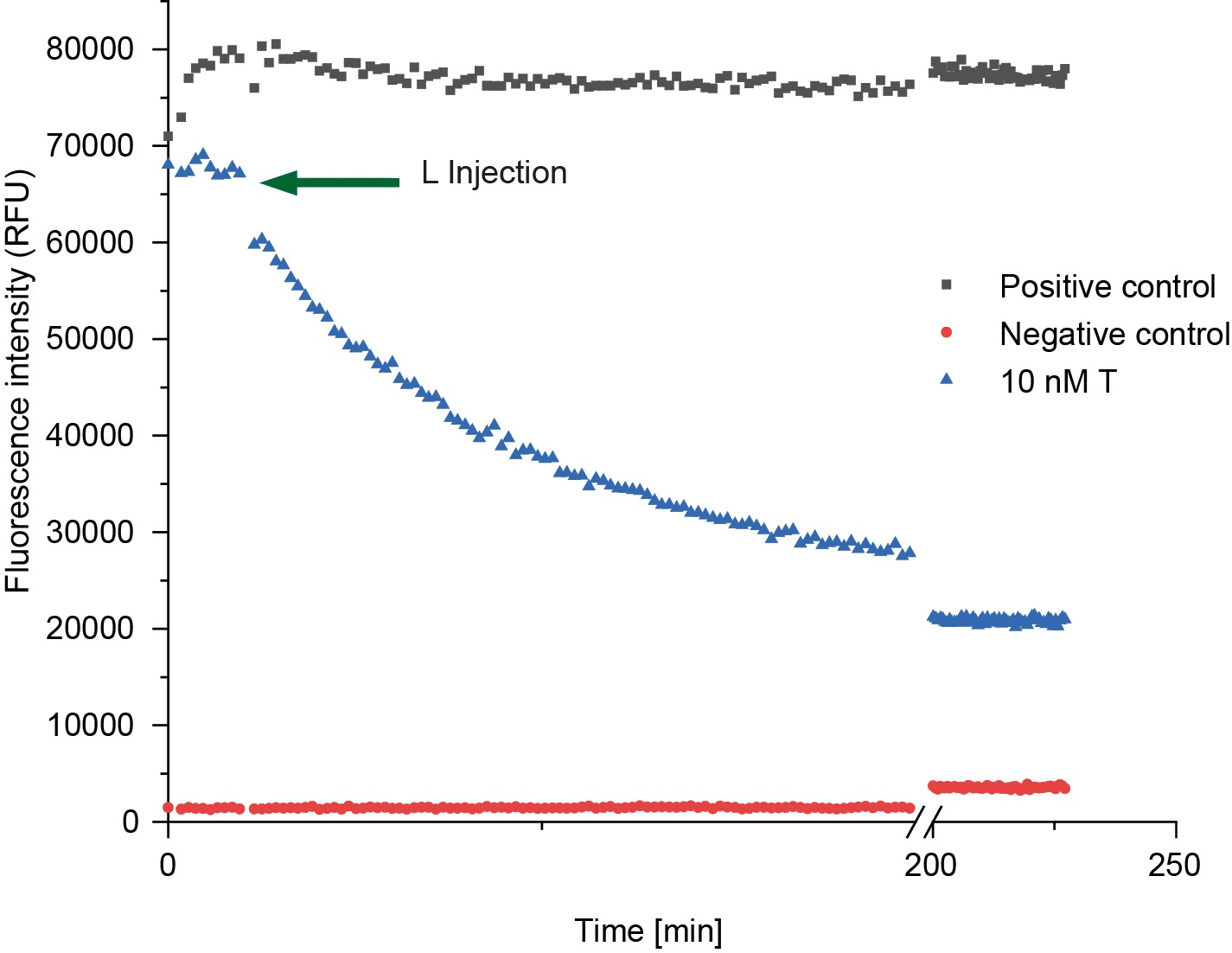

Figure S4: Example data of L binding to M_2_T complex

For measurement 1, 10 nM of *M_2_T* complex in buffer was put into a plate well and fluorescence was measured. After this, 0-100 nM of *L* was added to the well and the kinetics of the reaction was followed by monitoring Cy3 fluorescence (measurement 2). Pre-annealed *M_2_T* complex was used as a positive control.

### Supplementary note 3: Protocol for reporter characterisation for *MP* complexes

This protocol has been used to gather data shown in Figure S34-36 in the Supplementary information.

Given the fast intended rates of reporter reactions, these sets of experiments were done in ‘well mode’ kinetics. In this mode of measurement, a single well of the 96-well plate is scanned for the user-specified time, unlike the plate mode in which the entire plate or the all the specified wells are scanned together.

The experiments were performed in the following steps:

**Step 1:** 140-150 µL of buffer was added to each well to accommodate the volumes for different concentrations of other components.

**Step 2**: 0-10 µL of *MP* (200 nM stock) were added to different wells to give variable concentrations.

**Step 3:** 40 µL of the corresponding reporters and 10 µL of buffer were injected to each well via the injector to initiate the reactions. Final volumes of each well were 200 µL. Reaction kinetics measurements were started from here.

A negative control was obtained by only measuring 200 µL of buffer to account for the baseline fluorescence. A positive control was obtained by measuring the fluorescence of 5 nM of corresponding *MP* complex in buffer with a total reaction volume of 200 µL. The residual fluorescence values of the reporters were obtained from the well containing 0 nM *MP* complexes, essentially consisting of buffer and 40 nM reporter complex only. The actual amount of *MP* complex added were monitored by Cy3 fluorescence channel. Reporter complexes were measured in AlexaFluor-647 channel.

Table S3: Experiemntal method for reporter characterisation

| **Step** | **Components added** | **Purpose** | **Target concentration in 200µL** | **Volume in µL** | **Measurement no.** |
| --- | --- | --- | --- | --- | --- |
| Step 1 | 140-150 µL buffer | Dilution and volume adjustment | n/a | 140-150 | - |
| Step 2 | 0-10 µL *M_1_P* | Initial concentration of *M_1_P* | 0-10 nM of *M_1_P* | 150 | 1 |
| Step 3 | 40 µL reporter, 10 µL buffer | Tiggering the reaction | 40 nM reporter | 200 | 2 |

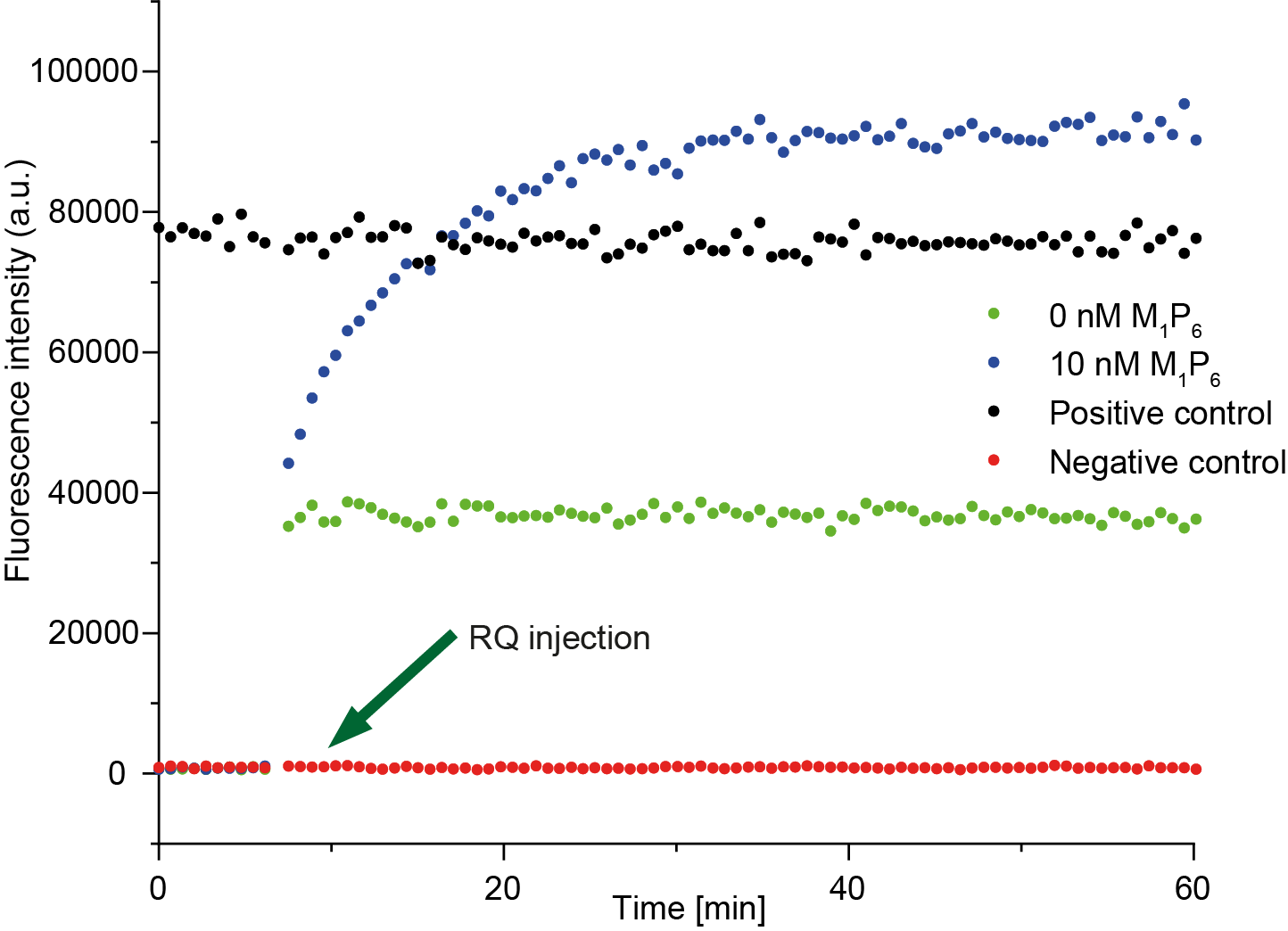

Figure S5: Example data for measuring reporter rate for M_1_P_6_ complex

For measurement 1, 0-10 nM of *M_1_P_6_* in buffer was put into a plate well and fluorescence was measured. After 10 scans, 40 nM of *RQ* complex was added to the well and the kinetics of the reaction was followed by monitoring AlexaFluor-647 fluorescence (measurement 2). Only buffer was used as a negative control and pre-annealed *M_1_P_6_R* complex was used as positive control. 40 nM *RQ* in buffer was used to estimate the residual fluorescence.

### Supplementary note 4: Protocol for template recovery

This method was used to gather experimental data shown in Figure 2d in the main text and Figure S37-39 in the Supplementary information.

**Step 1:** 114-130 µL buffer was added to each well and the fluorescence measurement was taken for 10 timepoints to determine the baseline fluorescence for both Cy3 and AlexaFluor-647 channels.

**Step 2:** 0-16 µL *MT* (100 nM stock) was added to each well to achieve a target range of concentration from 0 nM to 8 nM. The fluorescence measurements were taken to estimate the actual concentrations of the added *MT* complexes.

**Step 3:** 20 µL *RQ* (200 nM stock) was added to each well to obtain a target reporter concentration of 20 nM. 10 fluorescent measurements were taken to estimate the residual fluorescence of RQ complexes.

**Step 4:** 10 µL *P* (1 µM stock) and 40 µL buffer was injected to each well to trigger the reactions by obtaining a target *P* concentration of 50 nM. The reaction kinetics were followed by monitoring the changes in the fluorescence intensities of the reaction wells for several hours.

**Step 5:** After a few days, the plates were measured again to check the levels of Cy3 signal to ensure that there was no significant evaporation from the wells.

**Step 6:** 4 µL of the corresponding *M* (2 µM stock) was added to the wells to saturate the reporters to estimate the initial reporter concentrations.

Table S4: Experimental method for template recovery

| **Steps** | **Component added** | **Purpose** | **Target concentration in 200 µL** | **Volume in µL** | **Measurement no.** |
| --- | --- | --- | --- | --- | --- |
| Step 1 | 114-130 µL buffer | Baseline fluorescence | n/a | 114-130 | 1 |
| Step 2 | 0-16 µL *MT* (100 nM stock) | *MT* concentration | 0-8 nM *MT* | 130 | 2 |
| Step 3 | 20 µL reporter (200 nM stock) | Residual fluorescence from reporters | 20 nM reporter | 150 | 3 |
| Step 4 | 10 µL *P* (1 uM stock), 40 µL buffer | Trigger reaction | 50 nM *P* | 200 | 4 |
| Step 5 | n/a | Check anomalies |  | 200 | 5 |
| Step 6 | 4 µL *M* (2 uM stock) | Saturate the reporters | > 40 nM | 204 | 6 |

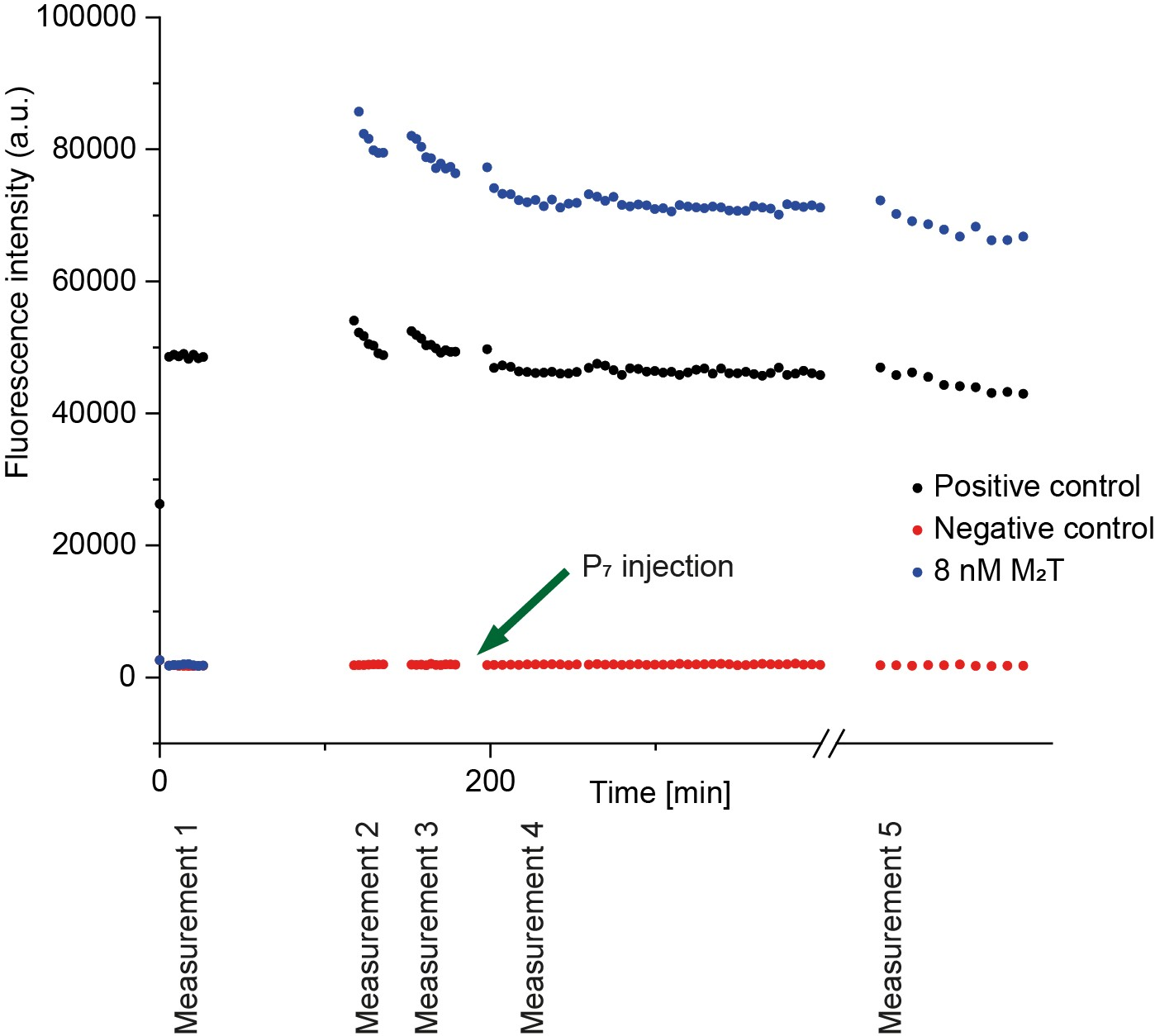

Figure S6: Example data for characterisation of template recovery from M_2_T complex by P_7_ in Cy3 channel

Cy3 fluorescence signal shows the amount of *M_1_T* complex added. Measurement 1 shows the baseline from the plate and buffer. Measurement 2 measures the change in fluorescence by adding 8 nM *M_2_T* complex. This measurement also shows the exact concentration of *M_2_T* added, which varies from the intended concentration due to minor dilution errors and experimental variations. Measurement 3 measures the residual fluorescence from the 20 nM *RQ* complex. Then the reaction is triggered by adding 50 nM *P_7_* and measurement 4 was performed for several hours. Measurement 5 was performed after several days to ensure that the reaction was complete, and to also ensure that there is minimal evaporation, if any. The decay in the signal of Cy3 at the beginning of each measurement is characteristic of temperature increase in the experiment. These signature decays are present in almost all the experiments at its initial phase where the 96-well plate is transferred from the ambient lab temperature of 22-23°C to the plate reader kept at 25°C.

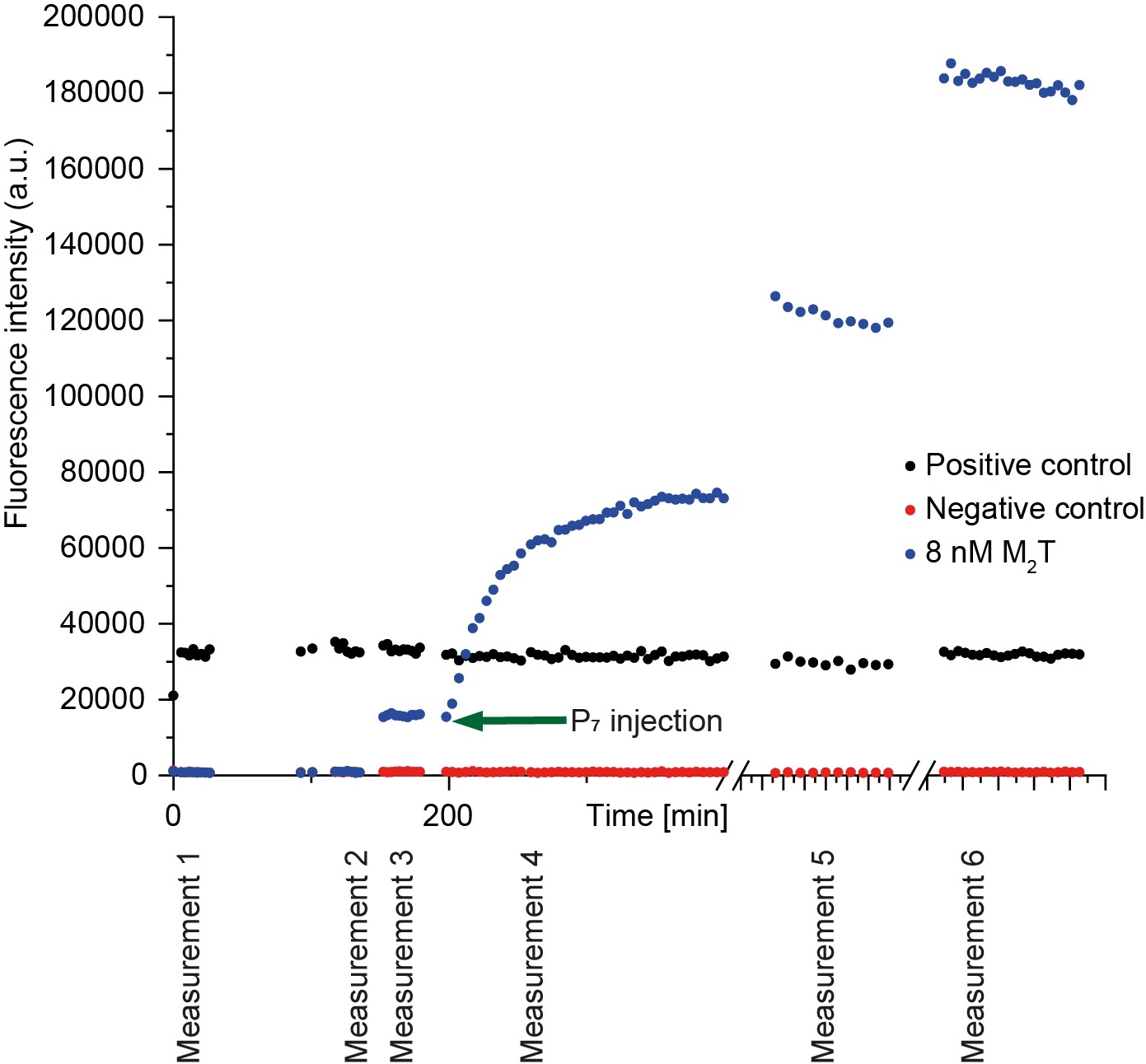

Figure S7: Example data for characterisation of template recovery from M_2_T complex by P_7_ in AlexaFluor-647 channel

AlexaFluor-647 fluorescence signal shows the formation of the *M_2_PR* complex triggered by the addition of 50 nM of *P_7_* into a solution containing 20 nM *RQ* and 8 nM *M_2_T*. Measurement 1 shows the baseline from the plate and buffer. Measurement 2 measures the change in fluorescence by adding 8 nM *M_2_T* complex. Measurement 3 gives the residual fluorescence from the 20 nM *RQ* complex. Then the reaction is triggered by adding 50 nM *P_7_* and measurement 4 occurs for several hours. Measurement 5 was performed after several days to ensure that the reaction is completed. For measurement 6, the reporter was saturated by the addition of another 40 nM of free *M_2_* which reacts with excess proofreader in the solution to form *M_2_P*, which then triggers the remaining reporter to estimate the initial concentration of *RQ*.

### Supplementary note 5: Protocol for discard pathway assays

This protocol was used for gathering the data shown in Figure 3b-c in the main text and Figure S40-42 in the Supplementary information.

**Step 1**: 104 µL buffer was added to each well.

**Step 2:** 16 µL *M_1_L* (100 nM stock) was added to each well.

**Step 3:** 20 µL reporter (200 nM stock) was added to each well.

**Step 4:** 10 µL *P* (1 µM stock) was added to each well.

**Step 5:** 0-24 µL *T* (50 nM stock) and 50-26 µL of buffer was injected to each well to trigger the reactions.

**Step 6:** After few days, Cy3 signals were checked to see if they all had reached a similar value. If not, the reactions were allowed more time, and checked again.

**Step 7:** 4 µL *M_1_*/*M_2_*/*M_3_* (2 µM stock) was added to each well to saturate the reporter and estimate its initial concentration.

Table S5: Experimental method for discard pathway assays

| **Step** | **Component added** | **Purpose** | **Target concentration in 200 µL** | **Volume in µL** | **Measurement no.** |
| --- | --- | --- | --- | --- | --- |
| Step 1 | 104 µL buffer | Baseline fluorescence | n/a | 104 | 1 |
| Step 2 | 16 µL *ML* (100 nM stock) | Residual fluorescence from ML | 8 nM *ML* | 120 | 2 |
| Step 3 | 20 µL reporter (200 nM stock) | Residual fluorescence from reporters | 20 nM reporter | 140 | 3 |
| Step 4 | 10 µL *P* (1 uM stock), 40 µL buffer | Fluorescence increase with P addition | 50 nM *P* | 150 | 4 |
| Step 5 | 0-24 µL *T* (50 nM stock) | Trigger reaction | 0-6 nM *T* | 200 | 5 |
| Step 6 | n/a | Check anomalies | n/a | 200 | 6 |
| Step 7 | 4 µL *M* (2 µM stock) | Saturate the reporters | ~48 nM | 204 | 7 |

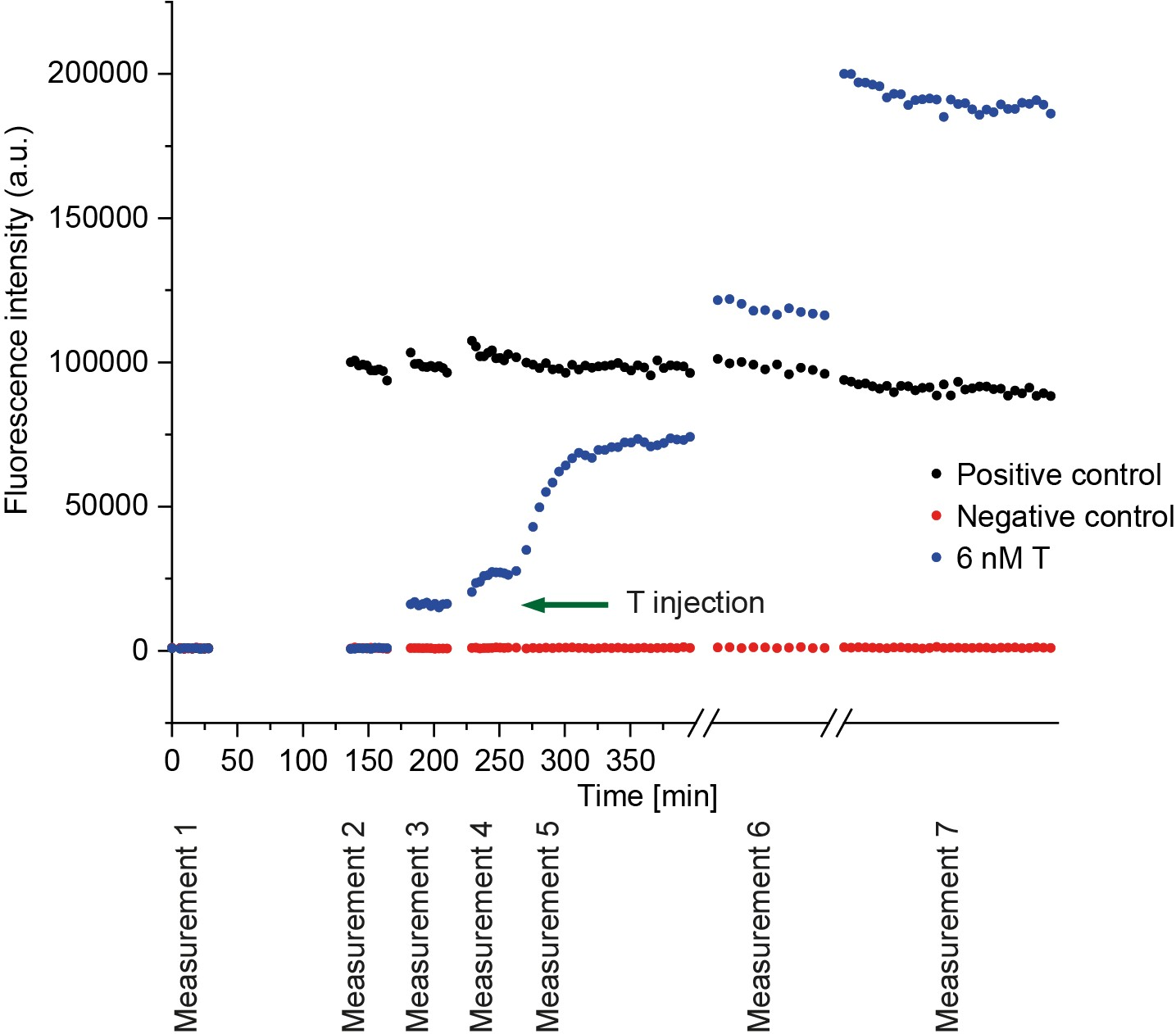

Figure S8: Example data for characterisation of the discard pathway of the M_2_T complex tiggered by P_7_ in AlexaFluor-647 channel

The AlexaFluor-647 fluorescence signal shows the formation of *M_2_PR* complex triggered by the addition of 6 nM of *T* into a solution containing 20 nM *RQ,* 8 nM *M_2_L,* and 50 nM *P_7_*. Measurement 1 shows the baseline from the plate and buffer. Measurement 2 gives the change in fluorescence caused by adding 8 nM *M_2_L* complex. Measurement 3 gives the residual fluorescence from the 20 nM *RQ* complex. Measurement 4 shows the change in fluorescence upon adding 50 nM *P_7_*. Then the reaction is triggered by adding 6 nM *T* and measurement 5 occurs for several hours. Measurement 6 was performed after several days to ensure that the reaction is completed. For measurement 7, the reporter was saturated by the addition of another 40 nM of free *M_2_* which reacts with excess proofreader in the solution to form *M_2_P*, which then triggers the remaining reporter to estimate the initial concentration of *RQ*.

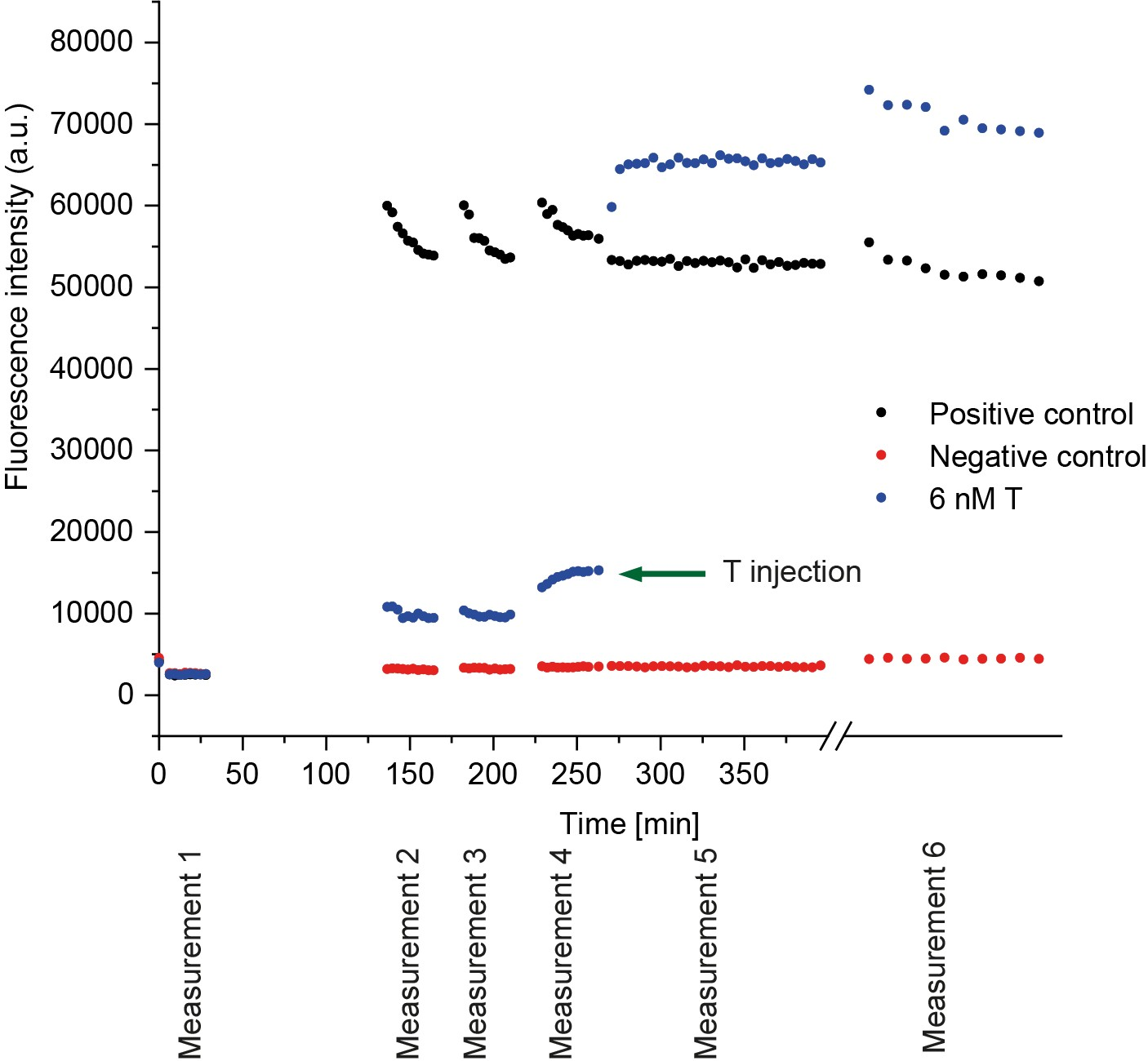

Figure S9: Example data for characterisation of the discard pathway of the M_2_T complex triggered by P_6_ in Cy3 channel

Cy3 fluorescence signal shows the displacement of strand *L* from the *M_2_L* complex by the template. Measurement 1 shows the baseline from the plate and buffer. Measurement 2 measures the change in fluorescence by adding 8 nM *M_2_L* complex. Measurement 3 measures the residual fluorescence from the 20 nM *RQ* complex. Measurement 4 shows the direct invasion of *M_2_L* by the added proofreader strand *P_7_*. Then the reaction is triggered by adding 6 nM *T* and measurement 5 was performed for several hours. Measurement 6 was performed after several days to ensure that the reaction was complete, and to also ensure that there is minimal evaporation, if any. The decay in the signal of Cy3 at the beginning of each measurement is characteristic of temperature increase in the experiment. These signature decays are present in almost all the experiments at its initial phase where the 96-well plate is transferred from the ambient lab temperature of 22-23°C to the plate reader kept at 25°C.

### Supplementary note 6: Protocol for dimer formation via HMSD

This method was used to gather the data shown in Figure 4a-b in the main text and Figure S44-S52 in the Supplementary information.

**Step 1:** 94 or 104 µL (for experiments done with and without proofreaders, respectively) buffer is added to each well.

**Step 2:** 16 µL *M’_1_L’* (100 nM stock) was added into each well.

**Step 3:** 10 or 0 µL *P’* (1 µM stock) was added into each well (for experiments with and without proofreader respectively).

**Step 4:** 10 µL *N* (200 nM stock) was added into each well.

**Step 5:** 0-24 µL of template (50 nM stock) and 50-26 µL of buffer was injected into each well to vary the concentrations of template.

**Step 6:** After the Cy3 signal for all reaction wells reach similar values, 20 µL reporter (200 nM stock) was added into each well to check the dimeric product formation.

Table S6: Experimental method for dimerisation via HMSD

| **Steps** | **Component added** | **Purpose** | **Target concentration in 200 µL** | **Volume in µL** | **Measurement no.** |
| --- | --- | --- | --- | --- | --- |
| Step 1 | 94 or 104 µL buffer | Baseline fluorescence | n/a | 94 or 104 | 1 |
| Step 2 | 16 µL *M’L’* (100 nM stock) | Residual fluorescence | 8 nM *M’L’* | 110 or 120 | 2 |
| Step 3 | 10 or 0 µL *P’* (1 µM stock), 40 µL buffer | Addition of proofreader | 50 nM *P’* | 120 | 3 |
| Step 4 | 10 µL *N* | Addition of second monomer | 10 nM *N* | 130 | 4 |
| Step 5 | 0-24 µL *T’* (50 nM stock) and 50-26 µL buffer | Trigger reaction | 0-6 nM of *T’* | 180 | 5 |
| Step 6 | 20 µL reporter (200 nM stock) | Measure product formation | 20 nM reporter | 200 | 6 |

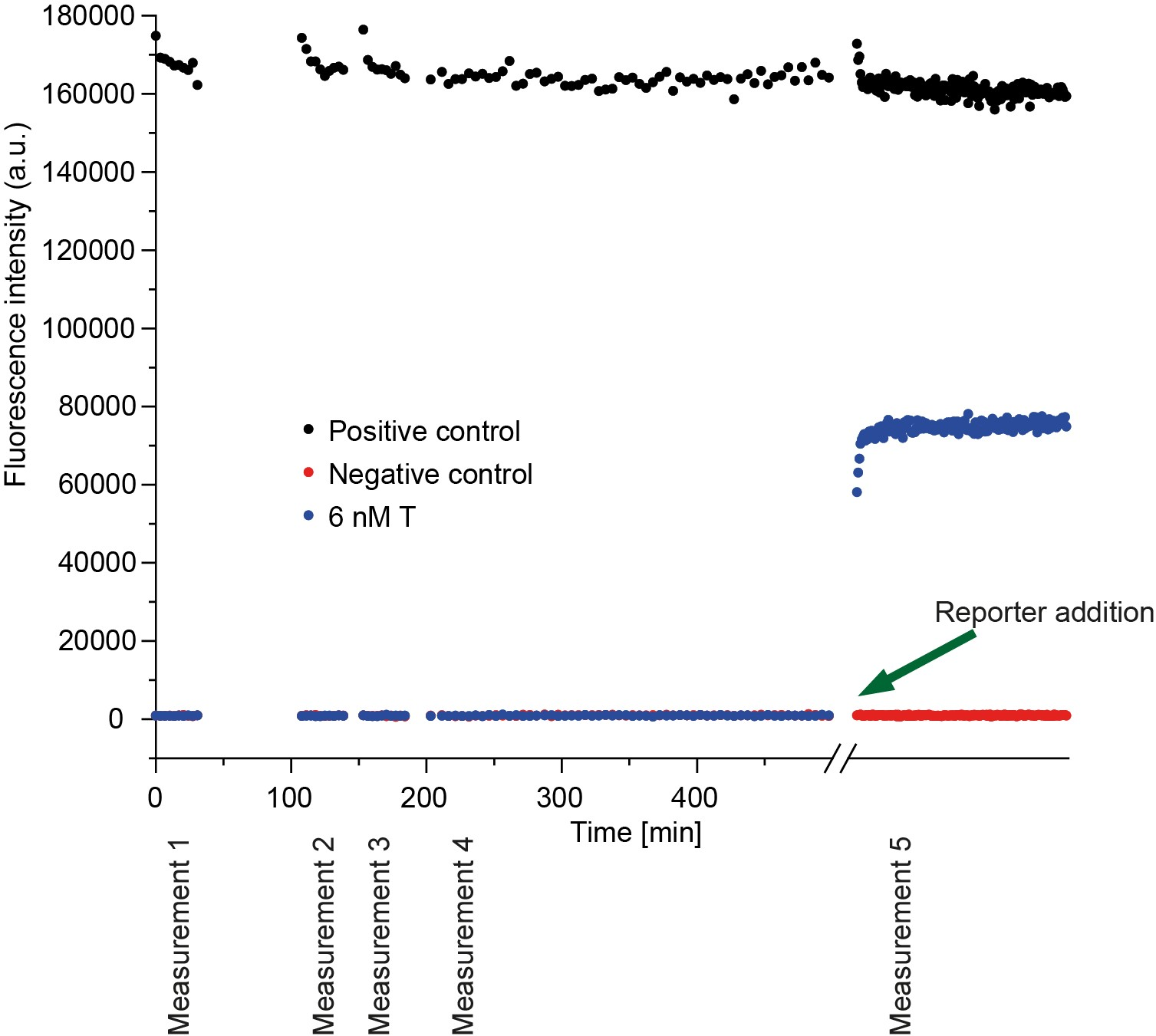

Figure S10: Example data in the AlexaFluor-647 channel for product formation in a dimerisation reaction with kinetic proofreading

AlexaFluor-648 signal showing the formation of dimeric product. 94 µL (where proofreader is added) or 104 µL (where no proofreader is added) of buffer is added into the plate well and the baseline is measured (measurement 1). For the second measurement, 8 nM *M’L’* complex is added in the well and measured. Then 10 µL or 0 µL *P’* is added to it and the fluorescence is read for the 3^rd^ measurement. 0-6 nM *T’* *is* added (6 nM in this case) to trigger the reaction and the reaction proceeds for a few days (measurement 4). Reaction progress is measured in the Cy3 channel (Figure S11). Once the corresponding Cy3 signals are plateaued, 20 nM reporter is added to measure the concentration of the formed dimer (measurement 5).

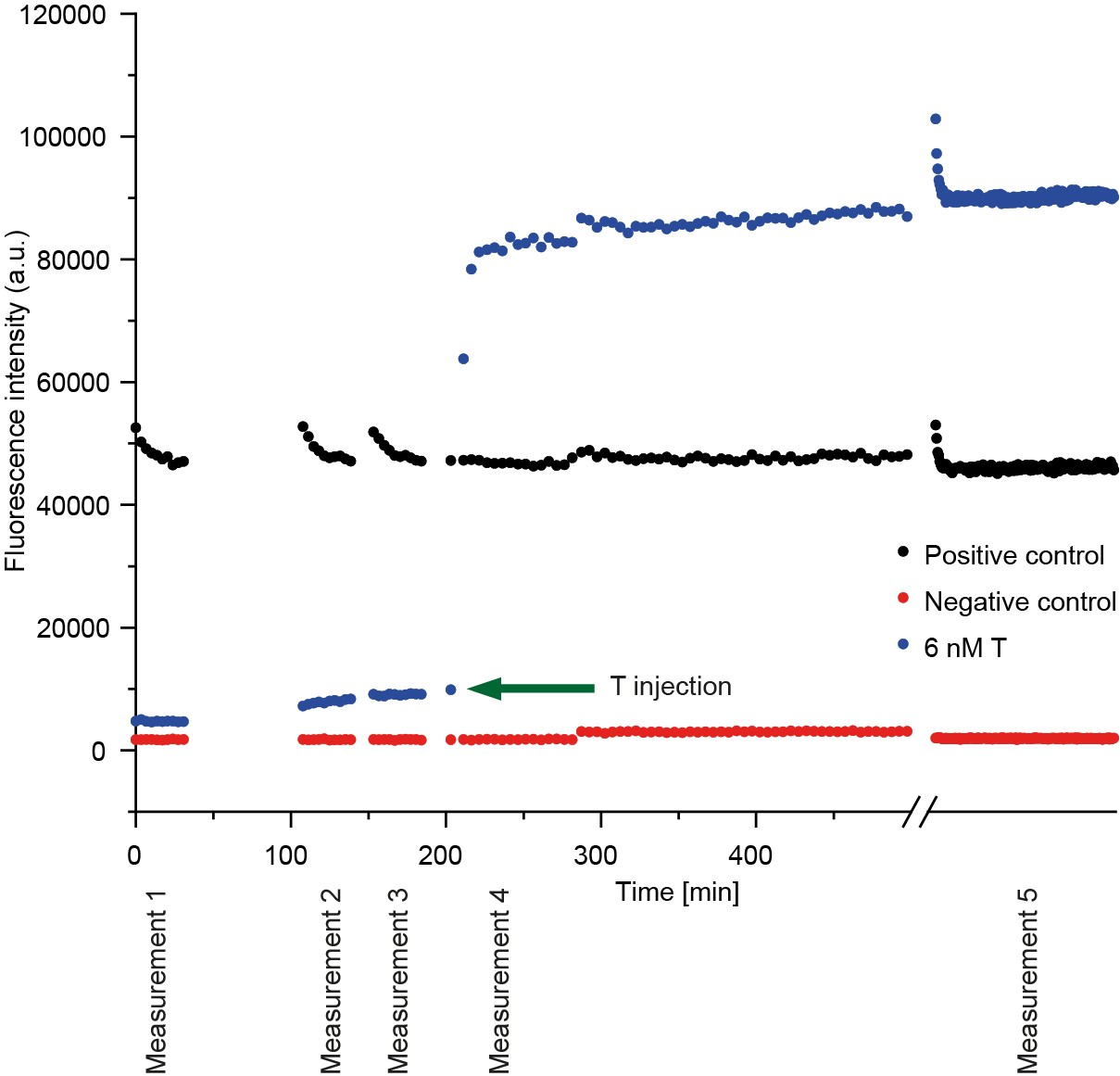

Figure S11: Example data in the Cy3 channel for the dimerisation process with kinetic proofreading

Cy3 signal showing the binding of *M’L’* to the template. 94 µL (where proofreader is added) or 104 µL (where no proofreader is added) of buffer is added into the plate well and the baseline is measured (measurement 1). For the second measurement, 8 nM *M’L’* complex is added in the well and measured. Then 10 µL or 0 µL *P’* is added to it and the fluorescence is read for the 3^rd^ measurement. 0-6 nM *T’* *is* added (6 nM in this case) to trigger the reaction and the fluorescence is monitored for few days as measurement 4 to check the progress of the reaction. Once the Cy3 signals are plateaued, 20 nM reporter is added to measure the concentration of the formed dimer (measurement 5, using the AlexaFluor-647 channel, Figure S10). Temperature-dependent decay of Cy3 signal is also observed here, as in Figure S9.

### Supplementary note 7: Protocol for dimer formation via HMSD from a mixture of monomers

This method was used to gather the data shown in Figure 4c-d in the main text and Figure S55-S60 in the Supplementary information. These experiments were performed in 3 identical sets. The only difference is in ‘Step 6’ where different reporters were added in each set to check separate dimeric products. Since all the reporters had the same fluorophore (AlexaFluor-647), they were not distinguishable in a single experiment.

**Step 1:** 85 or 95 µL (for experiments done with and without proofreaders, respectively) buffer is added to each well.

**Step 2:** 10 µL of each *M’_1_L’*, *M’_2_L’* and *M’_3_L’* (100 nM stock) was added into each well.

**Step 3:** 10 or 0 µL *P’* (1 µM stock) was added into each well (for experiments with and without proofreader respectively).

**Step 4:** 15 µL *M’_2_* (200 nM stock) was added into each well.

**Step 5:** 0-24 µL of template (50 nM stock) and 50-26 µL of buffer was injected into each well to vary the concentrations of template.

**Step 6:** After the Cy3 signal for all reaction wells reach similar values, 10 µL reporter (200 nM stock) was added into each well to check the dimeric product formation.

Table S7: Experimental method for dimerisation via HMSD from a mixture of monomers

| **Steps** | **Component added** | **Purpose** | **Target concentration in 200 µL** | **Volume in µL** | **Measurement no.** |
| --- | --- | --- | --- | --- | --- |
| Step 1 | 85 or 95 µL buffer | Baseline fluorescence | n/a | 85 or 95 | - |
| Step 2 | 10 µL *M’_1_L’, M’_2_L’, M’_3_L’* (100 nM stock) | Residual fluorescence | 5 nM each of *M’_1_L’, M’_2_L’, M’_3_L’* | 115 or 125 | - |
| Step 3 | 10 or 0 µL *P’* (1 µM stock), 40 µL buffer | Addition of proofreader | 50 nM *P’* | 125 | - |
| Step 4 | 15 µL *N* | Addition of second monomer | 15 nM *N* | 140 | - |
| Step 5 | 0-24 µL *T’* *(*50 nM stock) and 50-26 µL buffer | Trigger reaction | 0-6 nM of *T’* | 190 | 1 |
| Step 6 | 20 µL reporter (200 nM stock) | Measure product formation | 20 nM reporter | 200 | 2 |
| Step 7 | n/a | Product formation in long term |  | 200 | 3 |

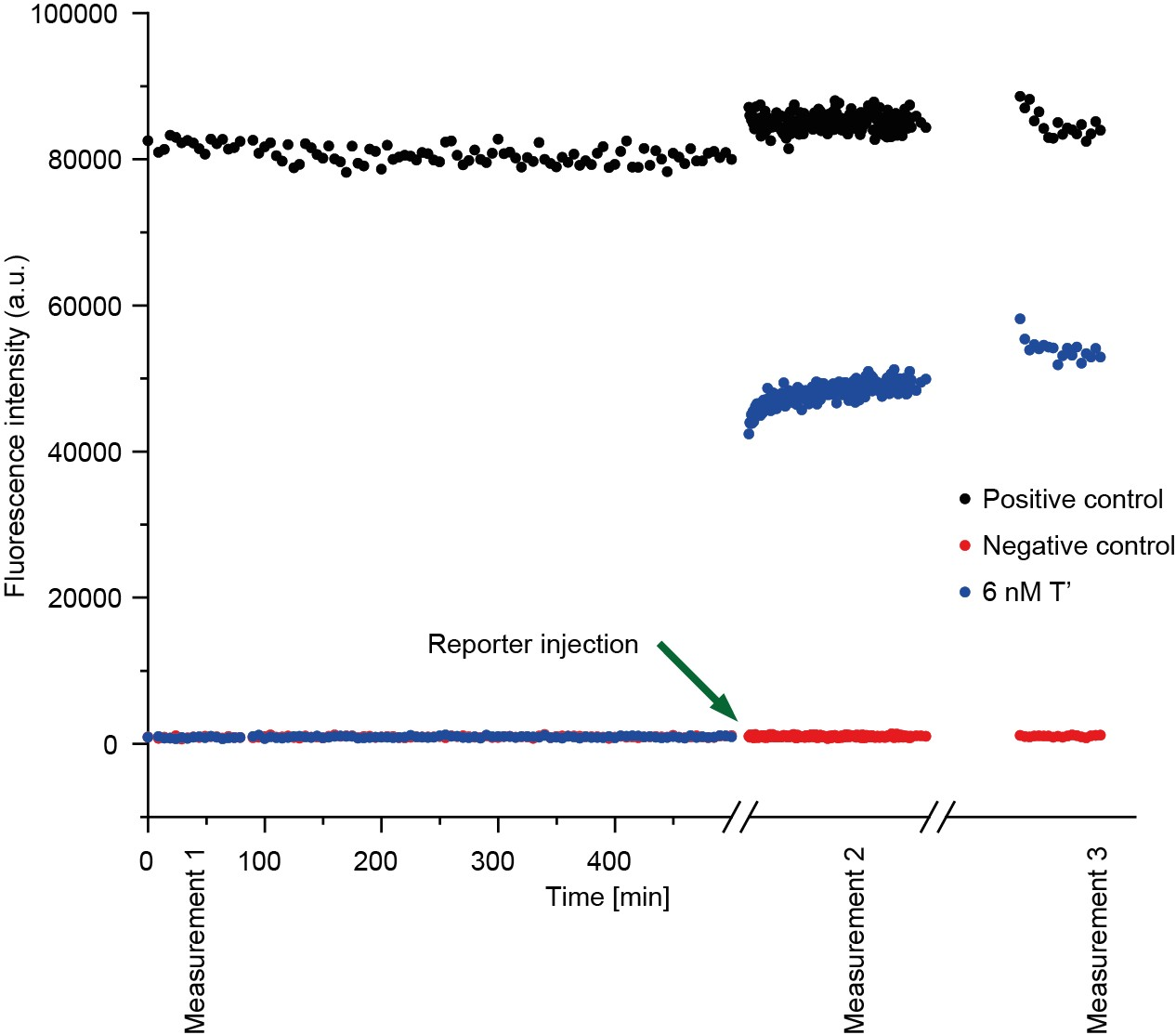

Figure S12: Example data of product formation in a dimerisation reaction with proofreading from an equimolar mixture of three different monomers monitored by AlexaFluor-647 fluorescence

AlexaFluor-648 signal showing the formation of dimeric product. 85 µL (where proofreader is added) or 95 µL (where no proofreader is added) of buffer is added into the plate well. Then 5 nM of each *M’L’* complex is added in the well. Then 10 µL or 0 µL *P’* is added to it. The reaction is triggered by adding 0-6 nM *T’* *is* added (6 nM in this case) to trigger the reaction and the fluorescence is monitored in the Cy3 channel (Figure S13) for few days as measurement 1. Once the corresponding Cy3 signals are plateaued, 10 nM reporter is added to measure the concentration of the formed dimer (measurement 2).

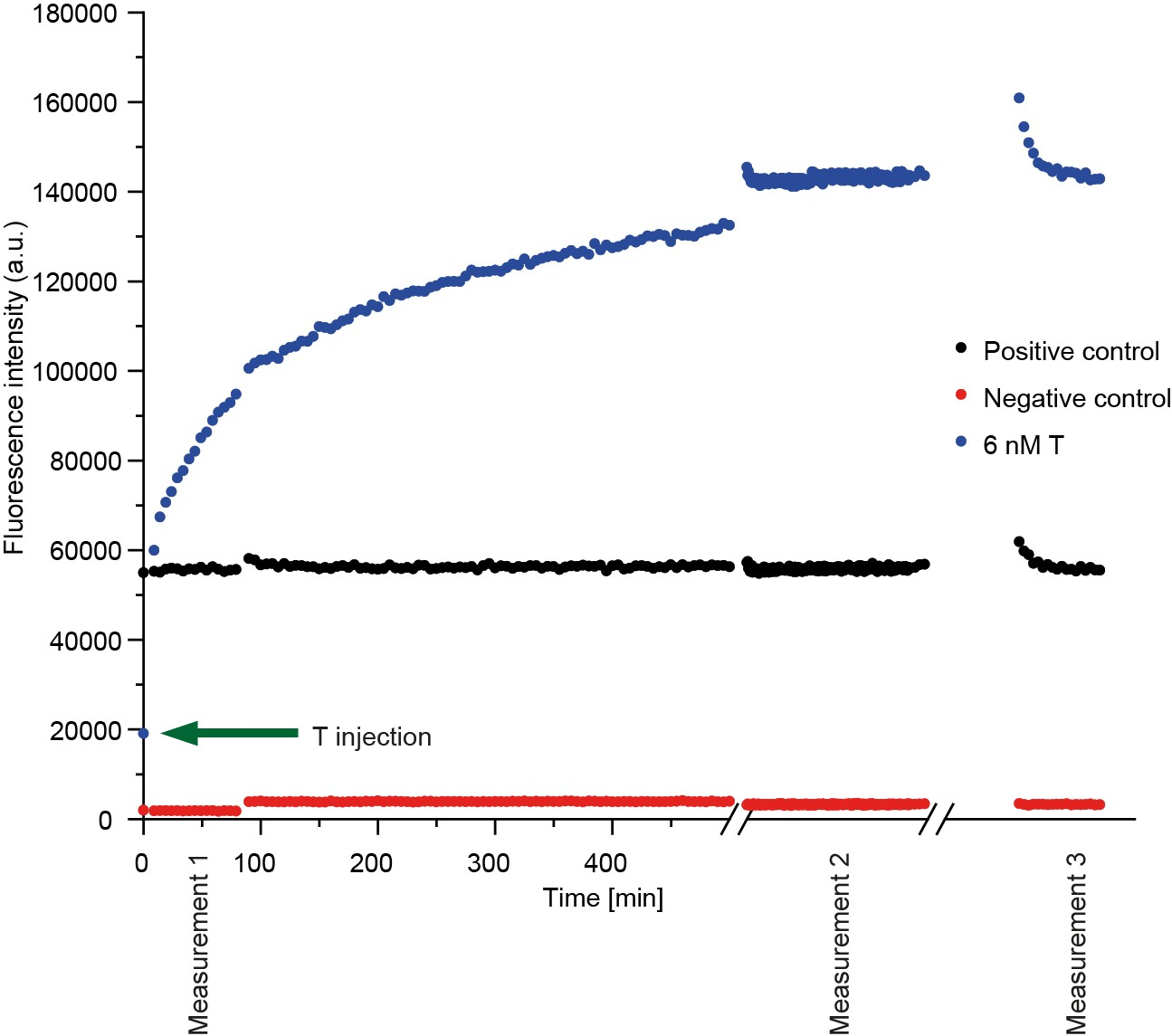

Figure S13: Example data in the Cy3 channel for the dimerisation process with proofreading from an equimolar mixture of three different monomers

Cy3 signal showing the formation of dimeric product. 85 µL (where proofreader is added) or 95 µL (where no proofreader is added) of buffer is added into the plate well. Then 5 nM of each *M’L’* complex is added in the well. Then 10 µL or 0 µL *P’* is added to it. The reaction is triggered by adding 0-6 nM *T’* *is* added (6 nM in this case) to trigger the reaction and the progress of the reaction is monitored by measuring Cy3 signal (measurement 1). Once the signals are plateaued, 10 nM reporter is added to measure product formation (measurement 2 and 3, signal observed in the AlexaFluor-647 channel as shown in Figure S12).

### Supplementary note 8: Protocol for SNP detection

This protocol was used to gather experimental data shown in Figure 5b-c in the main text and Figure S61 in the Supplementary information.

**Step 1:** 85-95 µL buffer was added into each well.

**Step 2:** 15 µL *TS* or *SNP* (200 nM stock) was added into each well.

**Step 3:** 20 µL sink complex (200 nM stock) was added into each well.

**Step 4:** 0-10 µL *P_snp_* was added in different wells to vary its concentration.

**Step 5:** 10 µL probe-blocker complex (200 nM stock) and 40 µL buffer was added into each well to trigger the reactions. The progress of the reactions was monitored by following the Cy3 fluorescence channel.

**Step 6:** The plate was scanned again after about 4 weeks to check for long-term triggering of the reporters.

Table S8: Experimental method for SNP detection

| **Steps** | **Component added** | **Purpose** | **Target concentration in 200µL** | **Volume in µL** | **Measurement no.** |
| --- | --- | --- | --- | --- | --- |
| Step 1 | 85- 95 µL buffer | Baseline fluorescence | n/a | 85-95 |  |
| Step 2 | 15 µL candidate strand | - | 15 nM candidate strand | 100-110 |  |
| Step 3 | 20 µL reporter (200 nM stock) | - | 20 nM reporter | 120-130 |  |
| Step 4 | 20 µL Sink complex | - | 20 nM Sink complex | 140-150 |  |
| Step 5 | 0-10 µL *P* (1 µM stock), 40 µL buffer | - | 0-50 nM *P* | 150 |  |
| Step 6 | 10 µL Probe-blocker complex, 40 µL buffer | Trigger reaction | 10 nM Probe-blocker complex | 200 | Measurement 1 |
| Step 7 | - | Long term signal | - | 200 | Measurements 2-4 |

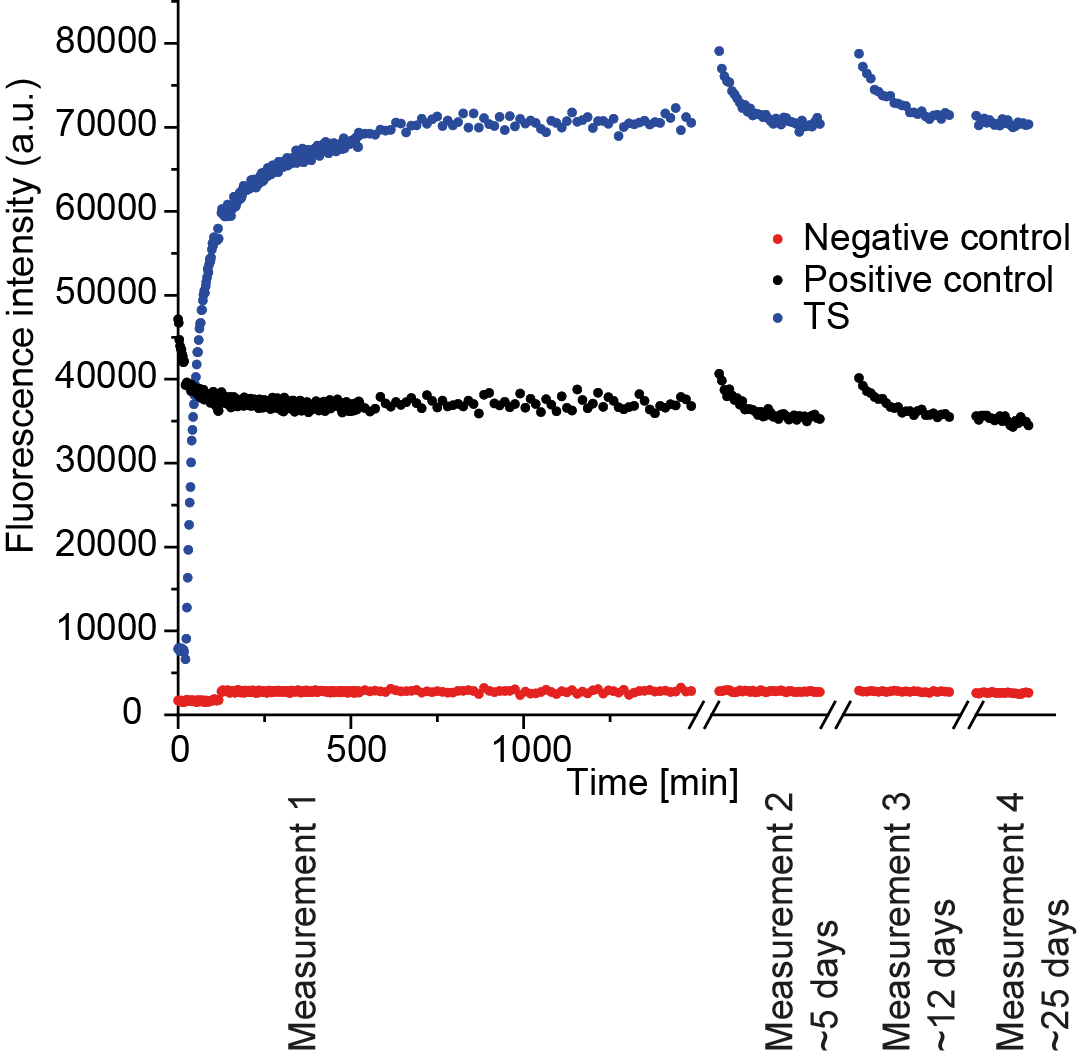

Figure S14: Example data of Cy3 signal from SNP detection scheme

Cy3 signal obtained from the external reporters in the SNP detection system. 15 nM of the *TS* strand, 20 nM of sink complex, 20 nM of the reporter, and 0-50 nM of the proofreader were mixed with buffer. The reaction was triggered by adding 10 nM of Probe-blocker complex. Produced *TS-Probe-Reporter* complexes generate the signal (measurement 1). Several short measurements were taken between 5 to 25 days to check the long-term generation of Cy3 signal (measurement 2-4).

### Supplementary note 9: Behaviour of *M’_1_* during dimerization experiments.

Figure 3a and b of the main text both show an approximately equal yield of *M’*_1_*N*, with or without proofreader P’_6_. This fact is perhaps surprising, since we would expect at least some of the monomers to be removed by the proofreader prior to dimerization. Indeed, the data on unblocked monomer concentration in the same figure shows an increased rate of monomer unblocking in the presence of proofreader – presumably due to template recovery being accelerated by template-bound monomers being removed via the discard pathway.

A clue to the resolution of this paradox is provided by the predictions of our ODE model (Supplementary note 14). The predicted *M’*_1_*N* concentration rises more slowly in the presence of proofreader P’_6_, which diverts some of the monomers from the dimerization pathway, as expected. However, the eventual yield of dimers is high – because the proofreading reaction for the correctly-matched monomer has an appreciable reverse reaction, allowing monomers to rebind to the template. This behaviour is also likely occurring in the experiment and is advantageous in achieving a high yield of the correct complex, provided that the incorrect monomers do not also participate. Both for the model as parameterized, and the experimental data (Supplementary Figure S44-52, S55-60), *M’*_2_*N* and *M’*_3_*N* concentrations remain suppressed at long times when P’_6_ is used as a proofreader*.*

The model also predicts that *M’*_2_*N* is not effectively suppressed by *P’_7_* at long times, due to the reverse proofreading reaction; this effect is not observed in experiment. The precise values of *k*_4_ are not well constrained by our fitting procedure; small errors in concentrations of reagents would feed through into very different estimates for these constants. We also note that the *Sink* complex supresses similar behaviour in the SNP detection scheme.

### Supplementary note 10: Fluorescence data processing and ODEs for lock opening reaction

The lock opening reaction is given by

$ML+T \begin{matrix} k_{1} \\ \leftrightharpoons\\ k_{2} \end{matrix} MT+L$.	(S1)

The ordinary differential equation (ODE) that can completely describe the system is:

$\frac{d}{dt}\left[ MT \right]= k_{1}\left[ ML \right]\left[ T \right]-k_{2}\left[ MT \right][L]$.	(S2)

The species conservation equations used are:

$\left[ ML \right]+\left[ MT \right]=\left[ ML_{total} \right]$,	(S3)

$\left[ T \right]+\left[ MT \right]=[T_{total}]$,	(S4)

$\left[ L \right]+[ML]=1.2\left[ ML_{total} \right]$.	(S5)

[*ML_total_*] and [*T_total_*] are the initial concentrations of locked monomer and the template, respectively. There is excess lock present in the solution when the reaction starts. This initial lock concentration is 20% of the initial [*ML*] concentration, i.e., 0.2[*ML_total_*]. All other species have 0nM initial concentration. To estimate the rate parameters {*k_1_*, *k_2_*}, experiments are performed involving the locked monomer with varying concentrations of the template *T*, with fluorescence monitored in the Cy3 channel. A fluorescent signal is produced as shown in Figure S15.

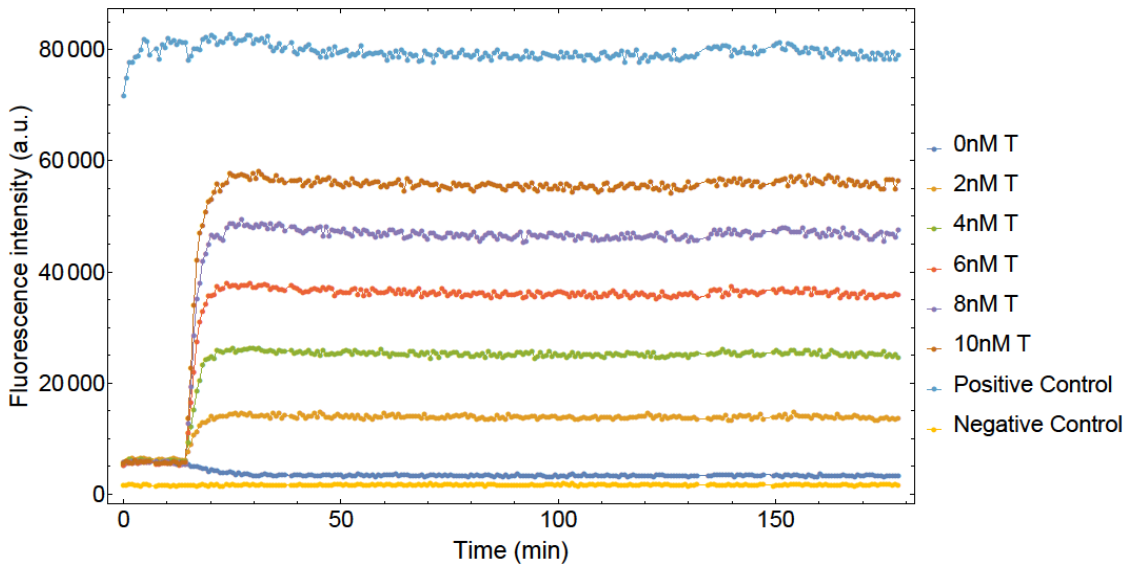

Figure S15: Fluorescent signal from the Cy3 channel obtained for the template binding reaction of M_1_L for varying [T_total_].

The jump in the fluorescent signal is the injection of template *T*. We redefine the time axis such that the point of injection is placed at t = 0 and then convert the fluorescent signal to concentration of the desired species, *MT* . The background fluorescence before injection is primarily due to the presence of *ML*. After injection, a mixture of *MT* and *ML* cause the fluorescence. We estimate the fraction of converted *ML* (equal to [*MT* ]) as:

$\frac{[MT]}{[ML_{total}]}=\frac{f\left( t \right)-X_{o}(t)}{P\left( t \right)-X_{o}(t)}$	(S6)

where *P(t)* is the positive control signal, which in this case corresponds to a system with [*M*]=[*ML_total_*], and *X_o_(t)* is the signal from the channel when no template was added (negative control). *f(t)* is the total fluorescence intensity at time *t* in the experiment in question. [*ML_total_*] is calculated by taking the mean of the difference between the positive control and negative control signals over time.

$\left[ ML_{total} \right]=\frac{\bar{P\left( t \right)}-\bar{X_{o}\left( t \right)}}{F_{MT,C}}$	(S7)

where *FMT* *_,C_* is the fluorescence signal density of *MT* in the Cy3 channel. This proportionality factor was obtained from initial calibration experiments, Supplementary note 17. [*ML_total_*] was estimated as 9.64nM for all 3 monomers *M_1_*, *M_2_* and *M_3_*. [*T_total_*] is assumed to be the same before and after injection. Figure S16 shows the inferred concentration of *MT* with time for different [*T_total_*].

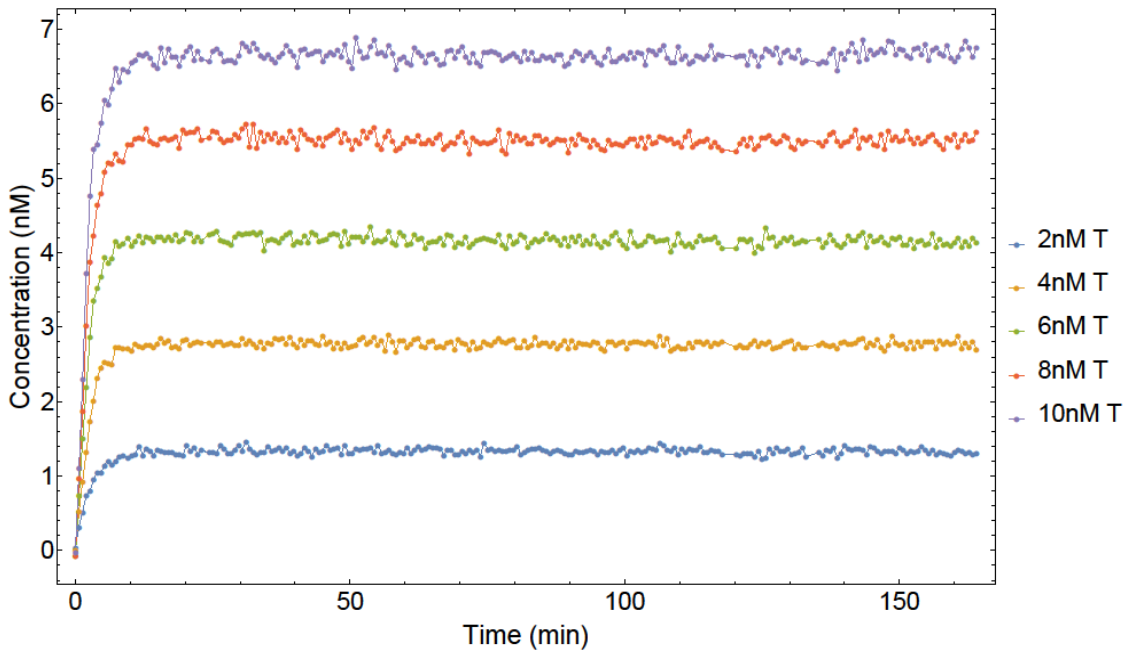

Figure S16: Concentration of unlocked template M_1_T as a function of time.

The time-dependence of [*MT* ] is used to estimate the forward (*k_1_*) and backward (*k_2_*) reaction rate constants. To calculate the best parameter fits, MSE analysis as shown in Eq. (3), was performed over [*MT* ] using the system described by Eqs.(S2)-(S5). *k_1_* and *k_2_* were sampled from values between 8.33 M^-1^ sec^-1^-8.33E+05 M^-1^ sec^-1^. The predicted {*k_1_*, *k_2_*} rate constants for the lock opening reaction can be found in Table S9.

### Supplementary note 11: Fluorescence data processing and ODEs for reporter characterization

The next step is to identify the rate of reporter reacting with the monomer bound to the proof-reader, {*k_5_*, *k_6_*}. The reaction occurs as follows.

$MP+RQ \begin{matrix} k_{5} \\ \leftrightharpoons\\ k_{6} \end{matrix} MPR+Q$.	(S8)

The ordinary differential equation (ODE) that can completely describe the system is:

$\frac{d}{dt}[MPR]= k_{5}\left[ MP \right][RQ]-k_{6}[MPR][Q]$.	(S9)

The species conservation equations that were used are as follows.

$\left[ MP \right]+\left[ MPR \right]=[MP_{total}]$,	(S10)

$\left[ RQ \right]+\left[ MPR \right]=\left[ RQ_{total} \right]$,	(S11)

$\left[ RQ \right]+[Q]=1.2[RQ_{total}]$	.	(S12)

[*MP_total_*] and [*RQ_total_*] are the initial concentrations of monomer bound proofreader and reporter respectively. There is excess quencher *Q* present in the solution when the reaction starts. This initial quencher concentration is 20% of the initial [*RQ*] concentration, i.e., 0.2[*RQ_total_*]. All other species have 0nM initial concentration.

To study the reporter reaction, experiments were performed between monomer-bound proofreader, *MP*, and reporter, *RQ* in the Cy3 and Alexa channels. The fluorescence intensities from the Cy3 and Alexa channels for monomer *M_1_* and proofreader *P_8_* are shown in Figure S17.

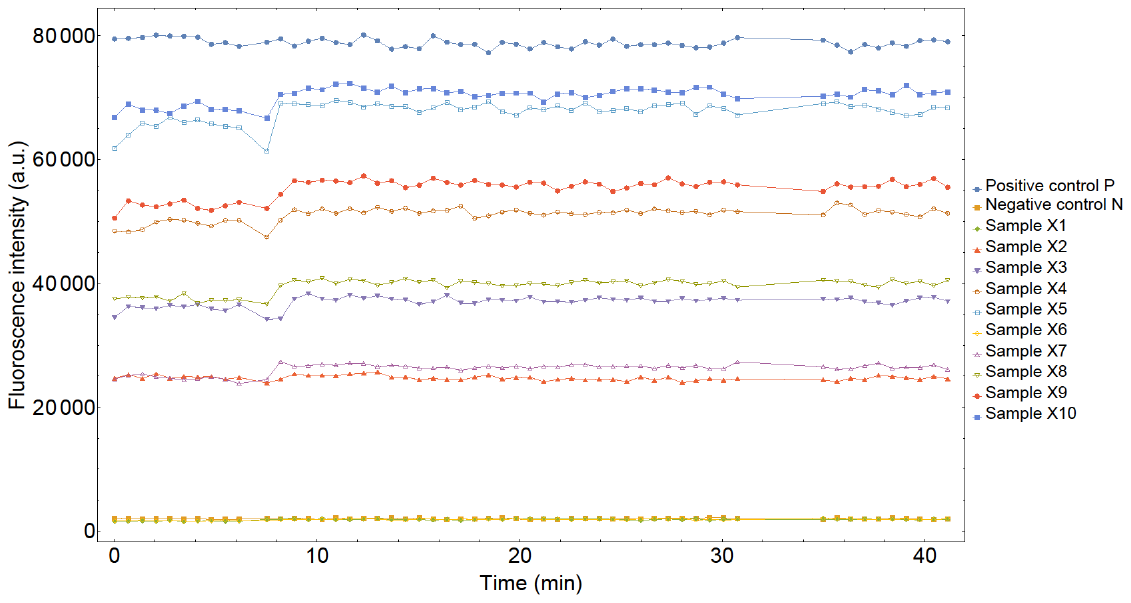

(a)

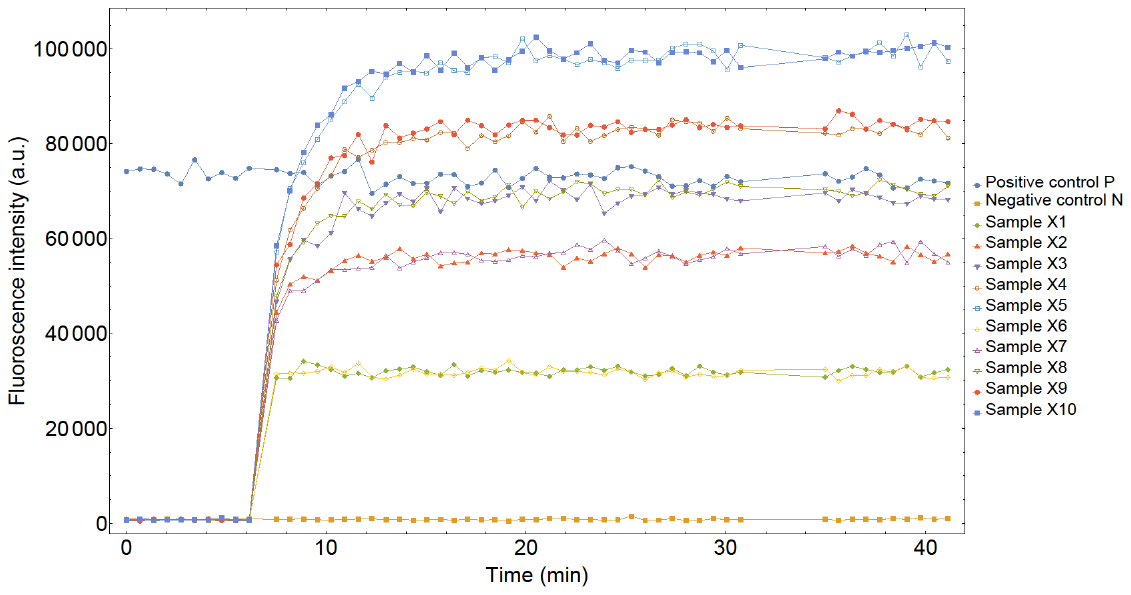

(b)

Figure S17: Fluorescent signal from the Cy3 channel (a) and the Alexa channel (b) obtained for the reporter reaction of M_1_P_8_

Samples *X1*-*X10* correspond to experiments with varying [*MP_total_*] performed in duplicate. Sample pairs {X1, X6; X2, X7; X3, X8; X4, X9; X5, X10} had intended [MP_total_] of {0nM; 4nM; 6nM; 8nM; 10nM}, respectively.

The solution contained *MP* prior to the injection point (first jump in Alexa channel). *RQ* was injected at approximately 7 minutes. The injected *RQ* concentration was intended to be the same across all experiments. Each experiment was performed in duplicate with the following sample pairs having same [*MP_total_*]: {*X1*,*X6*}; {*X2*, *X7*}; {*X3*, *X8*}; {*X4*, *X9*}; {*X5*, *X10*}. The Cy3 fluorescence signal densities of *MP* and *MPR*, Supplementary note 17, are similar and so the signal obtained from this channel is relatively flat.

After subtracting the negative control, we average the sample fluorescence values. Sample $\bar{X1}$ is the average over Sample *X1* and *X6*, sample $\bar{X2}$ is the average over Sample *X2* and *X7*, *etc*. We then normalize the time and fluorescence values at t=0 as follows. For each sample in the Cy3 channel, the injection time is set at t=0 and the fluorescence value just after point of injection is considered as that sample’s fluorescence value at t=0. For each sample in the Alexa channel, the injection time is set at t=0. Since the reporter itself has non-zero fluorescence, and the signal changes quite quickly, neither the time point before injection, nor the time point after injection, give a good estimate of the fluorescence at the initial time. We therefore assume that all samples have the same fluorescence value as Sample $\bar{X1}$ (which has 0nM [*MP_total_*]) at *t*=0. Having performed these data processing steps, we obtain Figure S18.

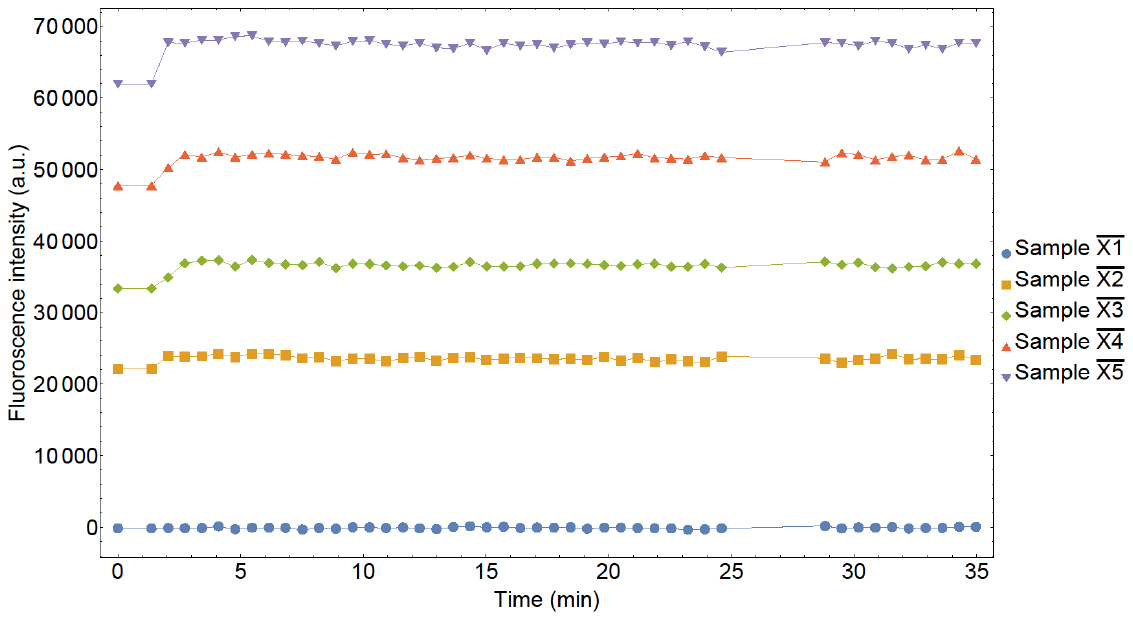

(a)

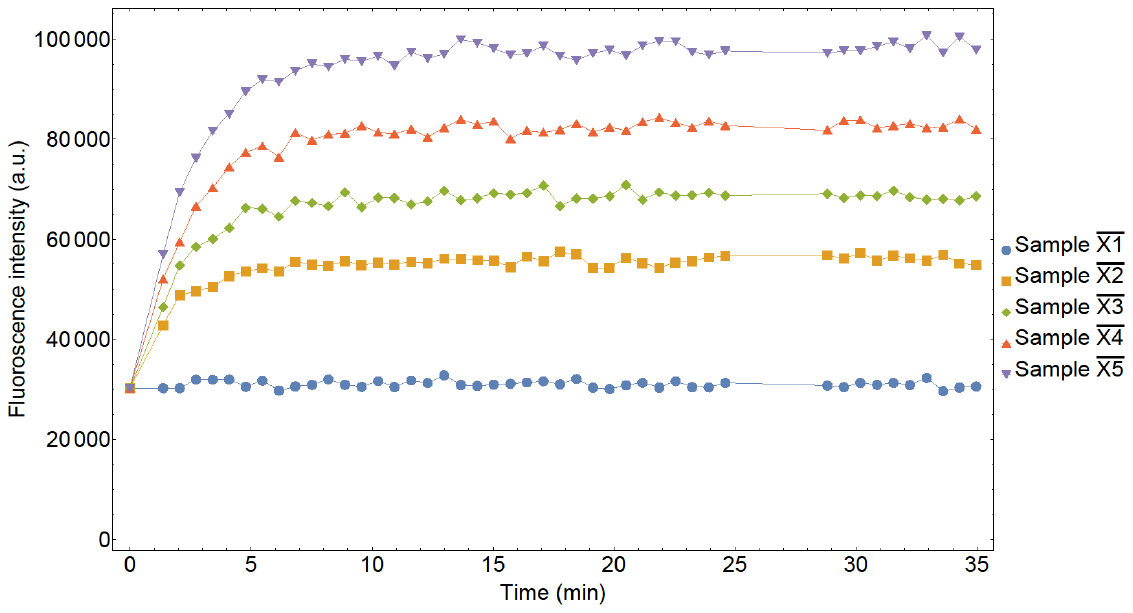

(b)

Figure S18: Processed fluorescent signal from the Cy3 channel (a) and the Alexa channel (b) obtained for the reporter reaction of M_1_P_8_

The next step is to extract relevant information from the 2 channels. The fluorescence in the Cy3 channel is caused by [*MP*] + [*MPR*], which is an estimate of the total monomer in the system, [*MP_total_*], Eq.(S10). The monomer concentration time signal, [*MP_total_(t)*] can be obtained as:

$[{MP}_{total}(t)]=\frac{f\left( t \right)_{C}}{F_{MP,C}}$ ,	(S13)

where *f(t)_C_* is the fluorescence intensity from the Cy3 channel and *F_MP,C_* is the fluorescence signal density of *MP* in the Cy3 channel, Supplementary note 17.

The fluorescent signal from the Alexa channel is used to estimate [*RQ_total_*] and the time varying *MPR* signal, [*MPR(t)*]. [*RQ_total_*] is calculated as

$[RQ_{total}]=\frac{1}{5}\frac{\sum_{I=1}^{5} f\left( t=0,\bar{X_{I}} \right)_{A}}{F_{RQ,A}}$ ,	(S14)

where $f\left( t=0,\bar{X_{I}} \right)_{A}$ is the fluorescence signal of Sample $\bar{X_{I}}$ in the Alexa channel at time *t*=0 and *F_RQ,A_* is the fluorescence signal density of *RQ* in the Alexa channel, Supplementary note 17. [*RQ_total_*] was calculated as 44.86nM for *M_1_P_8_*.

To estimate [*MPR(t)*], we make the following calculation. *f(t)_A_* consists of signal from *RQ* and *MPR*

${\left[ RQ \right]F}_{RQ,A}+{[MPR]F}_{MPR,A}=f\left( t \right)_{A}$,	(S15)

where *F_MPR,A_* is the fluorescence signal density of *MPR* in the Alexa channel. Combining Eq.(S11) and Eq.(S15) gives

$[MPR]=\frac{f\left( t \right)_{A}-\left[ RQ_{tot} \right]F_{RQ,A}}{F_{MPR,A}-F_{RQ,A}}$.	(S16)

The processed fluorescence signals from the Cy3 and Alexa channels are converted to total monomer concentration and *MPR* concentration, respectively; the results are shown Figure S19.

To obtain [*MP_total_*] in the system, we take the mean of all datapoints (except the first 10) of the concentration signal in the Cy3 channel. This gives us the following values of [*MP_total_*] from Sample $\bar{X1}$ to $\bar{X5}$: {0.01nM, 2.62nM, 4.08nM, 5.75nM, 7.50nM}.

It was noted that the initial *MP* concentrations obtained from the Cy3 channel did not match the initial concentrations intended when the solution was created. For each system under study, the intended initial *MP* concentrations were {0nM, 4nM, 6nM, 8nM, 10nM} whereas the ones obtained in this case (*M_1_P_8_*) are {0.01nM, 2.62nM, 4.08nM, 5.75nM, 7.50nM}. Moreover, in some cases the initial *MP* concentration indicated by the Cy3 channel was smaller than the apparent yield of *MPR* indicated by the Alexa channel. This inconsistency between Cy3 and Alexa channels made completing an unambiguous fit of the reaction rate constants challenging.

We therefore performed the fitting for 8 sets of initial *MP* concentrations, with the lower limit of the set defined by the concentrations generated from the Cy3 channel and the upper limit defined by the intended initial *MP* concentrations. For each set of initial *MP* concentration, the MSE calculation as shown in Eq. (3) was performed on the time varying [*MPR*] signal to generate 8 different sets of reaction rate constants {*k_5_,k_6_*}. The optimal set was identified by running 2 sets of MSE calculation on the [*MPR*]+[*PR*] signal extracted from the template recovery (Supplementary note 12) dataset (MSE_1_) and discard pathway (Supplementary note 13) dataset (MSE_2_). For each set of initial *MP* concentration, parameters *k_3_, k_4_, k_7_* were obtained by minimizing MSE_1_ + MSE_2_. Finally, the set of initial *MP* concentrations (and their corresponding k_5_, k_6_ values) that had the lowest MSE_1_+MSE_2_ was used to identify *k_3_, k_4_, k_5_, k_6_, k_7_.* For each MSE calculation, *k_3_, k_4_, k_5_, k_6_* were sampled from values between 8.33 M^-1^ sec^-1^-8.33E+05 M^-1^ sec^-1^. *k_7_* was sampled from values between 1 M^-1^ sec^-1^-1.0E+05 M^-1^ sec^-1^. The predicted {*k_5_*, *k_6_*} rate constants can be found in Table S10.

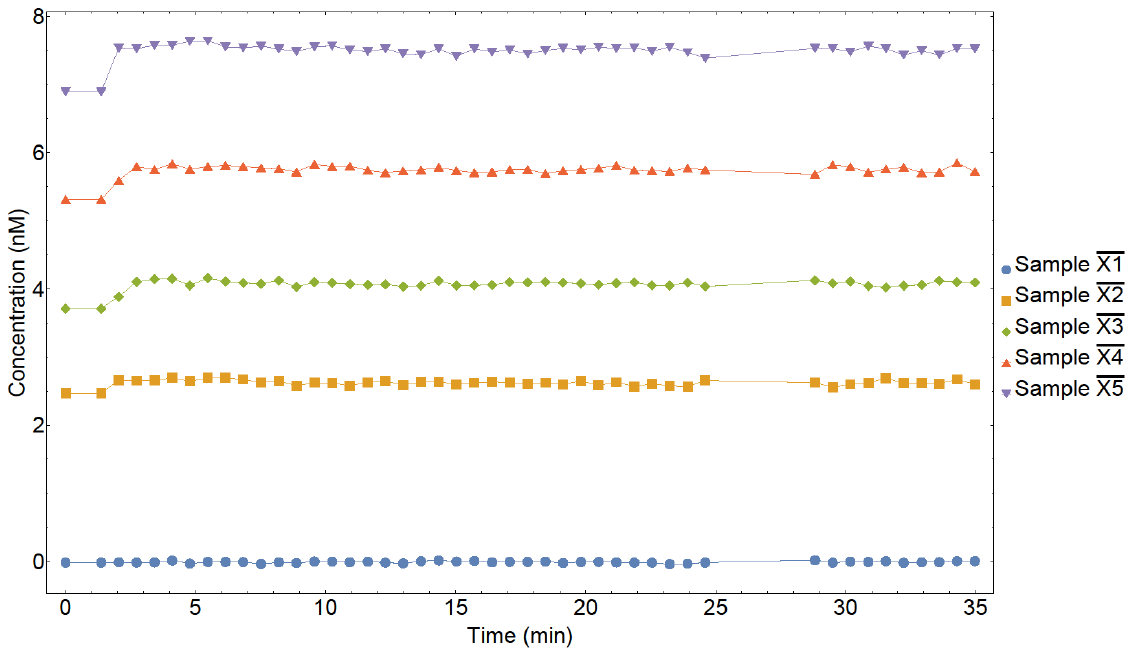

(a)

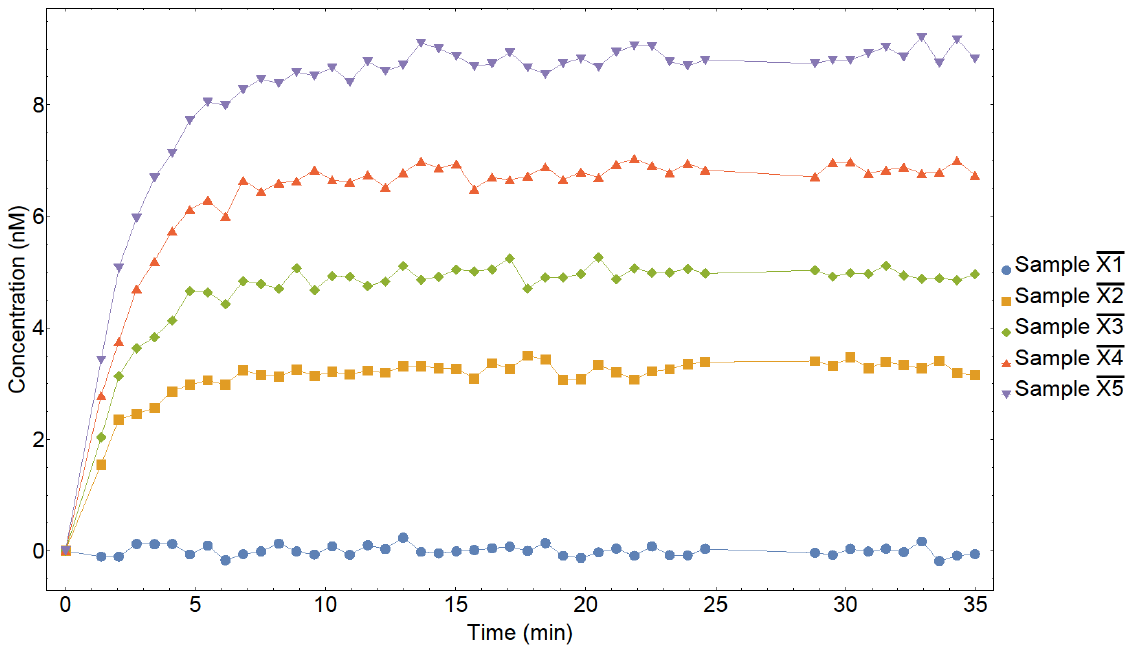

(b)

Figure S19: Total monomer concentration from the Cy3 channel (a) and [MPR(t)] time signal from the Alexa channel (b) obtained for the reporter reaction of M_1_P_8_

### Supplementary note 12: Fluorescence data processing and ODEs for template recovery

The template recovery reaction proceeds with the addition of proofreader to a monomer-bound template:

$MT+P \begin{matrix} k_{3} \\ \leftrightharpoons\\ k_{4} \end{matrix} MP+T.$	(S17)

The reaction regenerates the template while producing the monomer-bound proofreader, which is detected by the reporter quencher pair as per the following reaction:

$MP+RQ \begin{matrix} k_{5} \\ \leftrightharpoons\\ k_{6} \end{matrix} MPR+Q$	.	(S18)

A leak proofreader-triggered reporter signal reaction also proceeds as follows:
$P+RQ \underset{\to}{k_{7}}PR+Q$.	(S19)

The ordinary differential equations (ODEs) that can completely describe the system are:

$\frac{d}{dt}\left[ MP \right]= k_{3}\left[ MT \right][P]-k_{4}[MP][T]-k_{5}[MP][RQ]+k_{6}[MPR][Q]$,	(S20)

$\frac{d}{dt}[RQ]= -k_{5}[MP][RQ]+k_{6}[MPR][Q]-k_{7}[P][RQ]$,	(S21)

$\frac{d}{dt}\left[ PR \right]=k_{7}\left[ P \right]\left[ RQ \right].$	(S22)

The conservation equations are:

$\left[ MT \right]+\left[ MP \right]+\left[ MPR \right]=\left[ MT_{total} \right]$,	(S23)

$\left[ MP \right]+\left[ MPR \right]+\left[ PR \right]+\left[ P \right]=\left[ P_{total} \right],$	(S24)

$\left[ RQ \right]+\left[ PR \right]+\left[ MPR \right]=\left[ RQ_{total} \right]$,	(S25)

$\left[ MT \right]+\left[ T \right]=1.2[{MT}_{total}]$,	(S26)

$\left[ RQ \right]+\left[ Q \right]=1.2\left[ RQ_{total} \right]$	.	(S27)

[*MT* *_total_*], [*P_total_*] and [*RQ_total_*] are the initial concentrations of monomer-bound template, proofreader, and reporter respectively. There is excess quencher *Q* present in the solution when the reaction starts. This initial quencher concentration is 20% of the initial [*RQ*] concentration, i.e., 0.2[*RQ_total_*]. There is also excess template *T* present in the solution when the reaction starts. This initial template concentration is 20% of the initial [*MT*] concentration, i.e., 0.2[*MT* *_total_*]. All other species have 0nM initial concentration.

An independent reaction with *MT* *and the proofreader* was performed and monitored in the Cy3 and Alexa channel. The raw fluorescence intensities from the Cy3 and Alexa channels are shown in Figure S20 for templated monomer *M_1_* and proofreader *P_8_*.

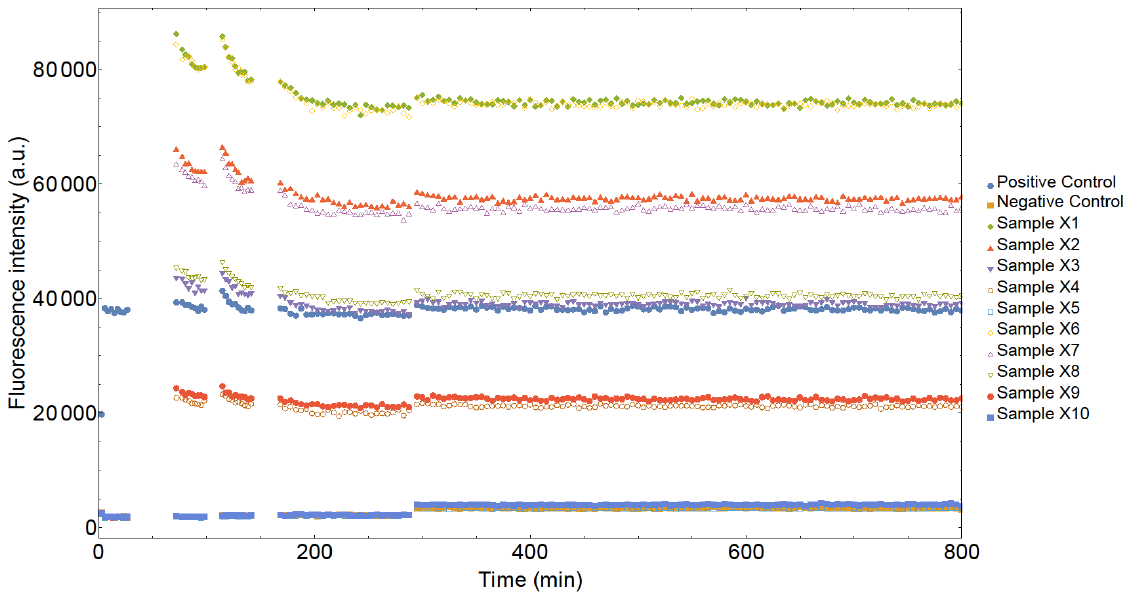

(a)

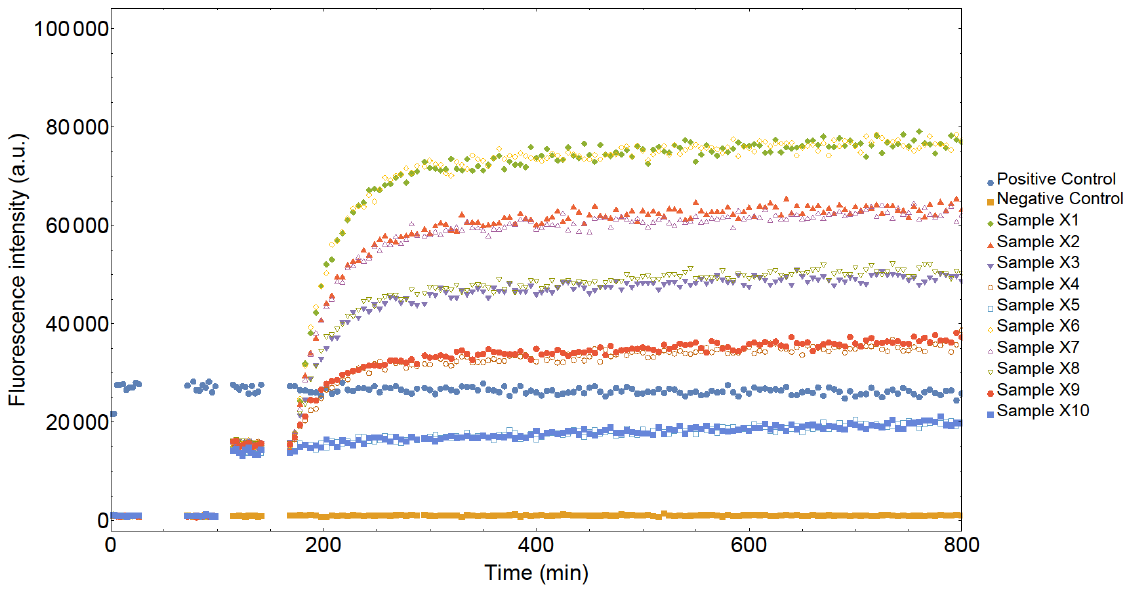

(b)

Figure S20: Fluorescent signal from the Cy3 channel (a) and the Alexa channel (b) obtained for the template recovery reaction for monomer M_1_ and proofreader P_8_

Samples *X1*-*X10* correspond to experiments with varying [*MT* *_total_*] performed in duplicate. Sample pairs {X1, X6; X2, X7; X3, X8; X4, X9; X5, X10} had intended [*MT* _total_] of {8.92nM; 6.77nM; 4.46nM; 2.35nM; 0.01nM}, respectively.

The first injection point (first jump in Cy3 channel) is the addition of monomer bound template, *MT* *,* which is varied across multiple experimental runs. The second injection point (first jump in Alexa channel) corresponds to the addition of the reporter quencher pair, *RQ*. Finally, the reaction starts near t=180 min at the third injection point where the proofreader is added (second jump in Alexa channel). All experiments were performed in duplicates.

After subtracting the negative control values, we take the average of the sample fluorescence values and normalize the fluorescence signal so that the reactions start at time t=0. For each sample in the Cy3 and Alexa channels, the third injection time is set at t=0 and the fluorescence value just before the point of injection is considered as that sample’s fluorescence value at t=0, see Figure S21.

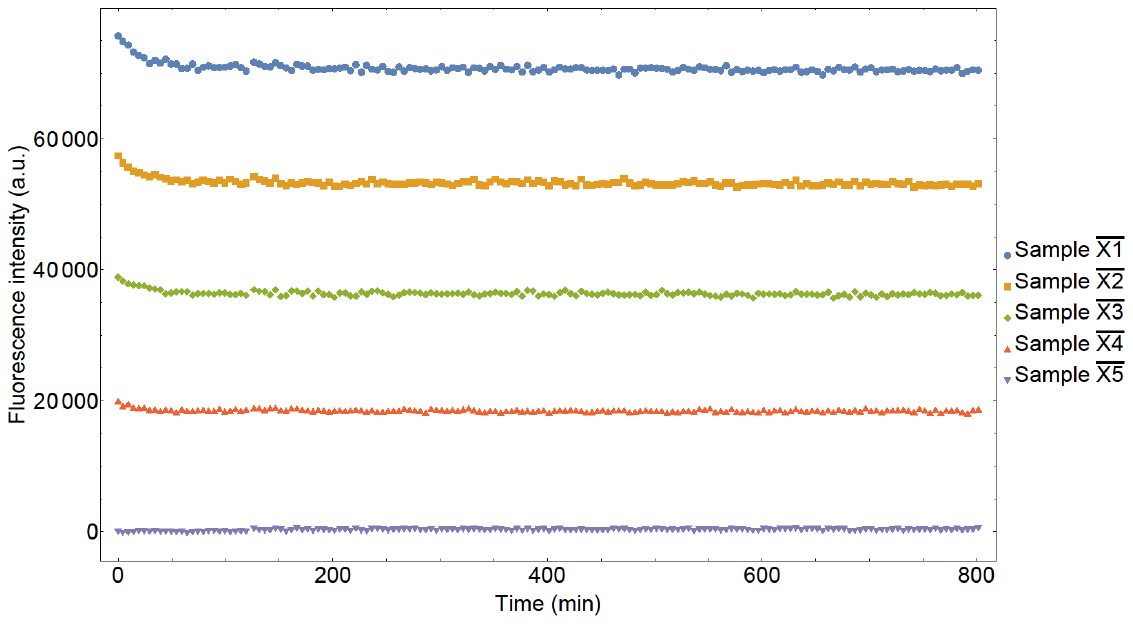

(a)

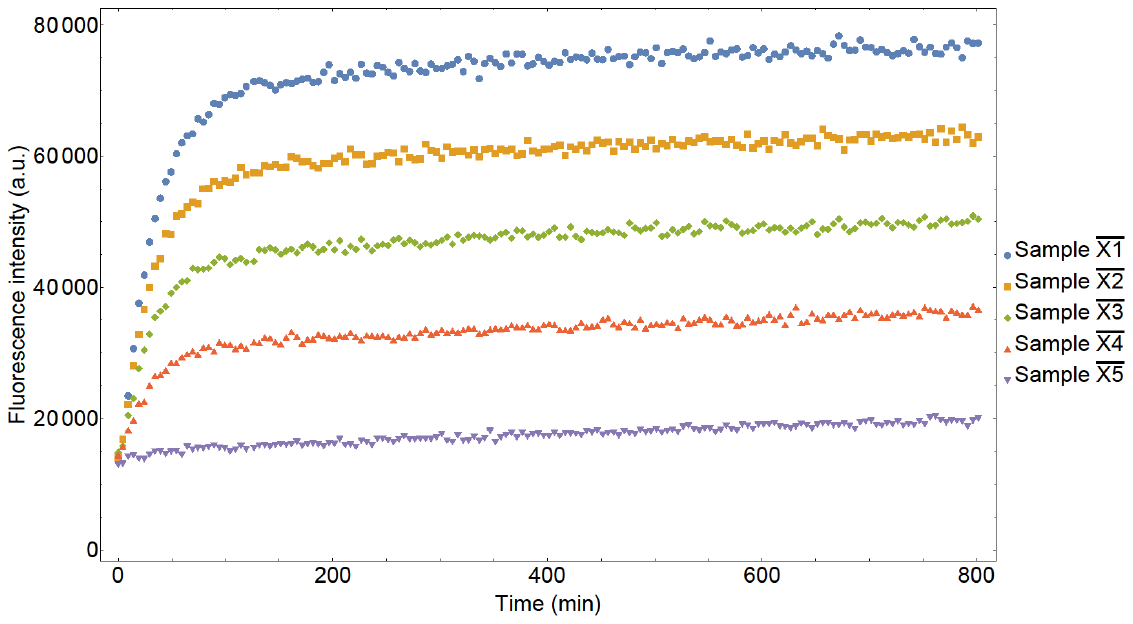

(b)

Figure S21: Normalized fluorescent signal from the Cy3 channel (a) and the Alexa channel (b) obtained for the template recovery of monomer M_1_ by proof-reader P_8_

The fluorescence in the Cy3 channel is due to *MT* , *MP* and *MPR,* but since the fluorescence signal density of each of these species is approximately the same, the signal generates a flat time series. These time series can be used to estimate the initial concentration of template-bound monomer, [*MT* *_total_*] by taking the time average of the fluorescent signal in the Cy3 channel after the final injection, and dividing the signal by the fluorescence signal density of *MPR* in the Cy3 channel, *F_MPR,C_*.

$[MT_{total}]=\frac{<f\left( t \right)_{C}>}{F_{MPR,C}}$ .	(S28)

From the Alexa channel, we can extract the concentration of reporter-bound complexes, *MPR* and *PR*. The total fluorescent signal in the Alexa channel is generated by *RQ*, *MPR* and *PR*:

${\left[ RQ \right]F}_{RQ,A}+\left[ MPR \right]F_{MPR,A}+\left[ PR \right]F_{PR,A}=f\left( t \right)$.	(S29)

Here *F_RQ,A_* is the fluorescence signal density of *RQ* in the Alexa channel. *F_MPR,A_* _­_and *F_PR,A_* are the fluorescence signal density of *MPR* and *PR* respectively in the Alexa channel and are considered equal (Supplementary note 17). Combining Eqs. (S25) and (S29) gives

$\left[ MPR \right]+[PR]=\frac{f\left( t \right)-(\left[ RQ \right]_{total})F_{RQ,A}}{F_{MPR,A}-F_{RQ,A}}$,	(S30)

where [*RQ_total_*] is obtained by saturating the solution with excess *M* to trigger all reporters, Supplementary note 5: Step 7. The final concentration of *MPR* is then determined by dividing the fluorescent signal with the fluorescence signal density of *MPR* in the Alexa channel. The concentration of *MPR* obtained at this point is equal to [*RQ_total_*]. Using this approach gives [*RQ_total_*] equal to 22.57nM for *M_1_P_8_*.

The normalized fluorescence signals from the Cy3 and Alexa channels are now converted to total monomer concentration and [*MPR*]+[*PR*] concentration respectively using Eq.(S30), Figure S22.

The values of [*MT* *_total_*] from extracted from Cy3 signal from Sample $\bar{X1}$ to $\bar{X5}$: {8.92nM, 6.77nM, 4.46nM, 2.35nM, 0.01nM}. The initial proofreader concentration, [*P_total_*] is assumed to be 50nM. The predicted {*k_3_*, *k_4_, k_7_*} rate constants, as has been discussed in Supplementary note 11, can be found in Table S11.

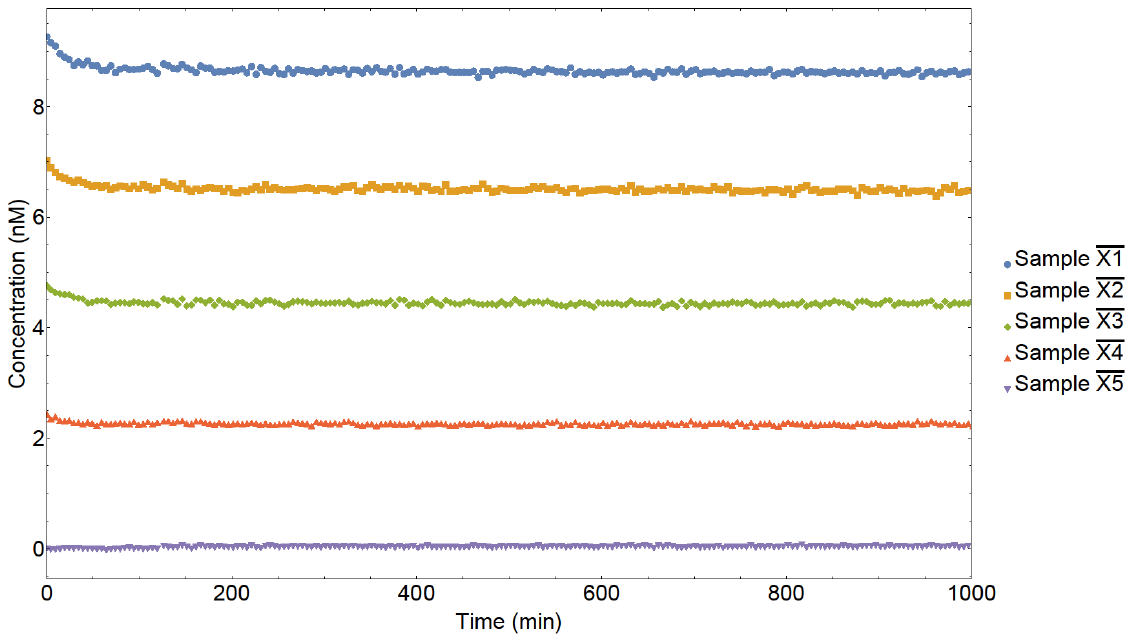

(a)

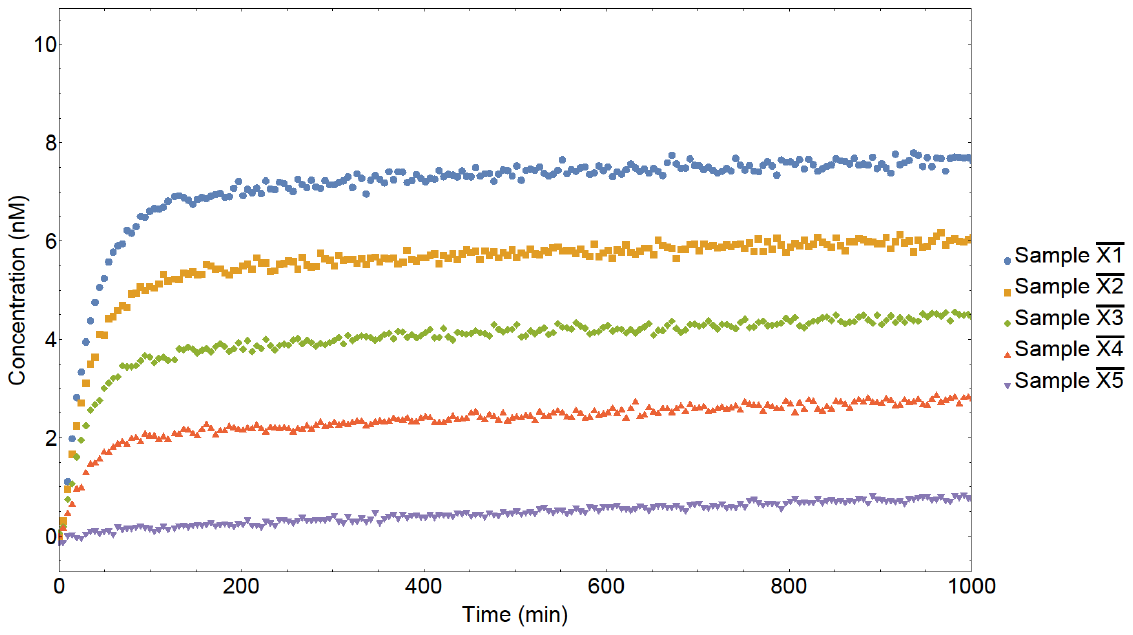

(b)

Figure S22: Total monomer concentration from the Cy3 channel (a) and [MPR]+[PR] time signal from the Alexa channel (b) obtained for the reporter reaction of M_1_P_8_

### Supplementary note 13: Fluorescence data processing and ODEs for discard pathway

The discard pathway consists of the following set of reactions:

**Lock opening**

$ML+T \begin{matrix} k_{1} \\ \leftrightharpoons\\ k_{2} \end{matrix} MT+L$,	(S31)

**Template recovery:**

$MT+P \begin{matrix} k_{3} \\ \leftrightharpoons\\ k_{4} \end{matrix} MP+T,$	(S32)

**MP-triggered reporter signal:**

$MP+RQ \begin{matrix} k_{5} \\ \leftrightharpoons\\ k_{6} \end{matrix} MPR+Q,$	(S33)

**Proofreader-triggered reporter signal (leak):**

$P+RQ \underset{\to}{k_{7}}PR+Q$.	(S34)

The ordinary differential equations (ODEs) that can completely describe the system are:

$\frac{d}{dt}[MT]= k_{1}[ML][T]-k_{2}[MT][L]-k_{3}[MT][P]+k_{4}[MP][T]$,	(S35)

$\frac{d}{dt}\left[ MP \right]= k_{3}\left[ MT \right][P]-k_{4}[MP][T]-k_{5}[MP][RQ]+k_{6}[MPR][Q]$,	(S36)

$\frac{d}{dt}\left[ RQ \right]= -k_{5}\left[ MP \right]\left[ RQ \right]+k_{6}\left[ MPR \right]\left[ Q \right]-k_{7}\left[ P \right]\left[ RQ \right]+k_{8}\left[ PR \right]\left[ Q \right],$	(S37)

$\frac{d}{dt}[PR]=k_{7}\left[ P \right][RQ]-k_{8}[PR][Q],$	(S38)

$\frac{d}{dt}\left[ L \right]=k_{1}\left[ ML \right]\left[ T \right]-k_{2}\left[ MT \right]\left[ L \right].$	(S39)

The conservation equations used are:

$\left[ ML \right]+\left[ MT \right]+\left[ MP \right]+\left[ MPR \right]=\left[ ML_{total} \right],$	(S40)

$\left[ RQ \right]+\left[ PR \right]+\left[ MPR \right]=\left[ RQ_{total} \right]$,	(S41)

$\left[ MP \right]+\left[ MPR \right]+\left[ PR \right]+\left[ P \right]=\left[ P_{total} \right],$	(S42)

$\left[ ML \right]+\left[ L \right]=1.2\left[ ML_{total} \right]$	,	(S43)

$\left[ MT \right]+\left[ T \right]=[T_{total}]$,	(S44)

$\left[ RQ \right]+\left[ Q \right]=1.2\left[ RQ_{total} \right].$	(S45)

As we will show below, there is some excess *MPR,* [*MPR_i_*], present in the solution when the discard pathway reaction starts. [*MPR_i_*] is subtracted from [*ML_total_*], [*P_total_*] and [*RQ_total_*] to give the initial concentrations of *ML*, *P* and *RQ* respectively as [*ML_total_*]-[*MPR_i_*], [*P_total_*]-[*MPR_i_*] and [*RQ_total_*]-[*MPR_i_*]. [*T_total_*] is the initial concentration of the template. There is also excess quencher *Q* and lock *L* present in the solution when the reaction starts. The conservation equations defined above can be used to extract their initial concentrations as 0.2[*RQ_total_*]+[*MPR_i_*] and 0.2[*ML_total_*]+[*MPR_i_*] respectively. All other species have 0nM initial concentration.

Reactions were performed by mixing *ML* (locked monomer), *T* (template), *P* (proofreader), with variable template concentration. The raw fluorescence intensities from the Cy3 and Alexa channels are shown in Figure S23 for monomer *M_1_* and proof-reader *P_8_*.

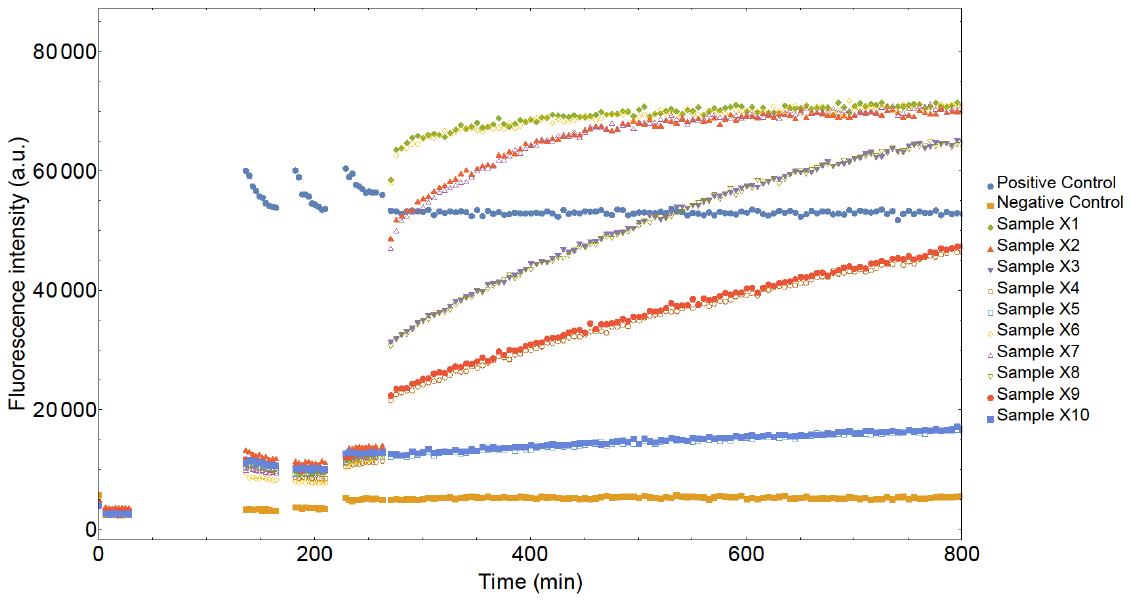

(a)

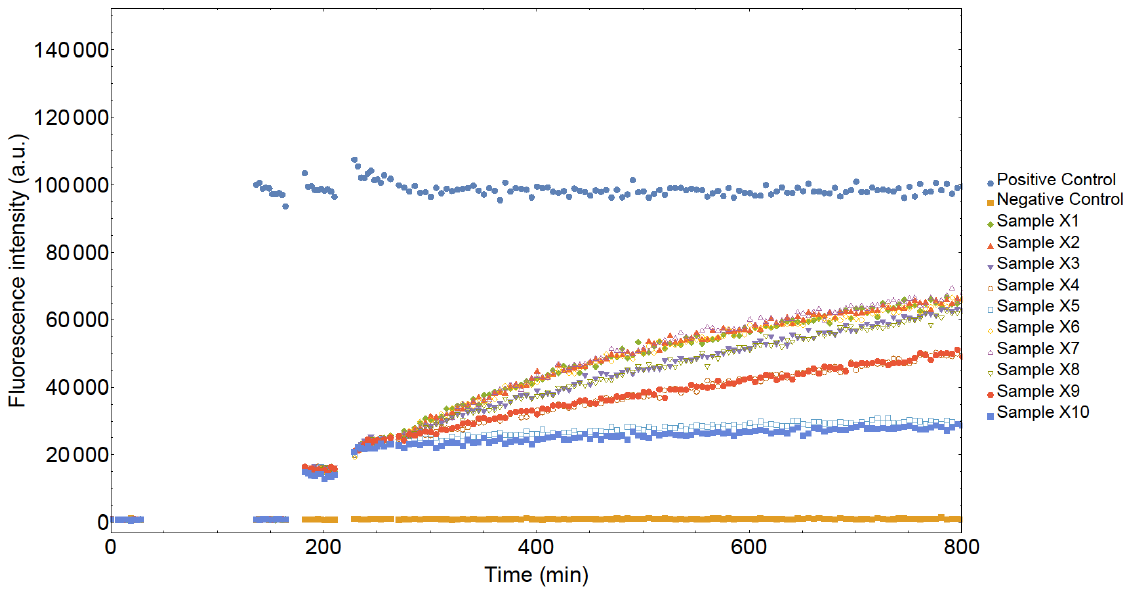

(b)

Figure S23: Fluorescent signal from the Cy3 channel (a) and the Alexa channel (b) for the template recovery reaction, where the monomer is M_1_ and proofreader is P_8_.

Samples X1-X10 correspond to experiments with varying [T_total_] performed in duplicate. Sample pairs {X1, X6; X2, X7; X3, X8; X4, X9; X5, X10} had intended [T_total_] of {6nM; 4nM; 2nM; 1nM; 0nM} respectively.

The first injection point (first jump in Cy3 channel) is the addition of the locked monomer, [*ML_total_*]. The second injection point (first jump in the Alexa channel) corresponds to the addition of the reporter quencher pair, *RQ* with concentration [*RQ_total_*]. The third injection point (second jump in Alexa channel) is the addition of proofreader with concentration [*P_total_*]. Finally, the reaction starts near t=180min at the fourth injection point where the template is added with concentration [*T_total_*]. All experiments were performed in duplicates.

After subtracting the negative control values, we take the average of the sample fluorescence values for repeat experiments and normalize the fluorescence signal so that the reactions start at time *t*=0. For each sample in the Cy3 and Alexa channels, the template injection time is set at *t*=0 and the fluorescence value just before the point of injection is considered as that sample’s fluorescence value at *t*=0.

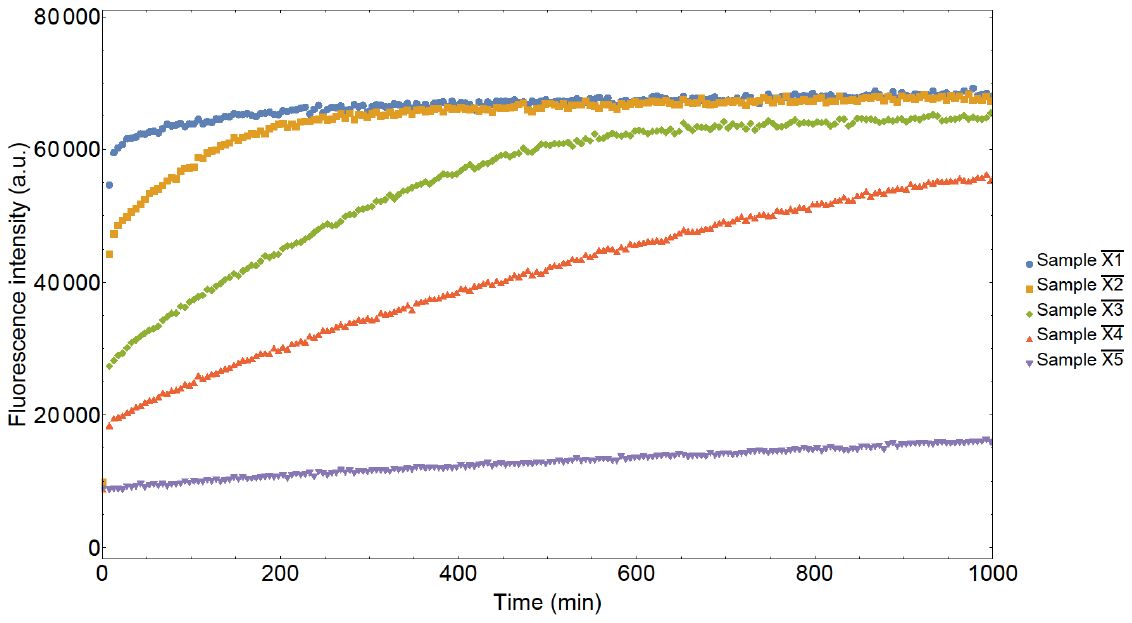

(a)

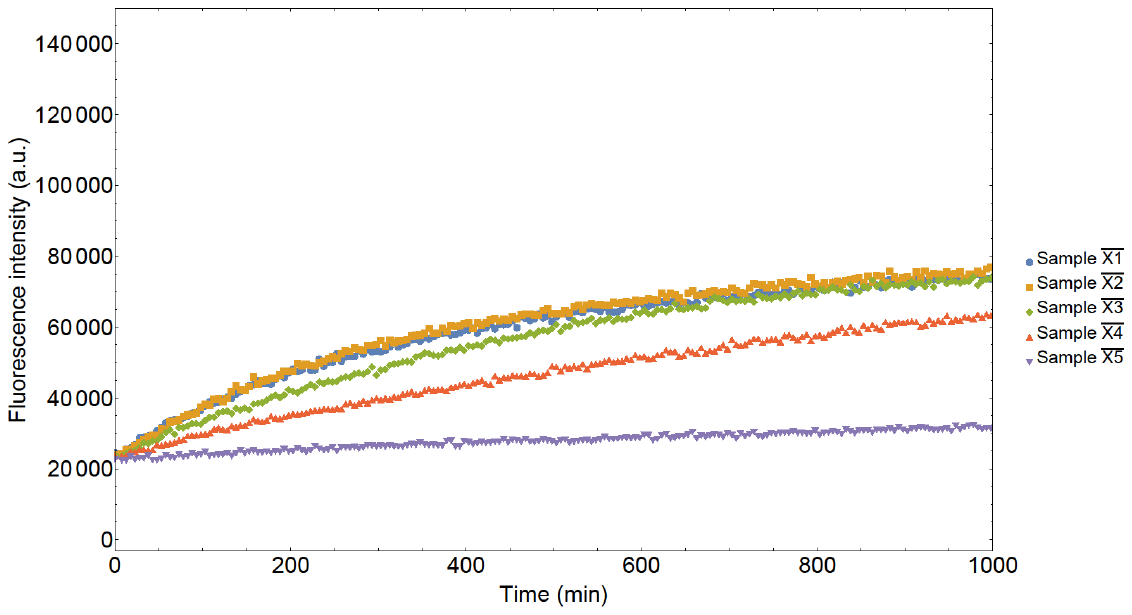

(b)

Figure S24: Normalized fluorescent signal from the Cy3 channel (a) and the Alexa channel (b) obtained for the discard pathway reaction with monomer M_1_ and proofreader P_8_

[*ML_total_*], the starting concentration of *ML*, was estimated by running the reaction for a long time in the Cy3 channel, Figure S25. At long times, all monomer complexes were unblocked, forming *MPR*. The concentration of *MPR* was estimated by taking the mean of the last 10 datapoints from the Cy3 channel (datapoints corresponding to reaction completion) and dividing by the signal density *F_MPR,C_*. This concentration is equal to [*ML_total_*] using Eq. (40). The experiment was then further saturated by addition of excess monomer to estimate [*RQ_total_*], as in Supplementary note 5, step 7. [*RQ_total_*] was estimated as 20.15nM for the case shown in Figure S24. The initial proofreader concentration [*P_total_*] was assumed to be 50nM. It is assumed that the concentration of the template, [*T_total_*], is as intended in the experimental setup {6nM, 4nM, 2nM, 1nM, 0nM}.

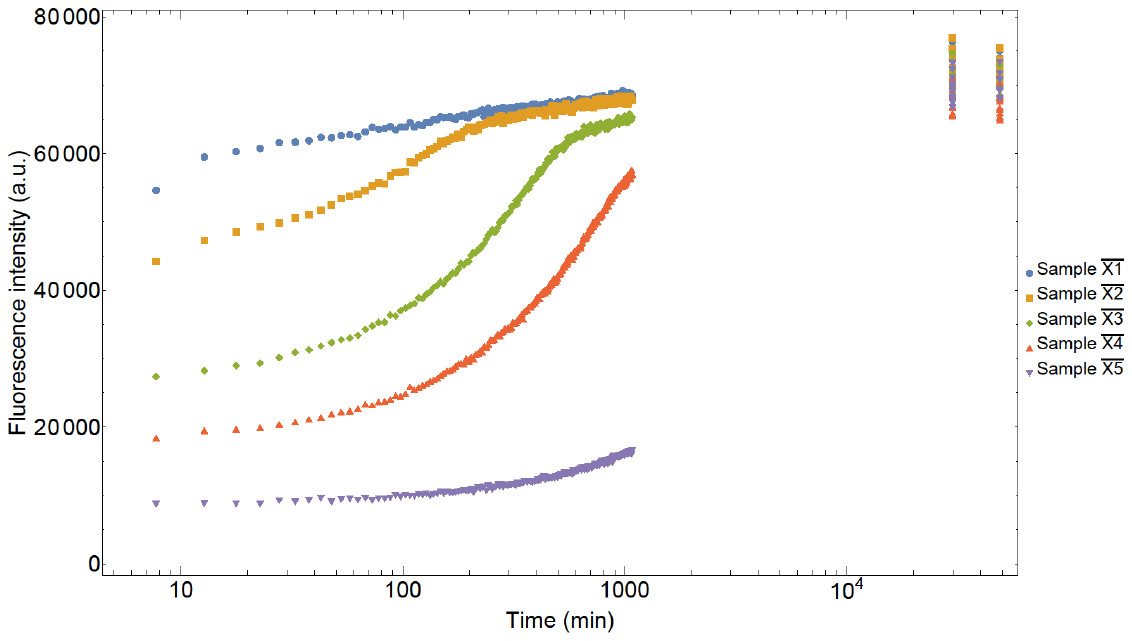

Figure S25: Normalized fluorescent signal from the Cy3 channel over a long time for the discard pathway reaction with monomer M_1_ and proof-reader P_8_

Next, we convert the fluorescent signal from the Alexa channel into triggered reporter [*MPR*]+[*PR*] concentration. This conversion can be done by the following protocols. In the Alexa channel, we know that the total fluorescent signal is generated by the 3 complexes: *RQ*, *MPR* and *PR*.

${\left[ RQ \right]F}_{RQ,A}+\left[ MPR \right]F_{MPR,A}+\left[ PR \right]F_{PR,A}=f\left( t \right)$	(S46)

Combining Eq.(S41) and (S46) gives

$\left[ MPR \right]+[PR]=\frac{F\left( t \right)-(\left[ RQ \right]_{total})F_{RQ,A}}{F_{MPR,A}-F_{RQ,A}}$	(S47)

The normalized fluorescence signals from the Cy3 and Alexa channels are now converted to total monomer concentration and [*MPR*]+[*PR*] concentration respectively, Figure S26.

Figure S 26: [MPR]+[PR] time signal from the Alexa channel obtained for the discard pathway reaction with monomer M1 and Proofreader P8

Note that there is an initial concentration of *MPR* recorded in the signal in Figure S26. This unexpected signal is attributed to the formation of *MPR* between the third and fourth injection points in Figure S23. This excess *MPR* concentration, [*MPR_i_*] is subtracted from [*RQ_total_*], [*P_total_*], [*ML_total_*] and the conservation equations are modified to reflect this. The predicted {*k_3_*, *k_4_, k_7_*} rate constants, as has been discussed in Supplementary note 11, can be found in Table S11.

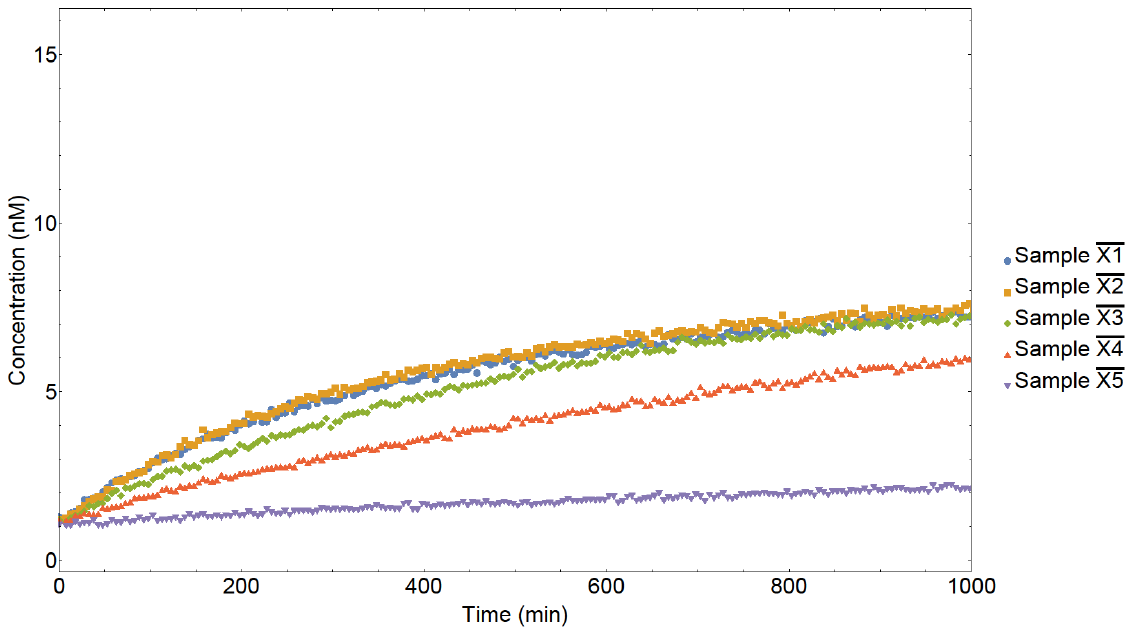

### Supplementary note 14: ODE for Dimerization reaction

The dimerization reaction is modelled by the following set of equations:

$ML+T \begin{matrix} k_{1} \\ \leftrightharpoons\\ k_{2} \end{matrix} MT+L$,	(S48)

$MT+P \begin{matrix} k_{3} \\ \leftrightharpoons\\ k_{4} \end{matrix} MP+T,$	(S49)

$MT+N \begin{matrix} k_{8} \\ \leftrightharpoons\\ k_{9} \end{matrix} MN+T$.	(S50)

Based on some preliminary experiments, *k_9_* ~ 0 The ODEs that represent the above set of equations completely are:

$\frac{d}{dt}[MT]= k_{1}[ML][T]-k_{2}[MT][L]-k_{3}[MT][P]+k_{4}[MP][T]-k_{8}[MT][N]+k_{9}[MN][T]$, (S51)

$\frac{d}{dt}\left[ MP \right]= k_{3}\left[ MT \right]\left[ P \right]-k_{4}[MP][T]$,	(S52)

$\frac{d}{dt}[MN]=k_{8}[MT][N]-k_{9}[MN][T]$.	(S53)

The conservation equations are:

$\left[ ML \right]+\left[ MT \right]+\left[ MP \right]+\left[ MN \right]=[ML_{total}]$,	(S54)

$\left[ N \right]+\left[ MN \right]=[N_{total}]$,	(S55)

$\left[ MP \right]+\left[ P \right]=[P_{total}]$,	(S56)

$\left[ ML \right]+\left[ L \right]=\left[ ML_{total} \right],$	(S57)

$\left[ MT \right]+\left[ T \right]=[T_{total}]$.	(S58)

The initial conditions at t=0 are: [*ML*]=[*ML_total_*]=8nM, [*T*]=[*T_total_*] is varied from {6nM, 4nM, 2nM, 1nM, 0nM}, [*N*] = [*N_total_*] = 10nM, [*P*] = [*P_total_*] = 50nM.

### Supplementary note 15: Dimerization via HMSD

From both the fluorescence signal f_c_(t) from Cy3 channel and f_a_(t) from AlexaFluor-647 channel the negative control (only buffer) signals are subtracted to generate the respective corrected signals f*_c_* and f*_a_*.

f_c_ is generated by the species *M’L’*, *M’T’*, *M’N*, *M’P’*, and *M’NR_1_*. From our experimental system, it is not possible to quantify the four fluorescent species *M’T’*, *M’N*, *M’P’*, and *M’NR_1_* separately. Therefore, for simplicity, we assume that all these species have the same fluorescence signal density, given by the calibration of *M’NR_1_* in Supplementary Figure S29-30.

Similarly, f_a_ is generated by *R_1_Q_1_*, *M’NR_1_*, and *NR_1_*. Since the local environment of fluorophore in *M’NR_1_* and *NR_1_* are same, we assume the fluorescence caused by unit concentration of *M’NR_1_* and *NR_1_* to be the same.

At any given time, the fluorescence signals can then be estimated by the following:

$fc=\left[ {M'NR}_{1} \right]F_{{M^{'}NR}_{1},C}+\left( 8\mathrm{nM}-\left[ {M'NR}_{1} \right] \right)F_{M'L'}$	(S59)

$fa=\left[ {M'NR}_{1} \right]F_{{M^{'}NR}_{1},A}+\left( 20\mathrm{nM}-\left[ {M'NR}_{1} \right] \right)F_{RQ}$	(S60)

Where 8nM is the initial concentration of *M’L’* and 20nM is the initial concentration of RQ. F_M’NR,C_ and F_M’L’_ are the Cy3 fluorescence signal densities of *M’NR_1_* and *M’L’*. F_M’NR,A_ and F_RQ_ are the AlexaFluor-647 fluorescence signal densities of *M’NR_1_* and *R_1_Q_1_* (Supplementary Figure S29-30).

### Supplementary note 16: Fluorescence data processing for SNP

From the fluorescence signal f_c_(t) from Cy3 channel, the negative control (only buffer) signals are subtracted to generate the corrected signal f*_c_*.

fc is generated by *RQ, TS-Probe-R*, and *Probe-R*. Since the local environment of fluorophore in *TS-Probe-R* and *Probe-R* are same, we assume the fluorescence caused by unit concentration of *TS-Probe-R* and *Probe-R* to be the same.

At any given time, we assume that the fluorescence signals are given by

$fc=\left[ WPR \right]F_{WPR}+\left( 20\mathrm{nM}-\left[ WPR \right] \right)F_{RQsnp}$	(S62)

Where 20nM is the initial concentration of RQ_snp_. F_WPR_ and F_RQsnp’_ are the Cy3 fluorescence signal densities for *TS-Probe-R_snp_* and *RQ_snp_* complexes (Supplementary Figure S31).

### Supplementary note 17: Protocol for generating calibration curves

The calibration curves were produced for complexes to convert the fluorescence data into concentration values. For this purpose, different DNA strands were annealed keeping the following considerations in mind:

a. The stocks’ concentrations were checked on the day using nanodrop.
b. The non-fluorescent/unlabelled strands were used in 20% excess to ensure that all the fluorescent strands are completely quenched or in duplexes with other strands to minimise any undesired signal from non-annealed complexes.
c. 100 nM or 200 nM stocks (as per the fluorescent strand concentrations) were prepared for each complex.
d. Fluorescence signals were measured over several hours to account for any changes in the fluorescence intensities.
e. A few initial data points were rejected since they show temperature-related decay while switching from laboratory ambient temperature (22-23°C) to experimental temperature in the plate-reader (25°C).
f. The average of fluorescence intensity data points, excluding these initial timepoints, was then plotted against concentration. A linear fit was used to extract fluorescence signal density for different complexes.

### Supplementary Figure 27: Cy3 calibration curves for *M_1_*‑containing species.

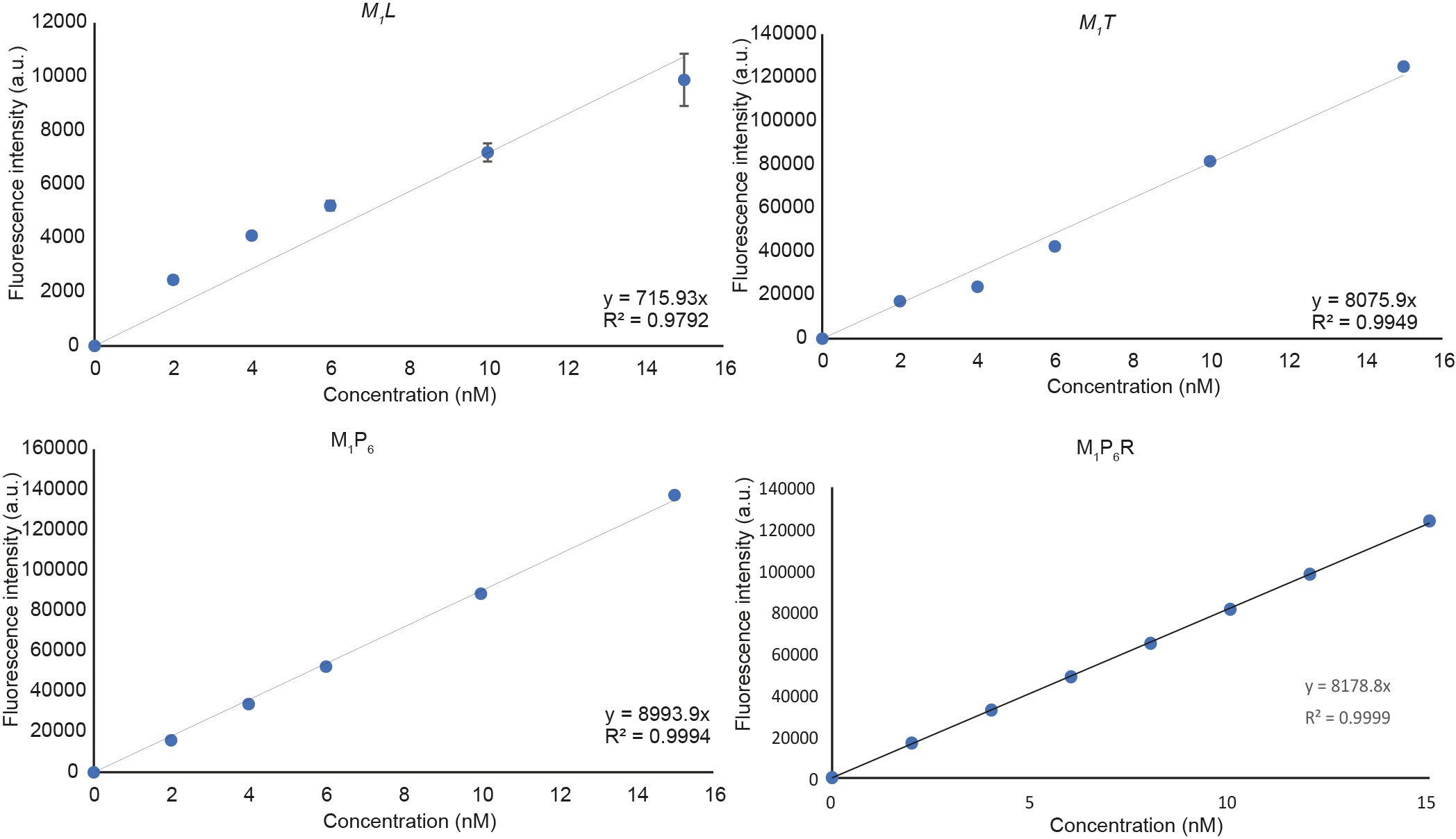

Figure S27: Cy3 calibration curve for complexes containing M_1_

100-200 nM *ML*, *MT* , *MP*, and *MPR* were pre-annealed. Different dilutions of these complexes were measured to record the Cy3 fluorescence intensities. The local environments of the fluorophore remain the same for all monomers and hence *M_1_* is used in all cases. Fluorescence intensities were plotted against concentrations to obtain the calibration curves and the fluorescence signal densities F_ML_, F*MT*, F_MP,C_, and F_MPR,C_.

### Supplementary Figure 28: AlexaFluor-647 calibration curves for triggered and quenched reporter for *M_1_P_6_*

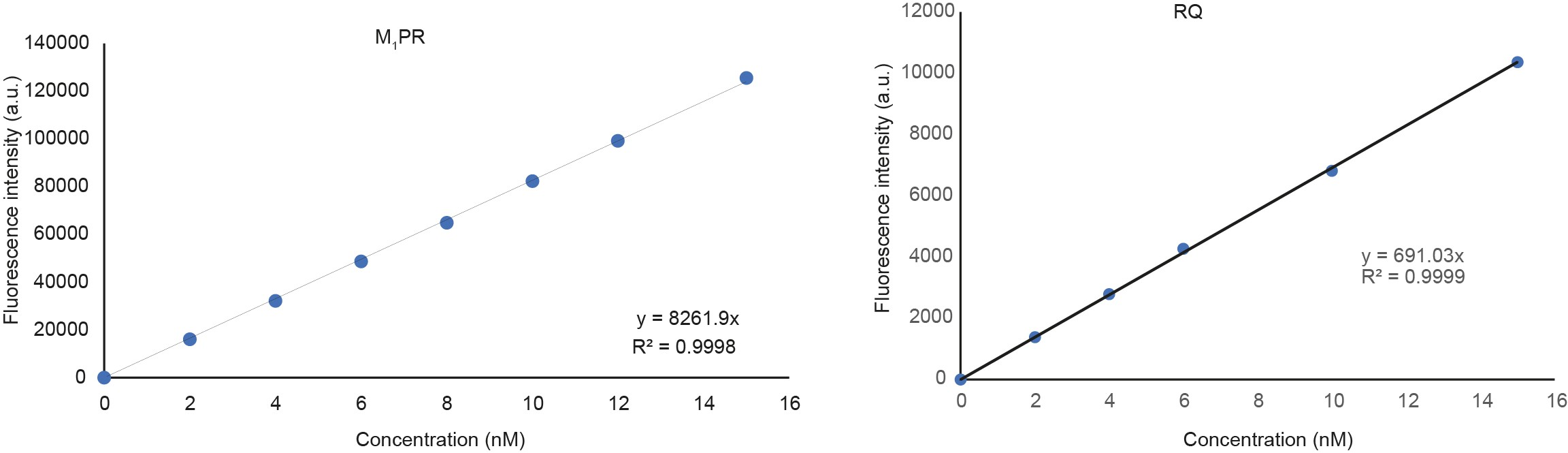

Figure S28: AlexaFluor-647 calibration curves for quenched and fluorescent reporters for M_1_P_6_

100-200 nM *M_1_PR* and *RQ* complexes were prepared. Different dilutions of those were measured to record the AlexaFluor-647 fluorescence intensities. Fluorescence intensities were plotted against concentrations to obtain the calibration curves and the fluorescence signal densities F_MPR,A_ and F_RQ._

### Supplementary Figure 29: Cy3 calibration curves for *M’_1_*‑containing species

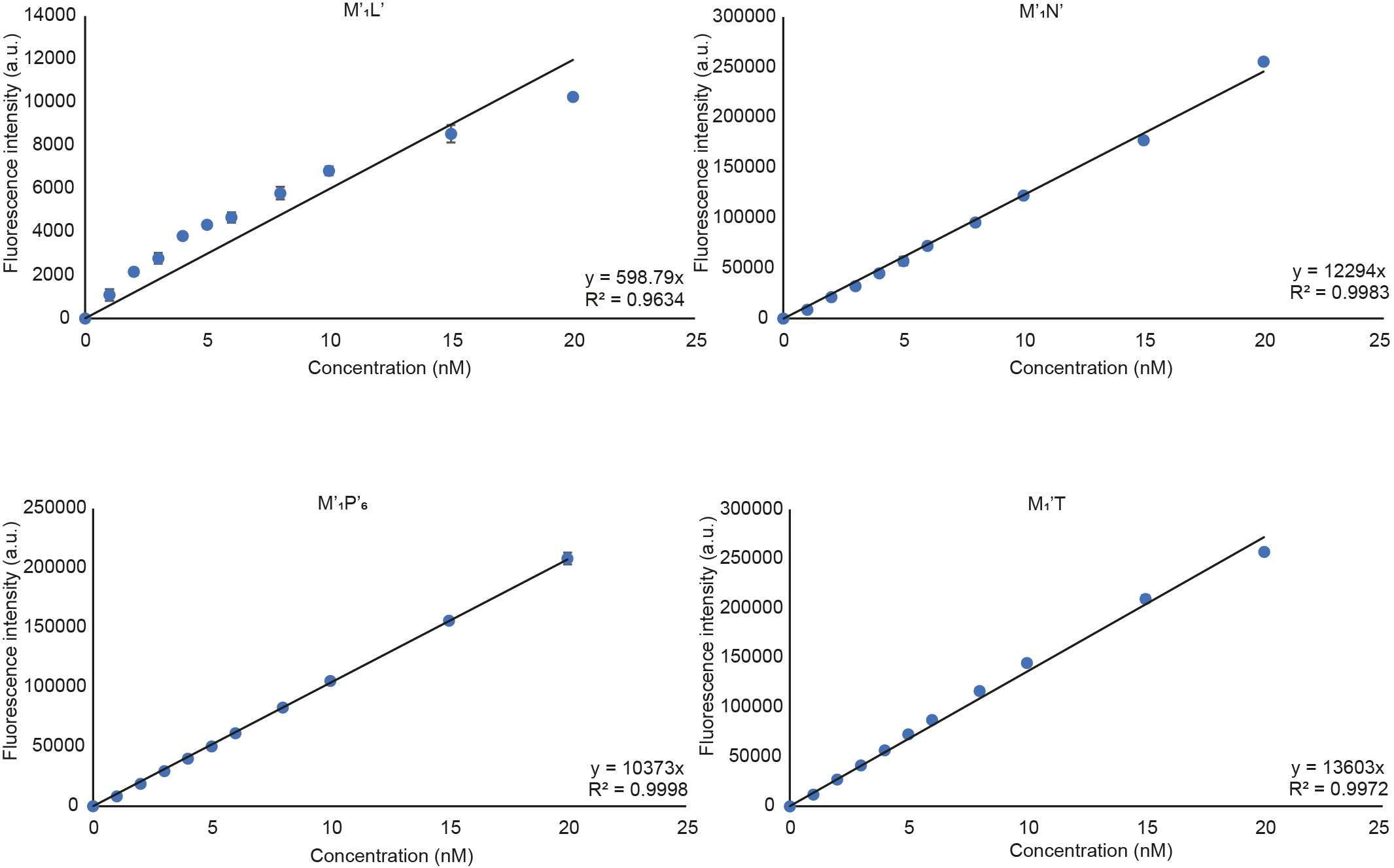

Figure S29: Cy3 calibration curves for complexes containing M_1_’

100-200 nM *M’L’, M’N, M’P’,* and *M’T’* were pre-annealed. Different dilutions of these complexes were measured to record the Cy3 fluorescence intensities. The local environments of the fluorophore remain the same for all monomers and hence *M’*_1_ is used in all cases. Fluorescence intensities were plotted against concentrations to obtain the calibration curves and the fluorescence signal densities F_M’L’,C_ F_M’N,C_ F_M’P’,C_ F_M’_*_T’_* _,C_

### Supplementary Figure 30: calibration curves for quenched and triggered reporters for *M’_1_N*

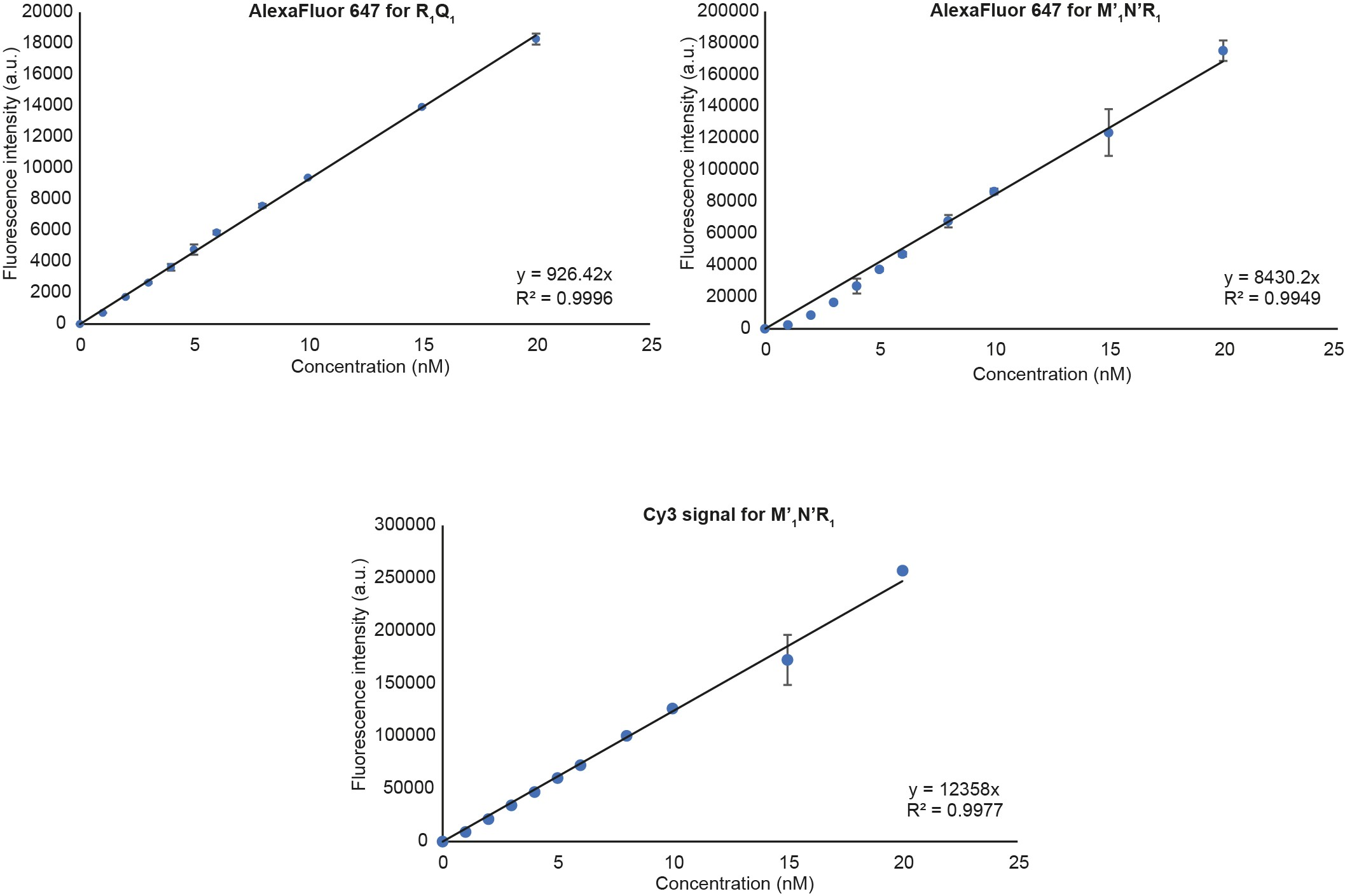

Figure S30: Calibration curves for quenched and fluorescent reporters for M_1_’N

100-200 nM *M_1_NR* and R_1_Q_1_ complexes were prepared. Different dilutions of those were measured to record the Cy3 and AlexaFluor-647 fluorescence intensities. Fluorescence intensities were plotted against concentrations to obtain the calibration curves and the fluorescence signal densities F_R1Q1,A_, F_M’NR,A_, and F_M’NR1,C_.

### Supplementary Figure 31: calibration curves for quenched and triggered reporter for SNP detection

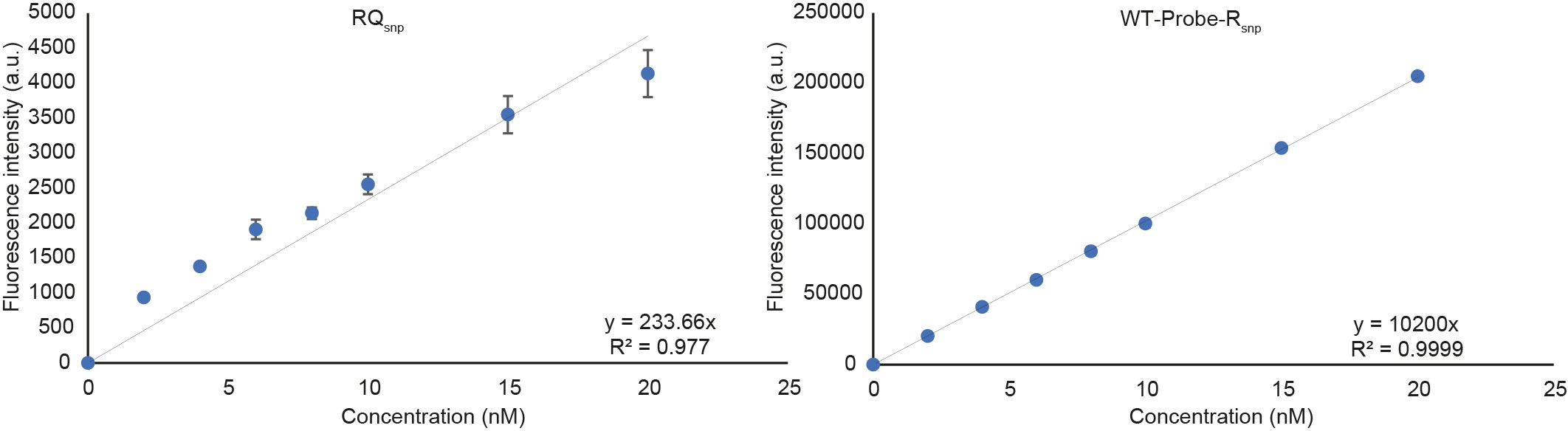

Figure S31: Cy3 calibration curves for quenched and fluorescent reporter complexes for SNP detection

100-200 nM *TS-Probe-R_snp_* and *RQ_snp_* complexes were prepared. Different dilutions of those were measured to record the Cy3 fluorescence intensities. Fluorescence intensities were plotted against concentrations to obtain the calibration curves and the fluorescence signal densities F_RQsnp_ and F_WPR_.

### Supplementary Figure 32: Binding of blocked monomers *M_1_L*, *M_2_L*, and *M_3_L* to *T*

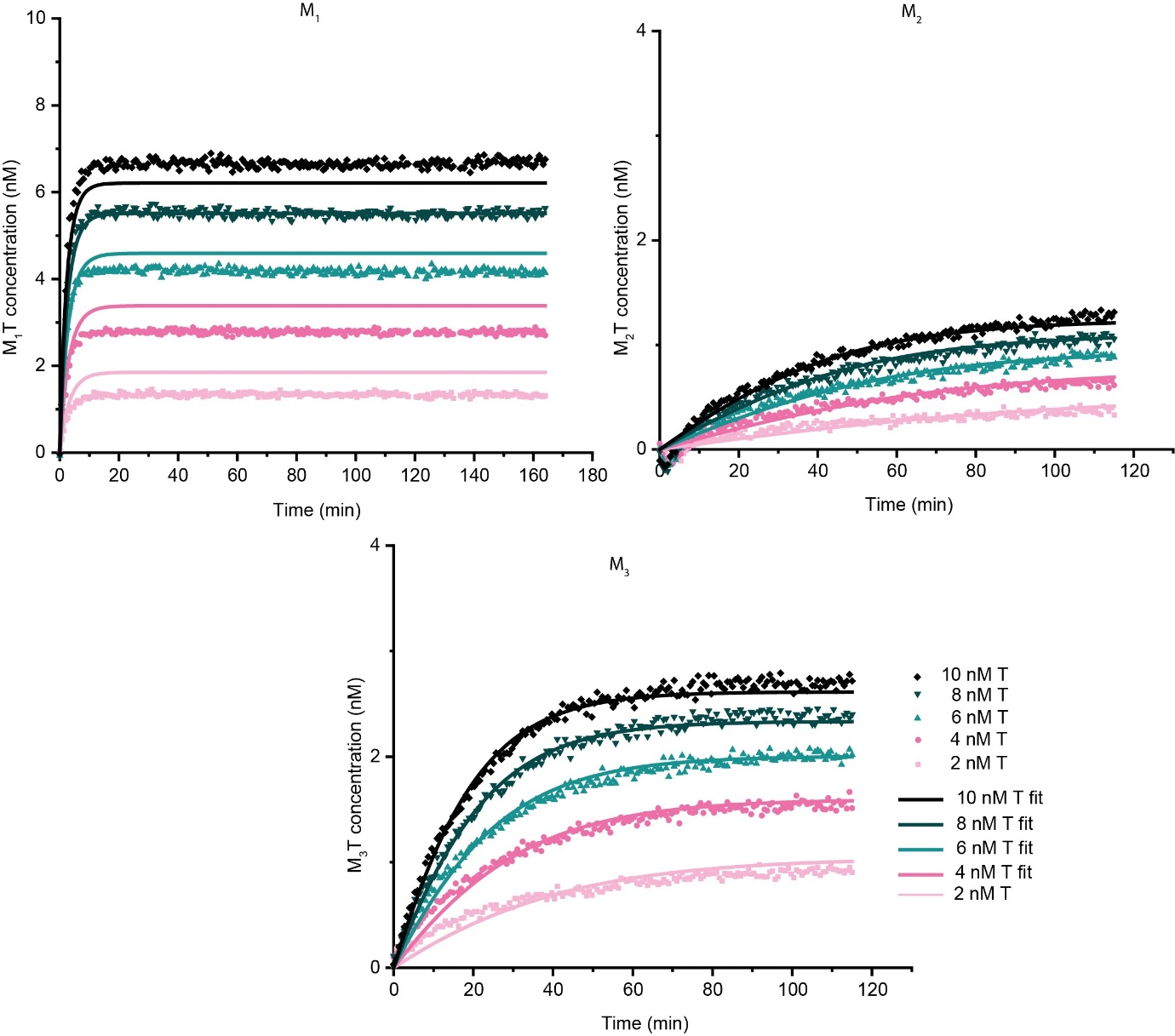

Figure S32: Template binding to ML complexes

4-10 nM *T* is added to 10 nM pre-annealed *ML* complexes. *T* invades *M_1_L* complex the fastest due to the perfectly matching toehold. *M_2_L* and *M_3_L* are invaded at slower rates due to the mismatches in the toehold domains. These data are the result of processing raw data as outlined in Supplementary note 10 and were used to fit the parameters *k_1_* and *k_2_* for each monomer given in Eq. (S1). The parameter values can be found in Table S9. The resultant fits are shown as solid lines.

Table S9: rate constants for the template binding process

| **Monomer** | **k_1_ M-^1^ sec^-1^** | **k_2_ M^-1^ sec^-1^** |
| --- | --- | --- |
| *M1* | 4.49E+05 | 1.36E+05 |
| *M2* | 5.38E+03 | 9.19E+04 |
| *M3* | 2.40E+04 | 9.75E+04 |

The parameters were obtained by fitting the ODE model to experimental data shown in Figure S32.The procedure is reported in Supplementary note 10.

### Supplementary Figure 33: displacement of the template from the monomer by *L*

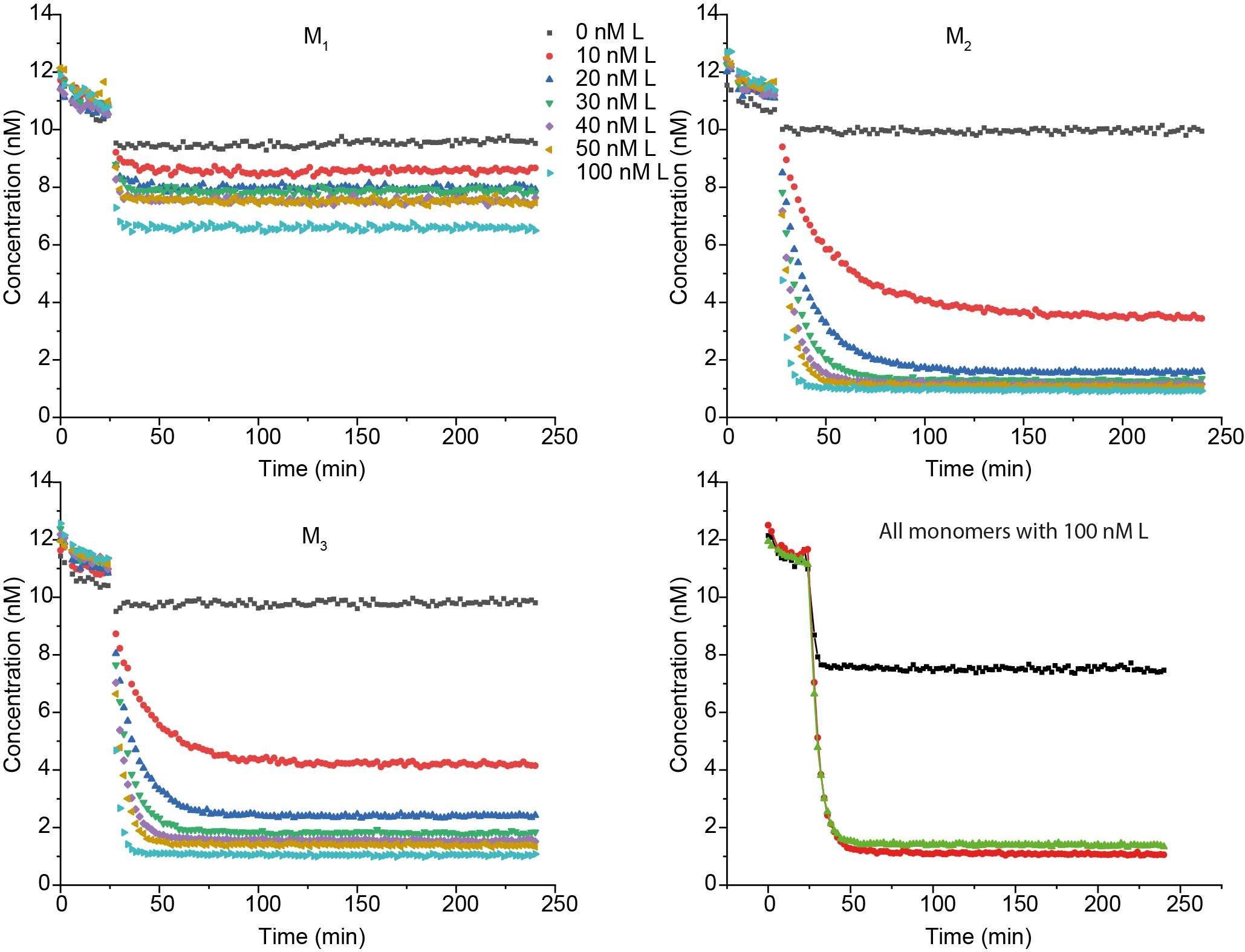

Figure S33: L binding to MT complexes

10-100 nM *L* is Injected to 10 nM pre-annealed *MT* complexes. Invasion of *M_1_T* by *L* is highly thermodynamically disfavoured and exhibits only a limited decay in fluorescence signal even after injecting 100 nM *L*. Although the equilibrium levels were different for different *M*, the time taken to reach the equilibrium were similar and no rate discrimination is seen. These data are the result of processing raw data as outlined in Supplementary note 10.

Table S10: Rate constants for the reporter binding of the MP complexes (best fit)

| **Monomer** | **Proofreader** | **k_5_ M^-1^ sec^-1^** | **k_6_ M^-1^ sec^-1^** |
| --- | --- | --- | --- |
| *M_1_* | *P_6_* | 3.78E+04 | 6.84E+03 |
|  | *P_7_* | 1.80E+05 | 3.28E+04 |
|  | *P_8_* | 1.48E+05 | 2.92E+04 |
| *M_2_* | *P_6_* | 2.43E+05 | 8.67E+04 |
|  | *P_7_* | 2.27E+05 | 5.47E+04 |
|  | *P_8_* | 2.92E+05 | 6.15E+04 |
| *M_3_* | *P_6_* | 2.14E+05 | 6.65E+04 |
|  | *P_7_* | 8.65E+04 | 3.16E+04 |
|  | *P_8_* | 3.34E+04 | 1.12E+04 |

In this fitting procedure, multiple possible initial monomer concentrations were tested, as outlined in Supplementary note 11. The results presented here are for the initial concentration assumption that allows the best fit of the reporter reaction, template recovery and the discard pathway process. The values of *k_5_, k_6_* are obtained alongside *k_3_, k_4_, k_7_* while using previously calculated values of *k_1_, k_2_*. All parameter values are given in Table S9-S11.

### Supplementary Figure 34: Kinetics of reporters for *M_1_P* complexes

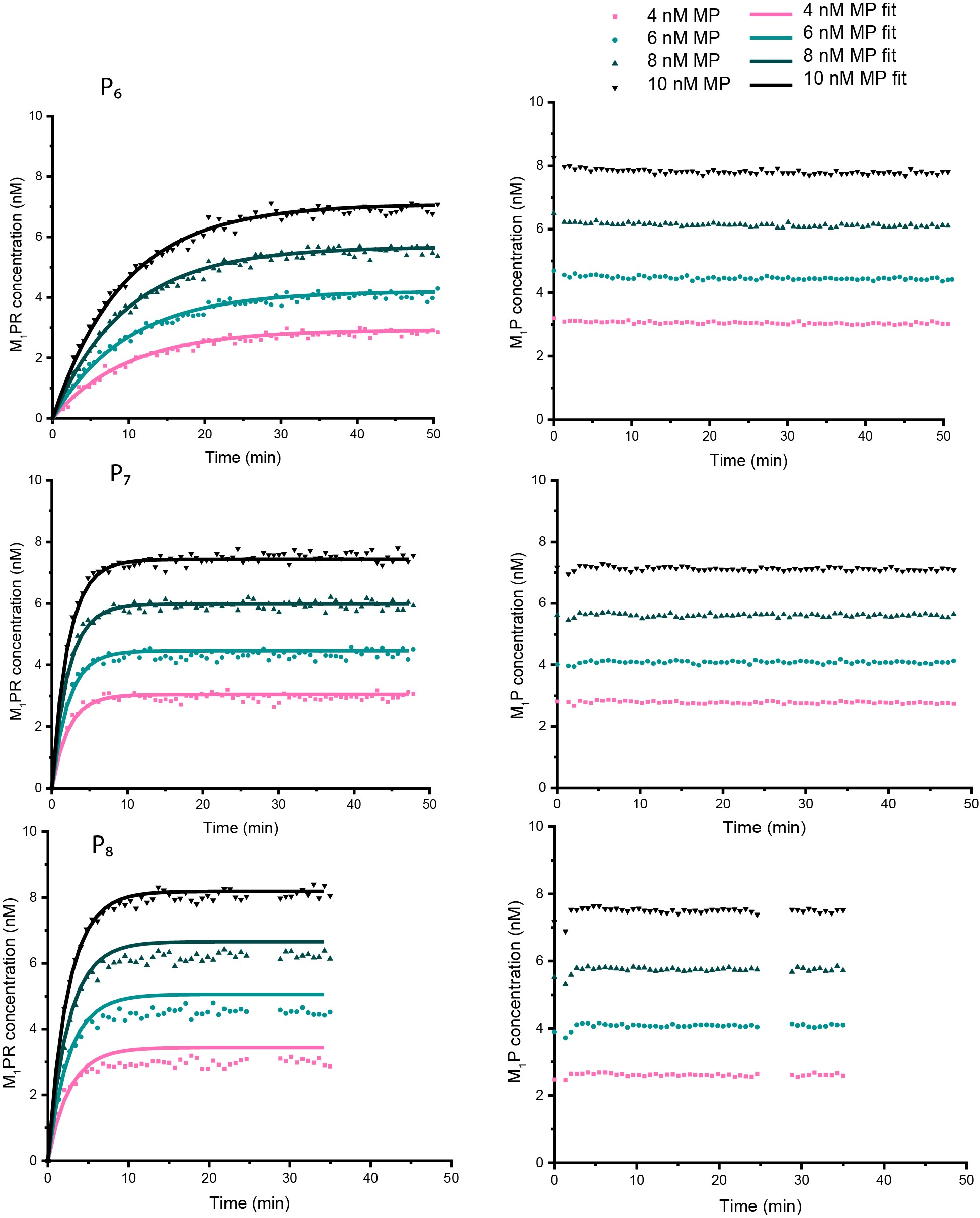

Figure S34: Reporter characterisation for M_1_P complexes

4-10 nM of pre-annealed *M_1_P* complexes were added to 40 nM of corresponding reporters, and fluorescence signals were measured in both AlexaFluor-647 (left column) and Cy3 (right column) channels. AlexaFluor-647 reading showed the progress of the experiment and Cy3 measurements were used for estimating the actual amount of added *M_1_P* complexes. Three proofreaders were tested; Top: *P_6_*, middle: *P_7_*, bottom: *P_8_*. Filled coloured symbols are experimentally obtained data. These data are the result of processing raw data as outlined in Supplementary note 11 and was used to fit the parameters *k_5_* and *k_6_* for each monomer (Table S10). Note that k_5_, k_6_ were not obtained independently and their calculation has been described in detail in Supplementary note 12. The values of *k_5_, k_6_* are obtained alongside *k_3_, k_4_, k_7_* while using previously calculated values of *k_1_, k_2_*. All parameter values are given in Table S9-S11. The resultant fits are shown as solid lines.

### Supplementary Figure 35: Kinetics of reporters for *M_2_P* complexes

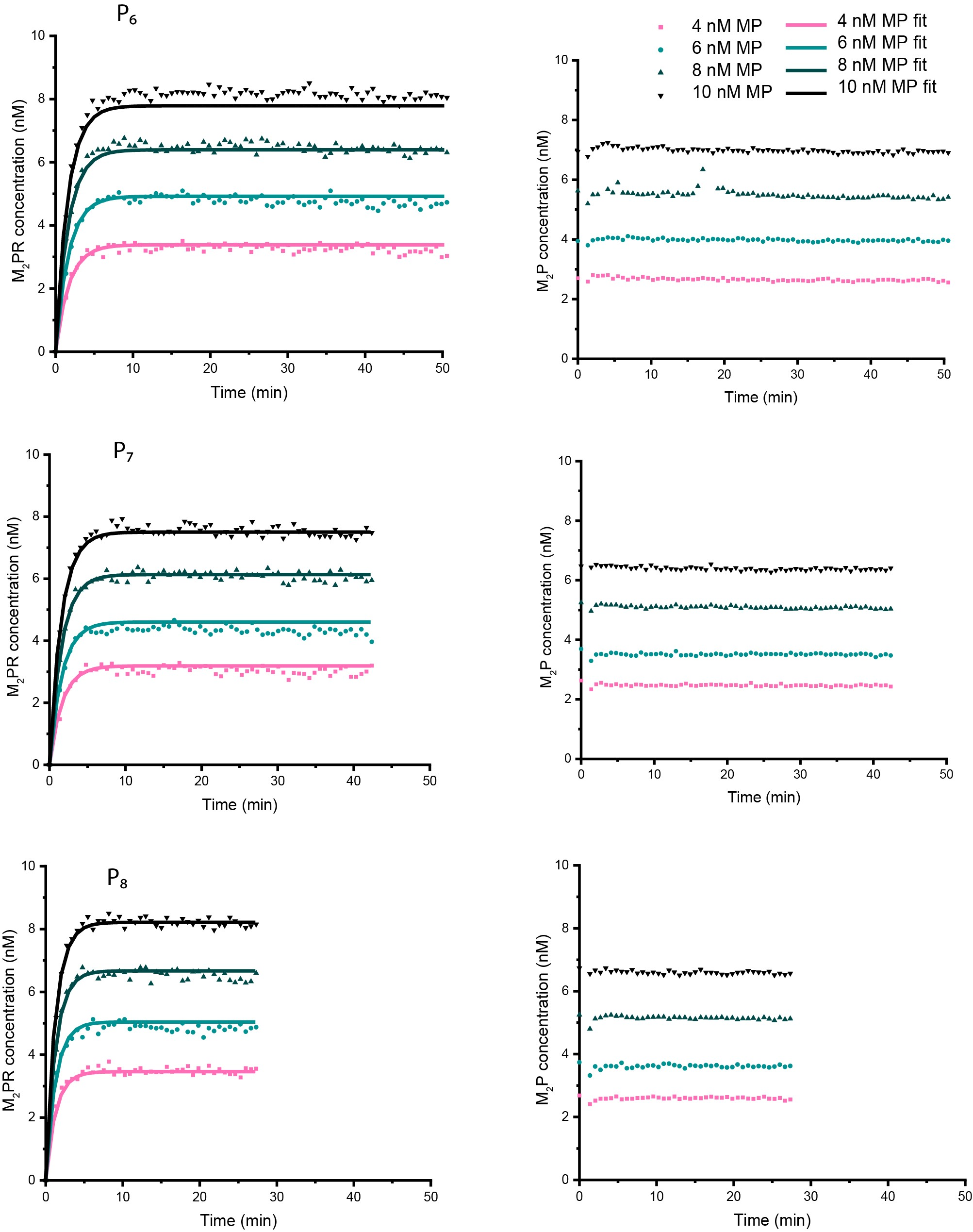

Figure S35: Reporter characterisation for M2P complexes

4-10 nM of pre-annealed *M_2_P* complexes were added to 40 nM of corresponding reporters, and fluorescence signals were measured in both AlexaFluor-647 (left column) and Cy3 (right column) channels. AlexaFluor-647 reading showed the progress of the experiment and Cy3 measurements were used for estimating the actual amount of added *M_2_P* complexes. Three proofreaders were tested; Top: *P_6_*, middle: *P_7_*, bottom: *P_8._* These data are the result of processing raw data as outlined in Supplementary note 11 and was used to fit the parameters *k_5_* and *k_6_* for each monomer (Table S10). Note that *k_5_, k_6_* were not obtained independently and their calculation has been described in detail in Supplementary note 12. The values of *k_5_, k_6_* are obtained alongside *k_3_, k_4_, k_7_* while using previously calculated values of *k_1_, k_2_*. All parameter values are given in Table S9-S11. The resultant fits are shown as solid lines.

### Supplementary Figure 36: Kinetics of reporters for *M_3_P* complexes

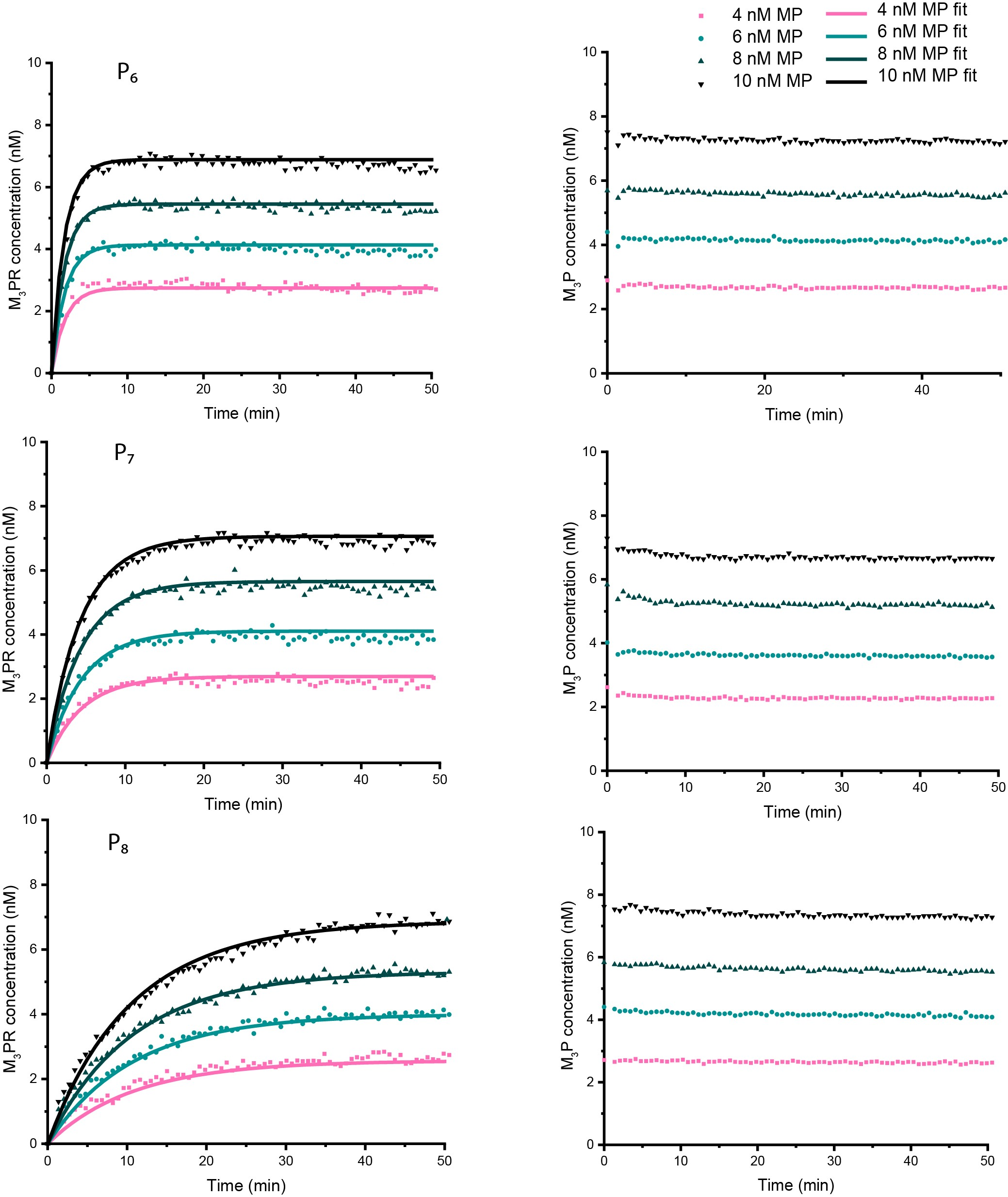

Figure S36: Reporter characterisation for M_3_P complexes

4-10 nM of pre-annealed *M_3_P* complexes were added to 40 nM of corresponding reporters, and fluorescence signals were measured in both AlexaFluor-647 (left column) and Cy3 (right column) channels. AlexaFluor-647 reading showed the progress of the experiment and Cy3 measurements were used for estimating the actual amount of added *M_3_P* complexes. Three proofreaders were tested; Top: *P_6_*, middle: *P_7_*, bottom: *P_._* These data are the result of processing raw data as outlined in Supplementary note 11 and was used to fit the parameters *k_5_* and *k_6_* for each monomer (Table S10). Note that *k_5_, k_6_* were not obtained independently and their calculation has been described in detail in Supplementary note 12. The values of *k_5_, k_6_* are obtained alongside *k_3_, k_4_, k_7_* while using previously calculated values of *k_1_, k_2_*. All parameter values are given in Table S9-S11. The resultant fits are shown as solid lines.

### Supplementary Figure 37: Template recovery for *M_1_T* complex

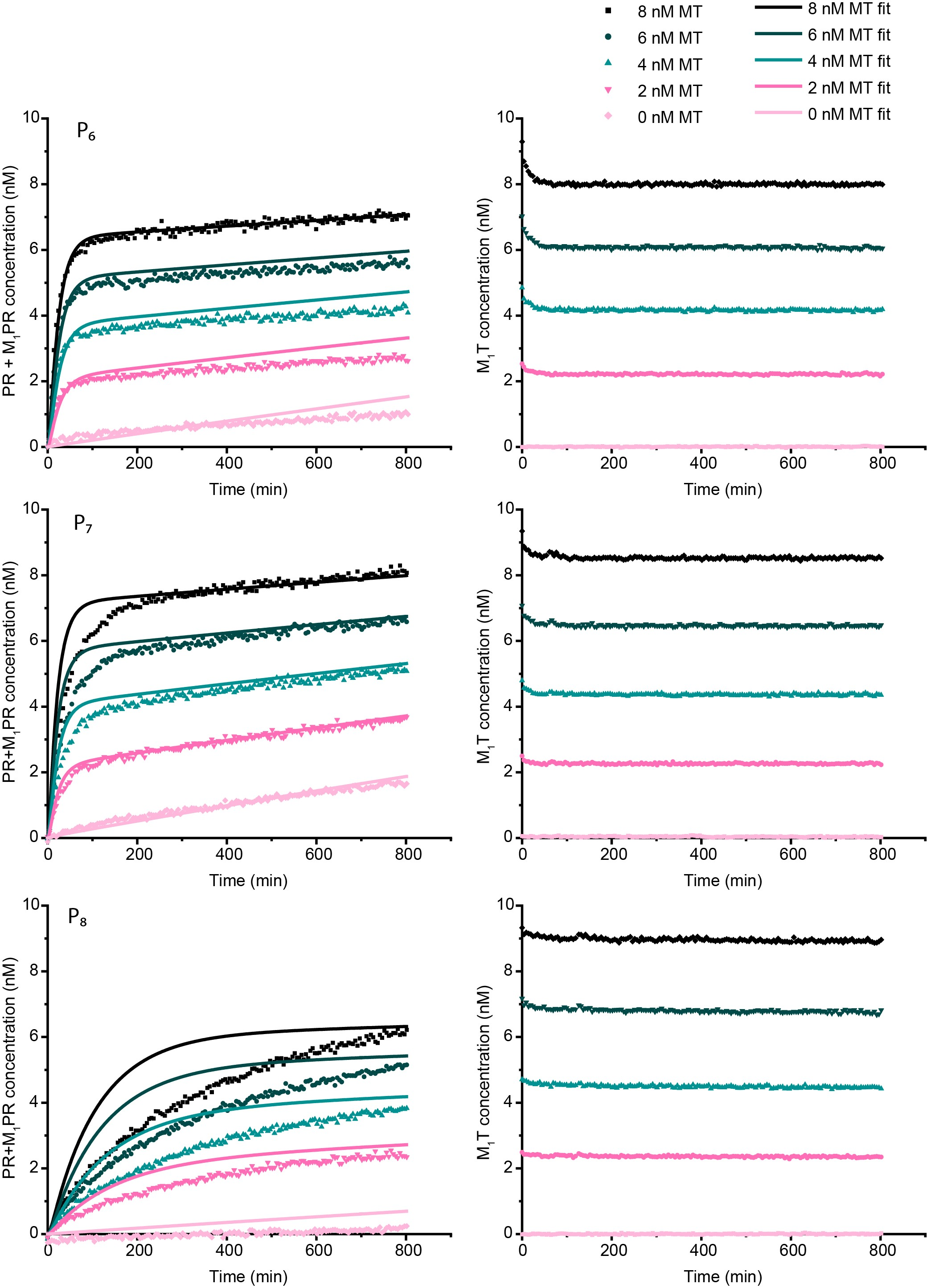

Figure S37: Template displacement by the proofreaders from M_1_T complex

50 nM of proofreader strands were added to a solution containing 20 nM *RQ* complex and 0-8 nM of *M_1_T* complex, and fluorescence signals were measured in both AlexaFluor-647 (left column) and Cy3 (right column) channels. AlexaFluor-647 reading showed the production of *M_1_PR* complexes and Cy3 measurements were used for estimating the exact concentration of added *M_1_T* complex. Three proofreaders were tested; Top: *P_6_*, middle: *P_7_*, bottom: *P_8._* Filled coloured symbols are experimentally obtained data processed as described in Supplementary note 12. Solid lines are the fits for these data. Note that *k_3_, k_4_* (forward and backward proofreader rates) were not obtained independently and their calculation has been described in detail in Supplementary note 12. The values of *k_3_, k_4_* are obtained alongside *k_5_, k_6_, k_7_* while using previously calculated values of *k_1_, k_2_*. All parameter values are given in Table S9-S11.

### Supplementary Figure 38: Template recovery for *M_2_T* complex

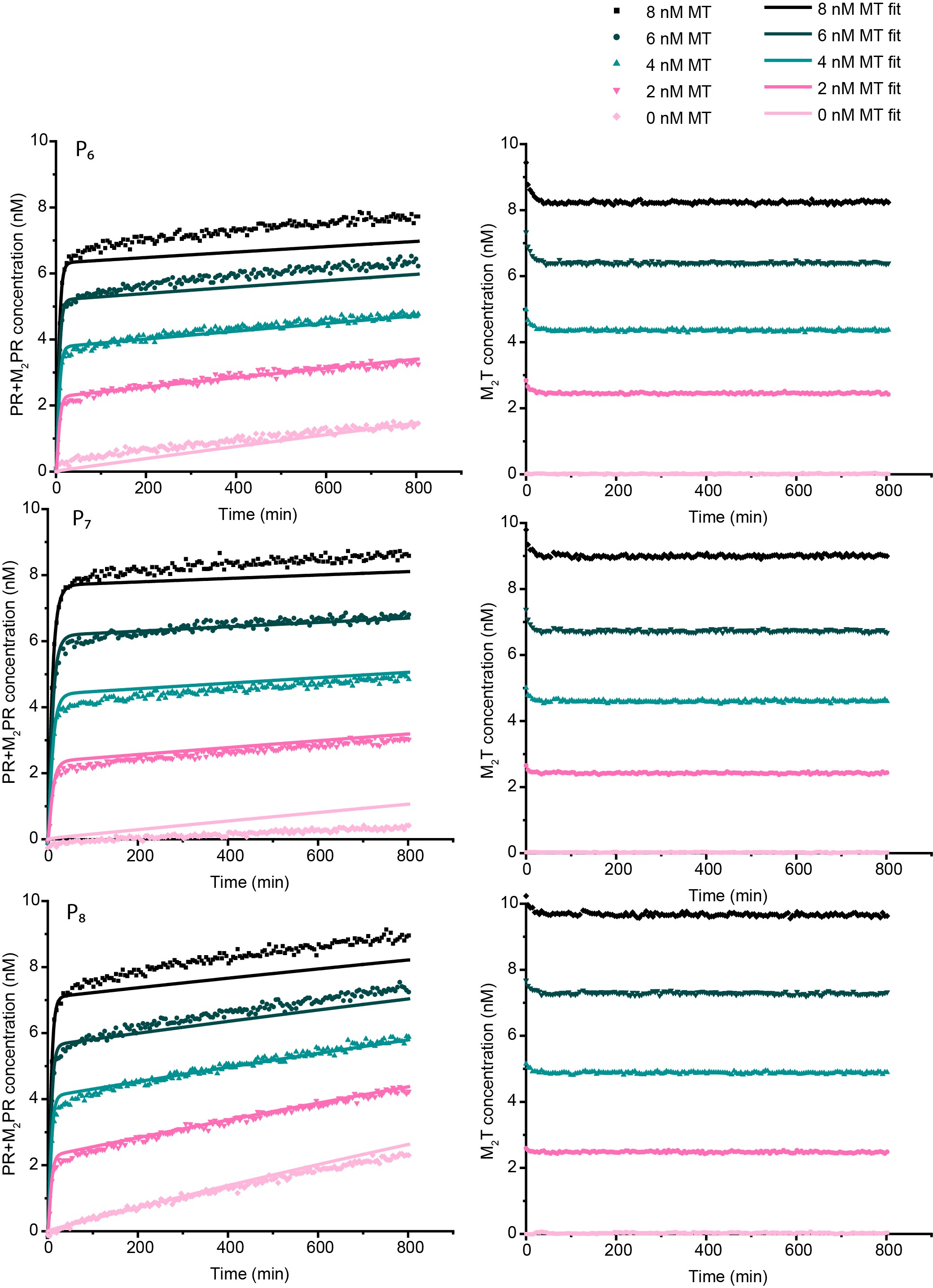

Figure S38: Template displacement by the proofreaders from M_2_T complex

50 nM of proofreader strands were added to a solution containing 20 nM *RQ* complex and 0-8 nM of *M_2_T* complex, and fluorescence signals were measured in both AlexaFluor-647 (left column) and Cy3 (right column) channels. AlexaFluor-647 reading showed the production of *M_2_PR* complexes and Cy3 measurements were used for estimating the exact concentration of added *M_2_T* complex. Three proofreaders were tested; Top: *P_6_*, middle: *P_7_*, bottom: *P_8._* Filled coloured symbols are experimentally obtained data as described in Supplementary note 12. Solid lines are the fits for these data. Note that *k_3_, k_4_* (forward and backward proofreader rates) were not obtained independently and their calculation has been described in detail in Supplementary note 12. The values of *k_3_, k_4_* are obtained alongside *k_5_, k_6_, k_7_* while using previously calculated values of *k_1_, k_2_*. All parameter values are given in Table S9-S11.

### Supplementary Figure 39: Template recovery for *M_3_T* complex

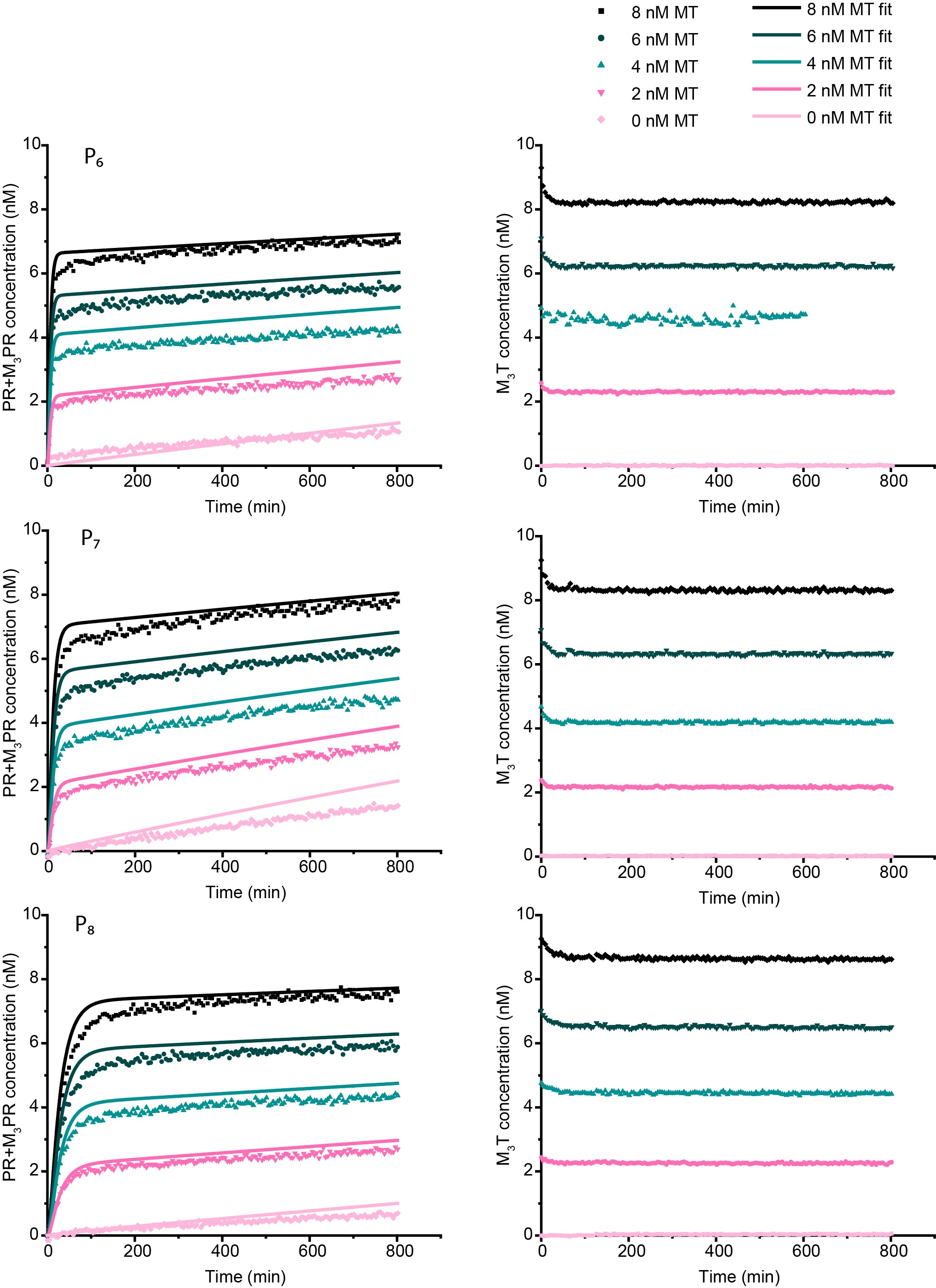

Figure S39: Template displacement by the proofreaders from M_3_T complex

50 nM of proofreader strands were added to a solution containing 20 nM *RQ* complex and 0-8 nM of *M_3_T* complex, and fluorescence signals were measured in both AlexaFluor-647 (left column) and Cy3 (right column) channels. AlexaFluor-647 reading showed the production of *M_3_PR* complexes and Cy3 measurements were used for estimating the exact concentration of added *M_3_T* complex. Three proofreaders were tested; Top: *P_6_*, middle: *P_7_*, bottom: *P_8_.* Filled coloured symbols are experimentally obtained data as described in Supplementary note 12. Solid lines are the fits for these data. Note that *k_3_, k_4_* (forward and backward proofreader rates) were not obtained independently and their calculation has been described in detail in Supplementary note 12. The values of *k_3_, k_4_* are obtained alongside *k_5_, k_6_, k_7_* while using previously calculated values of *k_1_, k_2_*. All parameter values are given in Table S9-S11.

### Supplementary Figure 40: Complete discard pathway for *M_1_L*

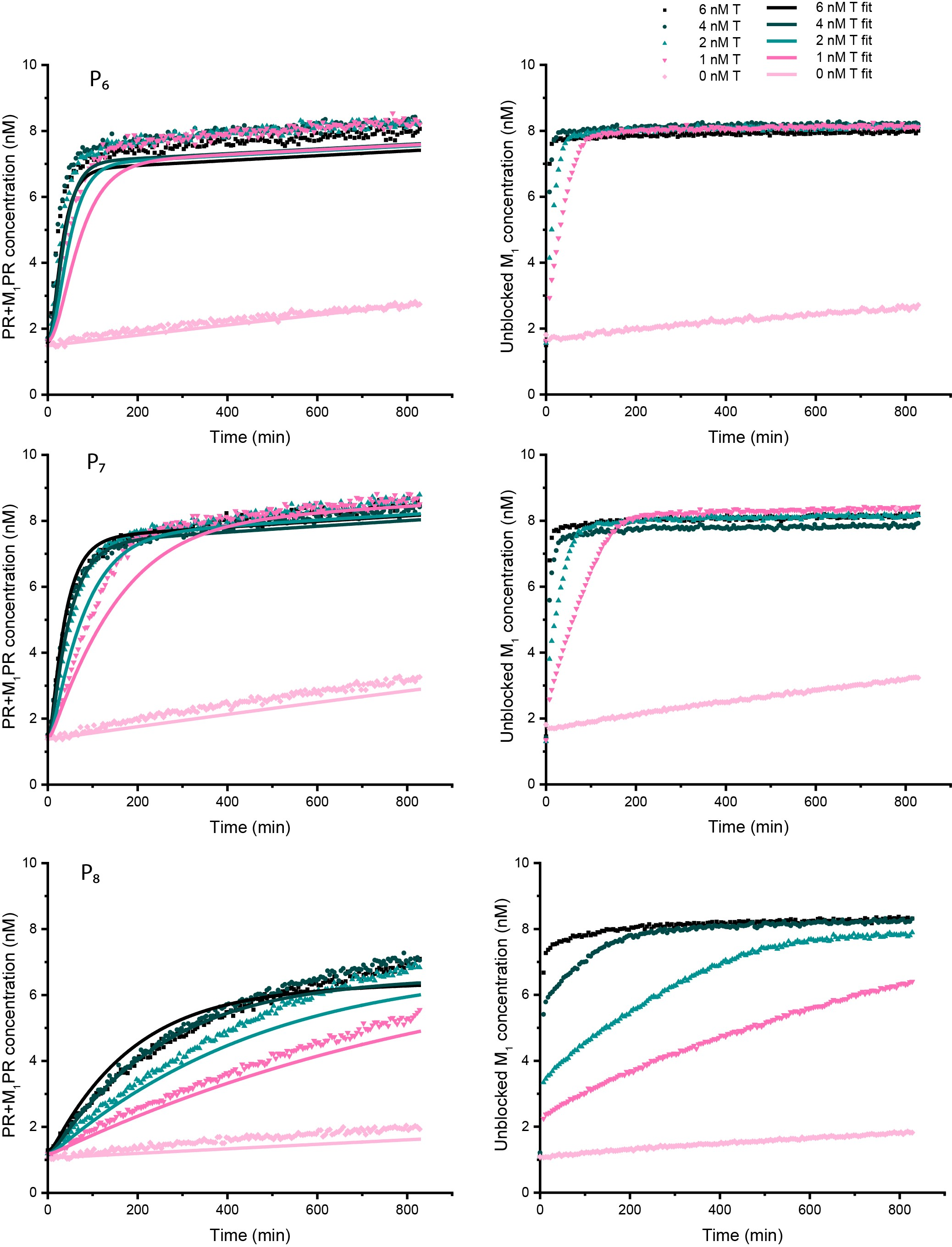

Figure S40: Conversion of blocked monomer M_1_L into waste M_1_P via template T

0-6 nM of template strands were added to a solution containing 20 nM *RQ* complex, 50 nM proofreader and 8 nM of *M_1_L* complex, and fluorescence signals were measured in both AlexaFluor-647 (left column) and Cy3 (right column) channels. AlexaFluor-647 reading showed the production of *M_1_PR* complexes and Cy3 measurements were used for monitoring the blocker strand removal from *M_1_L* complex. Three proofreaders were tested; Top: *P_6_*, middle: *P_7_*, bottom: *P_8_*. Filled coloured symbols are experimentally obtained data as described in Supplementary note 13. Solid lines are the fits for these data. Note that *k_3_, k_4_* (forward and backward proofreader rates) were not obtained independently and their calculation has been described in detail in Supplementary note 12. The values of *k_3_, k_4_* are obtained alongside *k_5_, k_6_, k_7_* while using previously calculated values of *k_1_, k_2_*. All parameter values are given in Table S9-S11.

### Supplementary Figure 41: Discard pathway for *M_2_L* complex

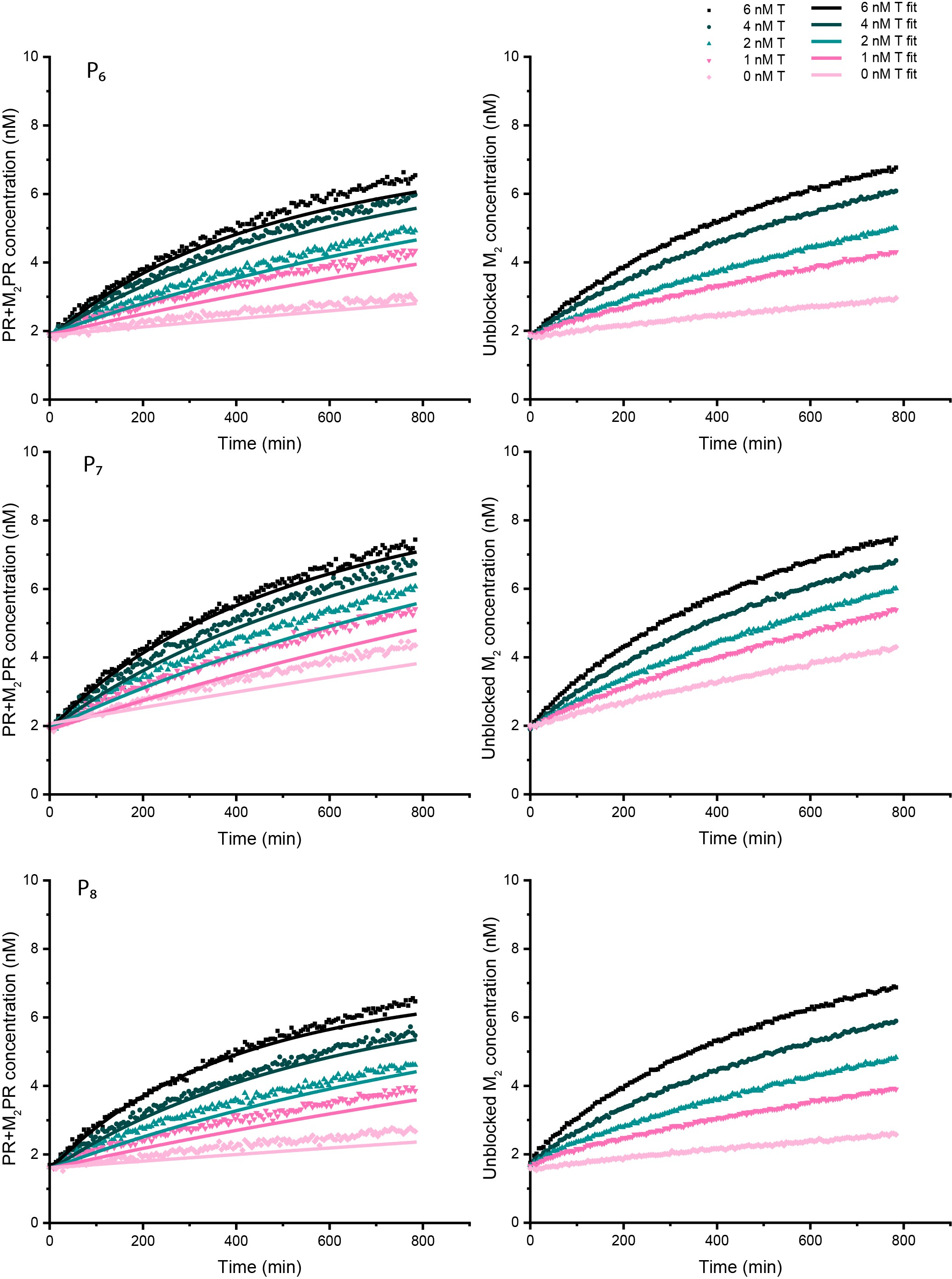

Figure S41: Conversion of blocked monomer M_2_L into waste M_2_P

0-6 nM of template strands were added to a solution containing 20 nM *RQ* complex, 50 nM proofreader and 8 nM of *M_2_L* complex, and fluorescence signals were measured in both AlexaFluor-647 (left column) and Cy3 (right column) channels. AlexaFluor-647 reading showed the production of *M_2_PR* complexes and Cy3 measurements were used for monitoring the blocker strand removal from *M_2_L* complex. Three proofreaders were tested; Top: *P_6_*, middle: *P_7_*, bottom: *P_8_*. Filled coloured symbols are experimentally obtained data as described in Supplementary note 13. Solid lines are the fits for these data. Note that *k_3_, k_4_* (forward and backward proofreader rates) were not obtained independently and their calculation has been described in detail in Supplementary note 12. The values of *k_3_, k_4_* are obtained alongside *k_5_, k_6_, k_7_* while using previously calculated values of *k_1_, k_2_*. All parameter values are given in Table S9-S11.

### Supplementary Figure 42: Discard pathway for *M_3_L* complex

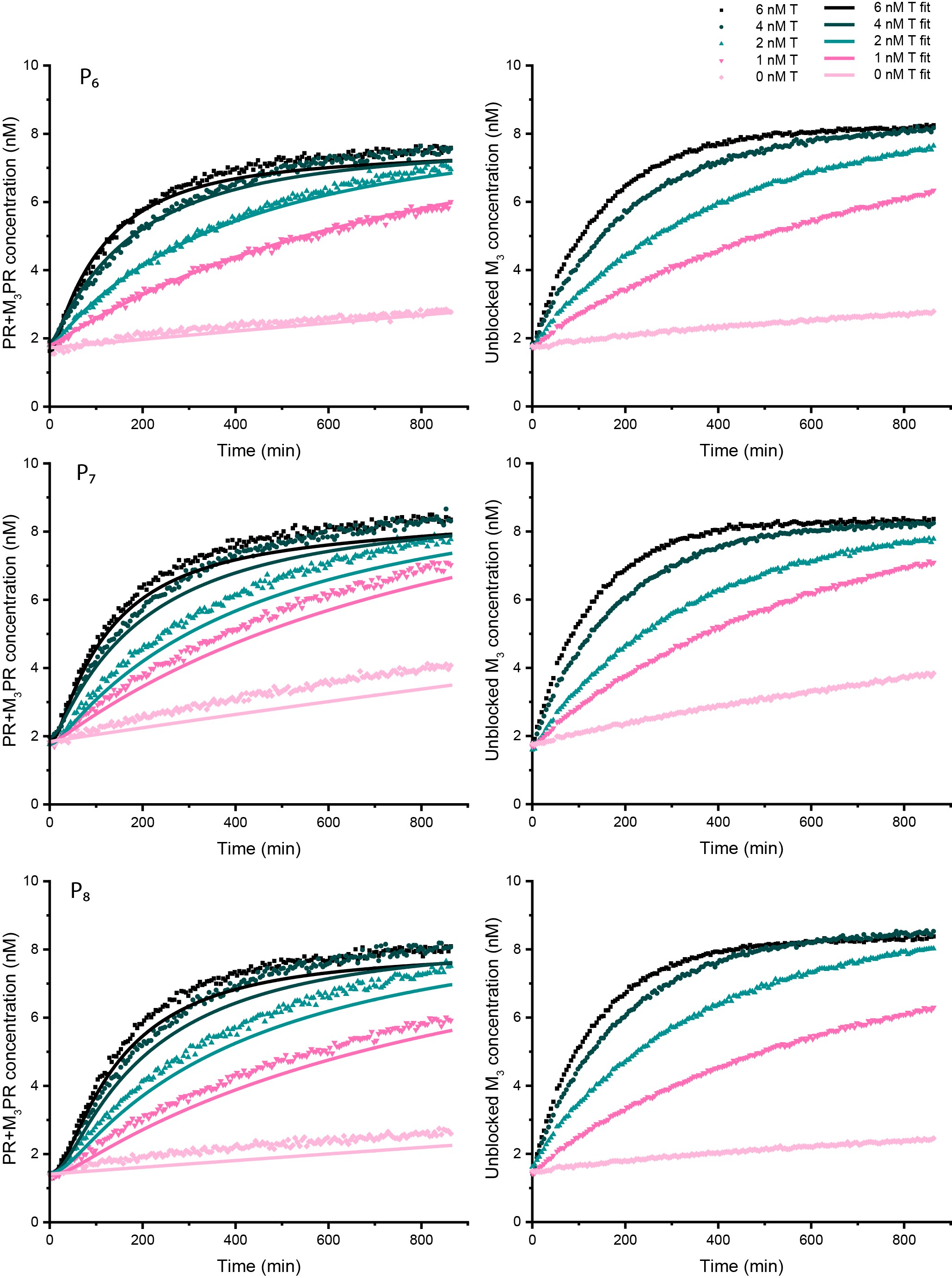

Figure S42: Conversion of blocked monomer M_3_L into waste M_3_P

0-6 nM of template strands were added to a solution containing 20 nM *RQ* complex, 50 nM proofreader and 8 nM of *M_3_L* complex, and fluorescence signals were measured in both AlexaFluor-647 (left column) and Cy3 (right column) channels. AlexaFluor-647 reading showed the production of *M_3_PR* complexes and Cy3 measurements were used for monitoring the blocker strand removal from *M_3_L* complex. Three proofreaders were tested; Top: *P_6_*, middle: *P_7_*, bottom: *P_8_*. Filled coloured symbols are experimentally obtained data as described in Supplementary note 13. Solid lines are the fits for these data obtained. Note that *k_3_, k_4_* (forward and backward proofreader rates) were not obtained independently and their calculation has been described in detail in Supplementary note 12. The values of *k_3_, k_4_* are obtained alongside *k_5_, k_6_, k_7_* while using previously calculated values of *k_1_, k_2_*. All parameter values are given in Table S9-S11.

Table S11: Rate constants for P invading MT complex

| **Monomer** | **Proofreader** | **k_3_ M^-1^ sec^-1^** | **k_4_ M^-1^ sec^-1^** | **k_7_ M^-1^ sec^-1^** |
| --- | --- | --- | --- | --- |
| *M_1_* | *P_6_* | 9.84E+04 | 2.13E+04 | 1.99E+01 |
|  | *P_7_* | 1.89E+04 | 7.30E+03 | 2.36E+01 |
|  | *P_8_* | 1.95E+03 | 1.64E+04 | 7.94E+00 |
| *M_2_* | *P_6_* | 7.20E+04 | 1.67E+01 | 3.57E+01 |
|  | *P_7_* | 7.11E+04 | 3.41E+04 | 5.91E+01 |
|  | *P_8_* | 3.70E+04 | 8.53E+03 | 1.89E+01 |
| *M_3_* | *P_6_* | 7.40E+04 | 1.67E+01 | 2.26E+01 |
|  | *P_7_* | 8.19E+04 | 1.67E+01 | 4.23E+01 |
|  | *P_8_* | 1.16E+05 | 1.67E+01 | 1.69E+01 |

The parameter values were obtained using the protocol defined in Supplementary note 11 to calculate reaction rate constants *k_3_, k_4_, k_5_, k_6_, k_7_* while using previously identified parameters *k_1_, k_2_*. All parameter values are given in Table S9-S11.

### Supplementary Figure 43: *MT* concentration estimates and model predictions

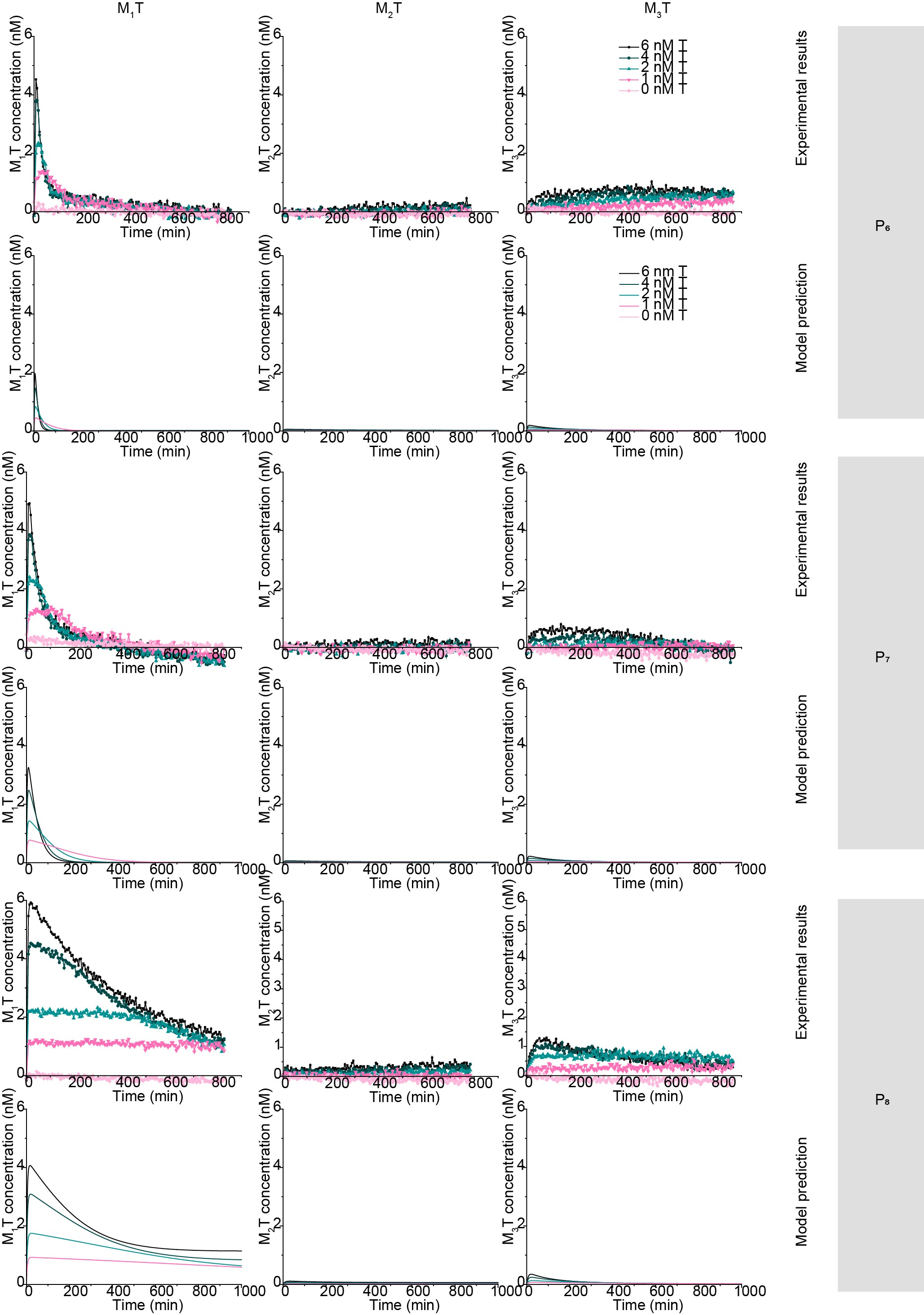

Figure S43: Experimental estimation and model prediction of MT concentrations in the discard pathway experiments

The experimental *MT* concentrations were obtained by subtracting the MPR concentrations from the unblocked monomer concentration for the discard pathway experiments (Supplementary Figure S40-42). The parameters *k_1_-k_7_* given in Tables S9-S11 were used to solve the system of ODEs given by Eq. (S31) -(S34) to estimate the concentration of *MT* (Tables S9-S11) complexes in the solution for different monomers (top two row: *M_1_*, middle two rows: *M_2_*, bottom two rows: *M_3_*) and different proofreaders (left: *P_6_*, middle: *P_7_*, right: *P_8_*). *M_1_T* concentrations are relatively high in the initial part of the experiments, providing a window for dimerization. The concentrations of *M_2_T* and *M_3_T* are predicted to remain low throughout the experiments for all three proofreaders.

### Supplementary Figure 44: Dimer formation for *M’_1_N* in presence and absence of *P’_6_*

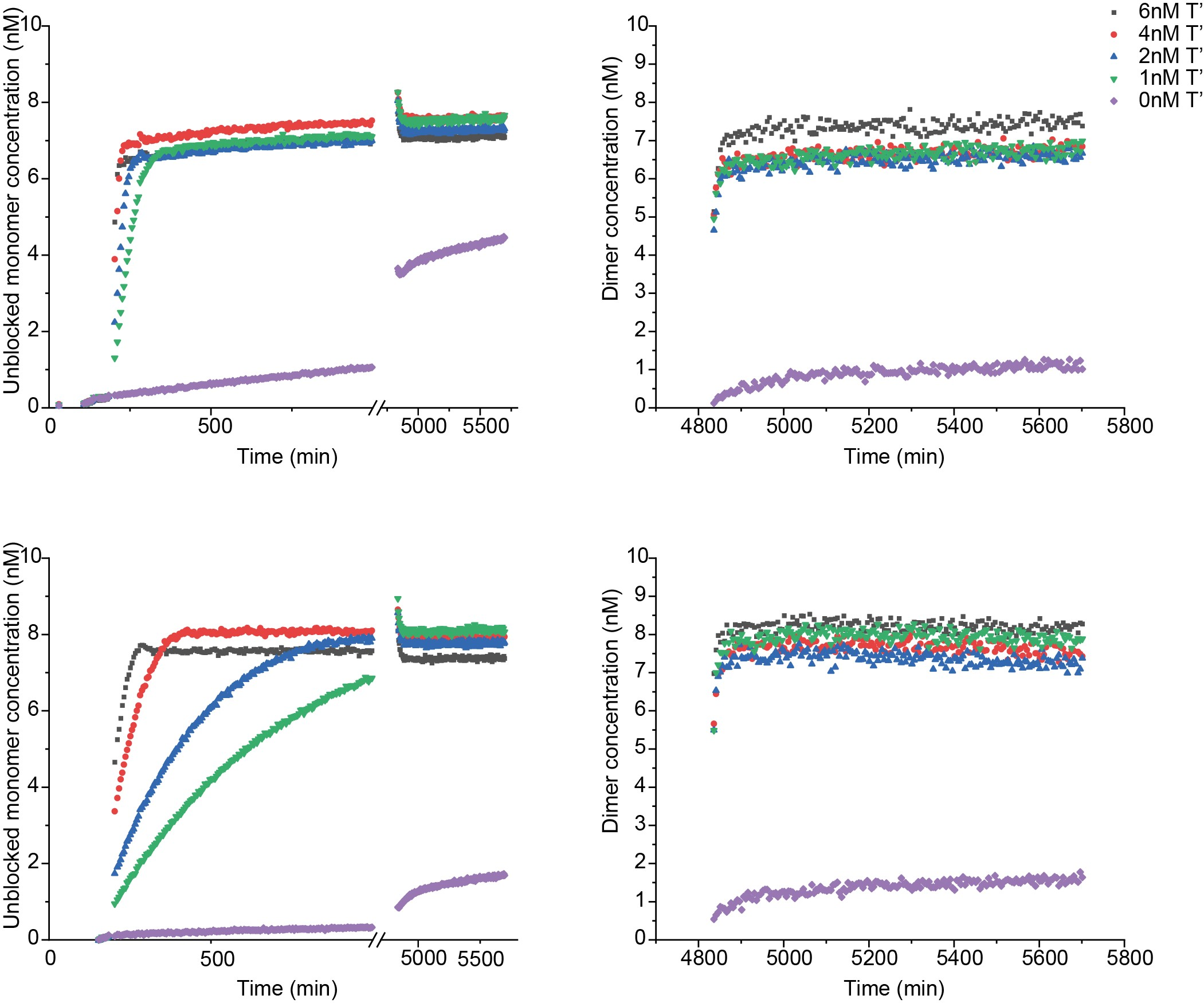

Figure S44: M’_1_N dimer formation in presence (top row) and absence (bottom row) of P’_6_.

Reactions are triggered by adding 0-6 nM of *T’* into a solution containing 8 nM of *M’_1_L’*, 50 or 0 nM *P’_6_*, and 10 nM *N*. The progress of the reactions was followed by monitoring the Cy3 channel (left column). When all the experiments reached a plateau at a similar level of fluorescence signal intensities, 20 nM of external reporter *RQ* was added to the reaction to measure the amount of *M’_1_N* formed by monitoring the AlexaFluor-647 fluorescence signal (right column). The reporter was not added at t=0 as it induces leak in dimerization reaction. The leak is more evident on the 0 nM *T’* plot on the bottom left Figure - note the jump in Cy3 signal after the break on x-axis when reporter is added. The data are processed as described in Supplementary note 15.

### Supplementary Figure 45: Dimer formation for *M’_2_N* in presence and absence of *P’_6_*

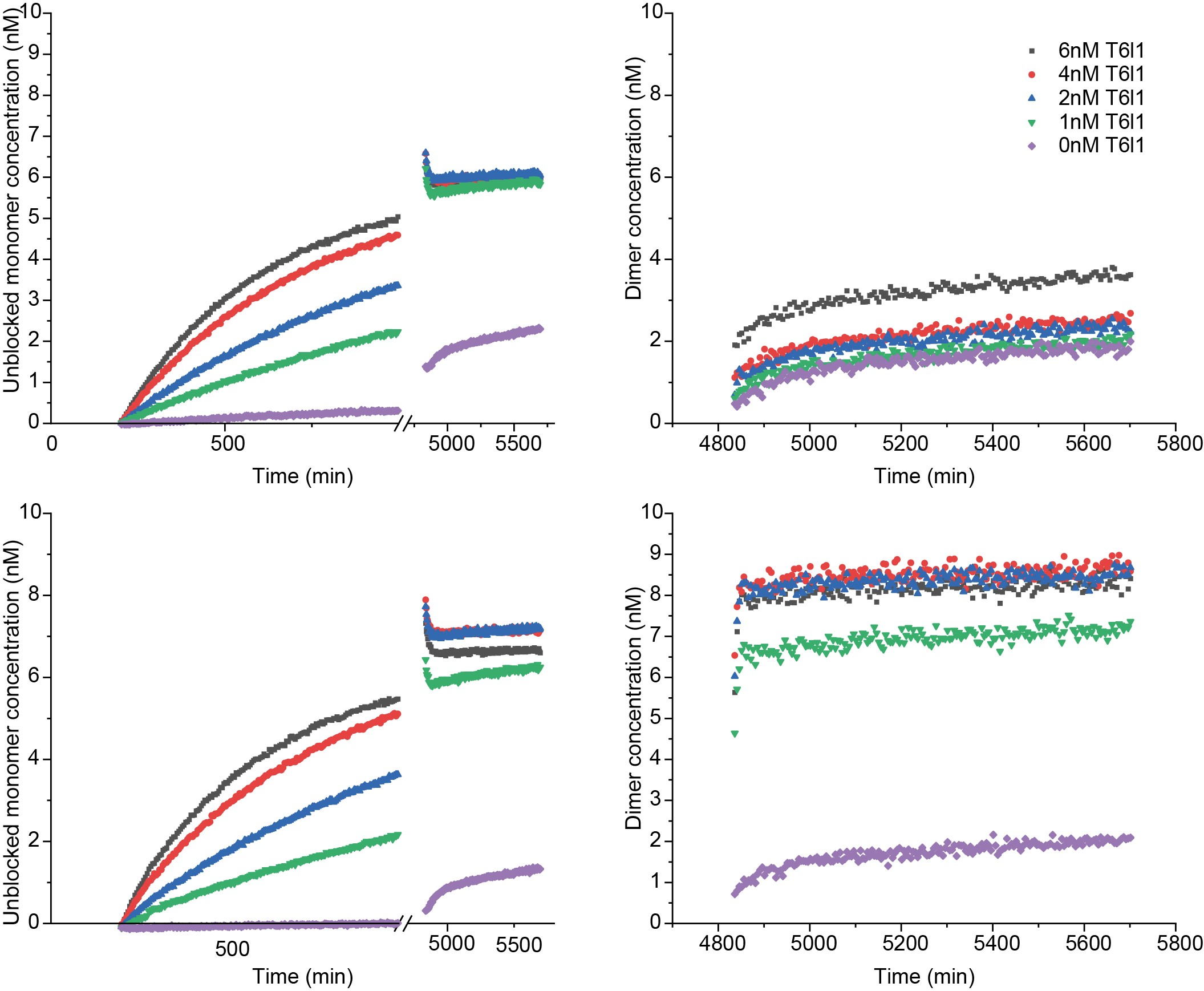

Figure S45: M’_2_N dimer formation in presence (top row) and absence (bottom row) of P’_6_.

Reactions are triggered by adding 0-6 nM of *T’* into a solution containing 8 nM of *M’_2_L’*, 50 or 0 nM *P’_6_*, and 10 nM *N*. The progress of the reactions was followed by monitoring the Cy3 channel (left column). When all the experiments reached a plateau at almost similar level of fluorescence signal intensities, 20 nM of external reporter *RQ* was added to the reaction to measure the amount of *M’_2_N* formed by monitoring the AlexaFluor-647 fluorescence signal (right column). The data are processed as described in Supplementary note 15.

### Supplementary Figure 46: Dimer formation for *M’_3_N* in presence and absence of *P’_6_*

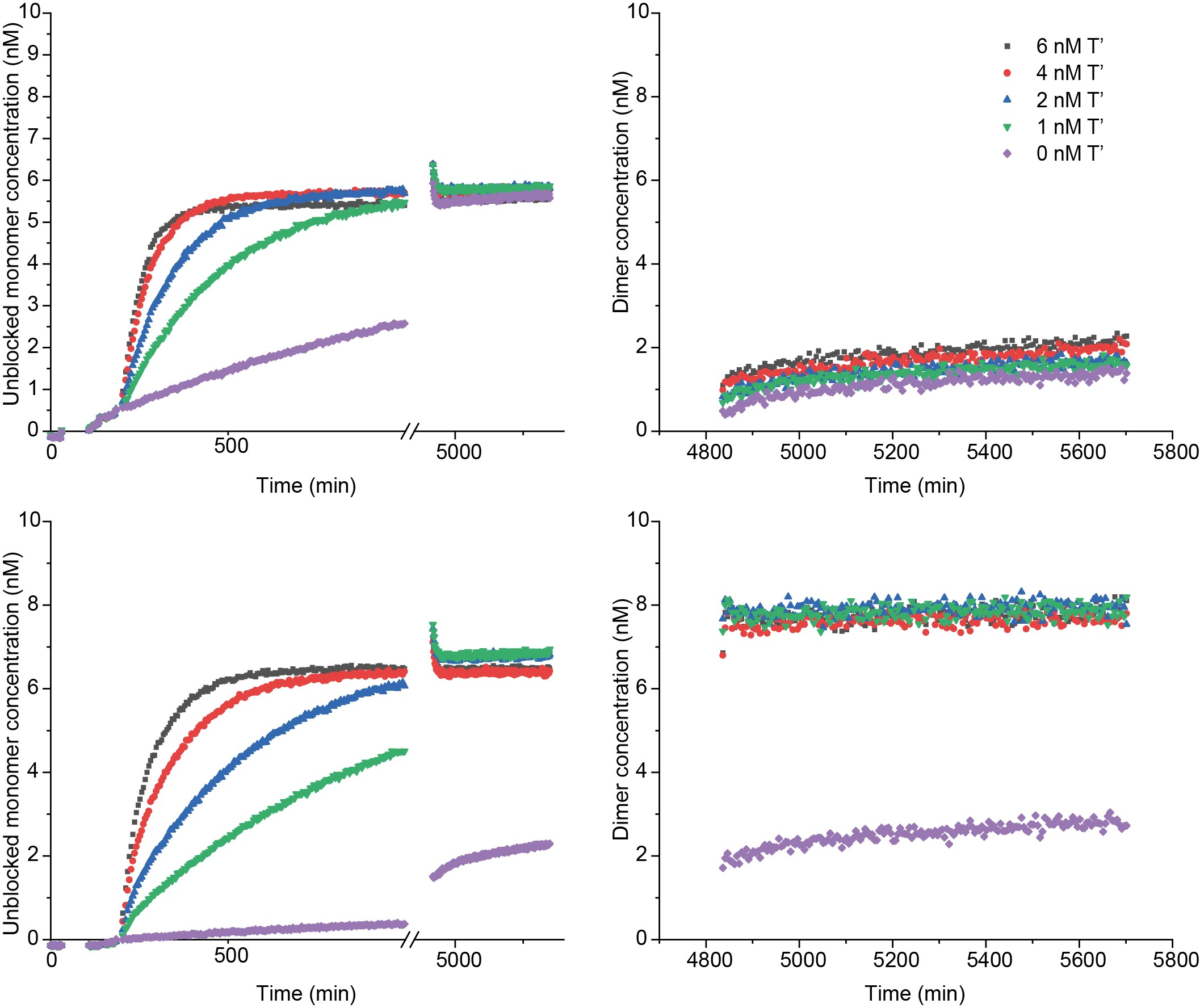

Figure S46: M’_3_N dimer formation in presence (top row) and absence (bottom row) of P’_6_.

Reactions are triggered by adding 0-6 nM of *T’* into a solution containing 8 nM of *M’_3_L*’, 50 or 0 nM *P’_6_*, and 10 nM *N*. The progress of the reactions was followed by monitoring the Cy3 channel (left column). When all the experiments reached a plateau at almost similar level of fluorescence signal intensities, 20 nM of external reporter *RQ* was added to the reaction to measure the amount of *M’_3_N* formed by monitoring the AlexaFluor-647 fluorescence signal (right column). The data are processed as described in Supplementary note 15.

### Supplementary Figure 47: Dimer formation for *M’_1_N* in presence and absence and absence of *P’_7_*

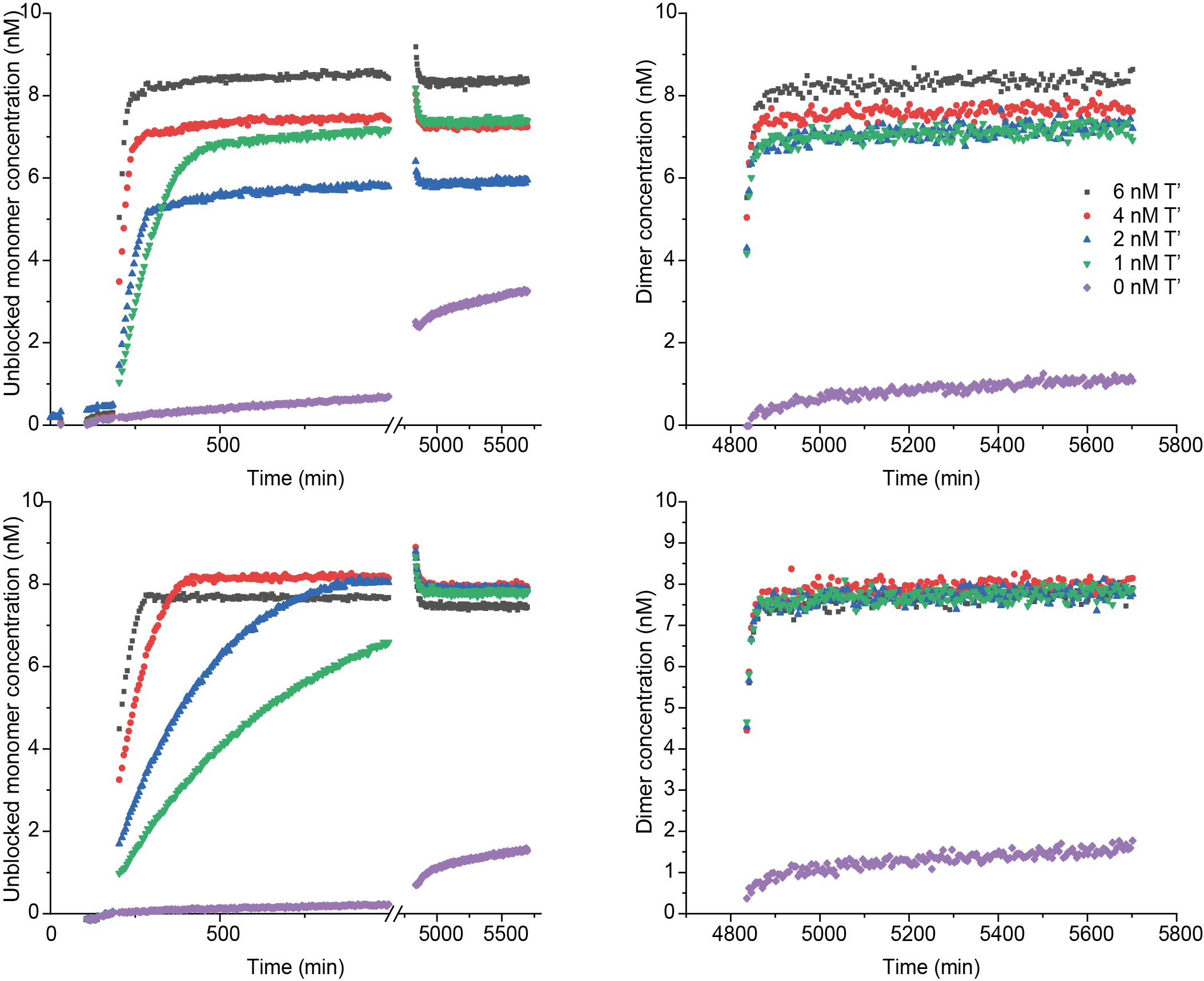

Figure S47: M’_1_N dimer formation in presence (top row) and absence (bottom row) of P’_7_.

Reactions are triggered by adding 0-6 nM of *T’* into a solution containing 8 nM of *M’_1_L’*, 50 or 0 nM *P’_7_*, and 10 nM *N*. The progress of the reactions was followed by monitoring the Cy3 channel (left column). When all the experiments reached a plateau at almost similar level of fluorescence signal intensities, 20 nM of external reporter *RQ* was added to the reaction to measure the amount of *M’_1_N* formed by monitoring the AlexaFluor-647 fluorescence signal (right column). The data are processed as described in Supplementary note 15.

### Supplementary Figure 48: Dimer formation for *M’_2_N* in presence and absence of *P’_7_*

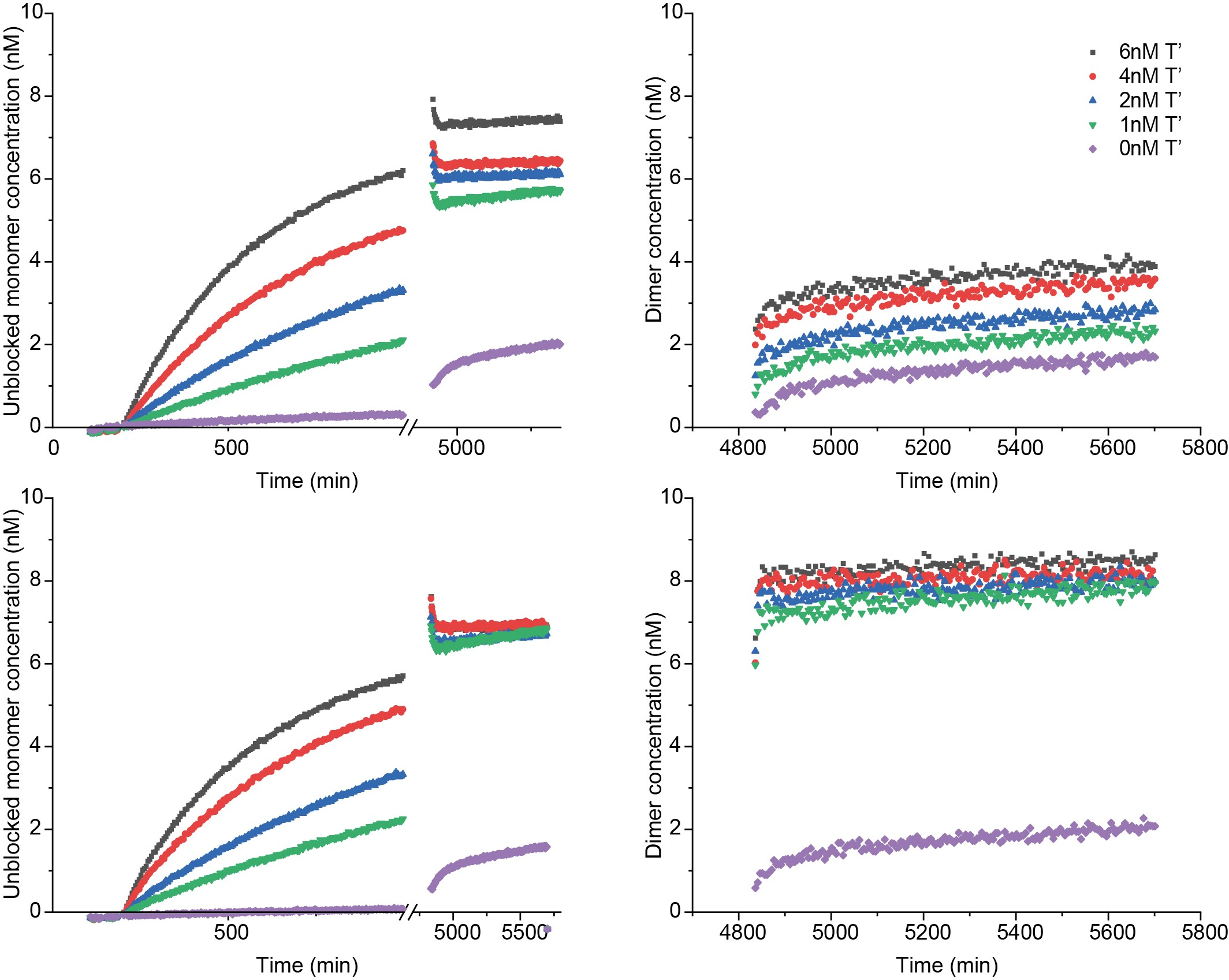

Figure S48: M’_2_N dimer formation in presence (top row) and absence (bottom row) of P’_7_

Reactions are triggered by adding 0-6 nM of *T’* into a solution containing 8 nM of *M’_2_L’*, 50 or 0 nM *P’_7_*, and 10 nM *N*. The progress of the reactions was followed by monitoring the Cy3 channel (left column). When all the experiments reached a plateau at almost similar level of fluorescence signal intensities, 20 nM of external reporter *RQ* was added to the reaction to measure the amount of *M’_2_N* formed by monitoring the AlexaFluor-647 fluorescence signal (right column). The data are processed as described in Supplementary note 15.

### Supplementary Figure 49: Dimer formation for *M’_3_N* in presence and absence of *P’_7_*

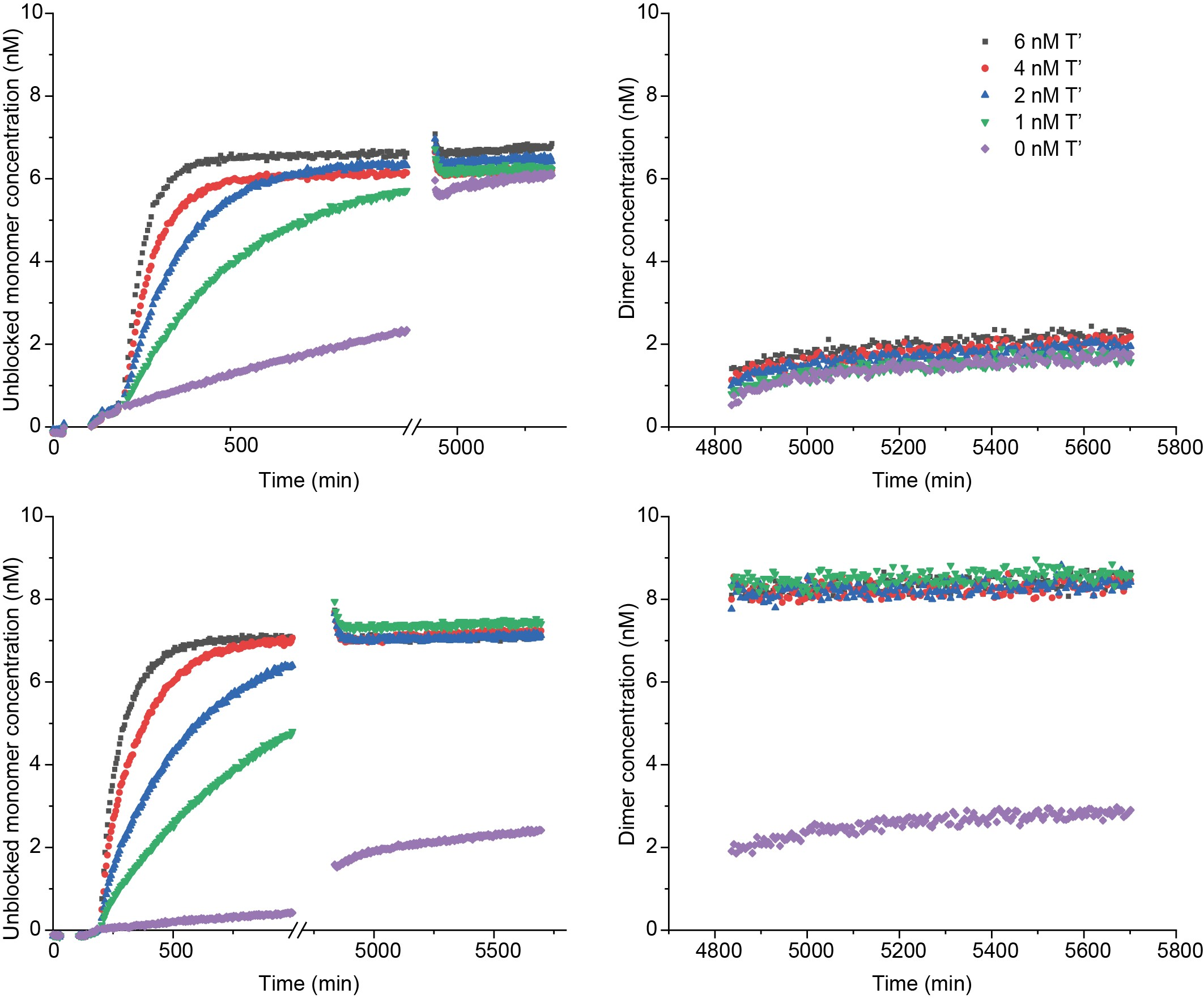

Figure S49: M’_3_N dimer formation in presence (top row) and absence (bottom row) of P’_7_.

Reactions are triggered by adding 0-6 nM of *T’* into a solution containing 8 nM of *M’_3_L’*, 50 or 0 nM *P_7_*, and 10 nM *N*. The progress of the reactions was followed by monitoring the Cy3 channel (left column). When all the experiments reached a plateau at almost similar level of fluorescence signal intensities, 20 nM of external reporter *RQ* was added to the reaction to measure the amount of *M’_3_N* formed by monitoring the AlexaFluor-647 fluorescence signal (right column). The data are processed as described in Supplementary note 15.

### Supplementary Figure 50: Dimer formation for *M’_1_N* in presence and absence of *P’_8_*

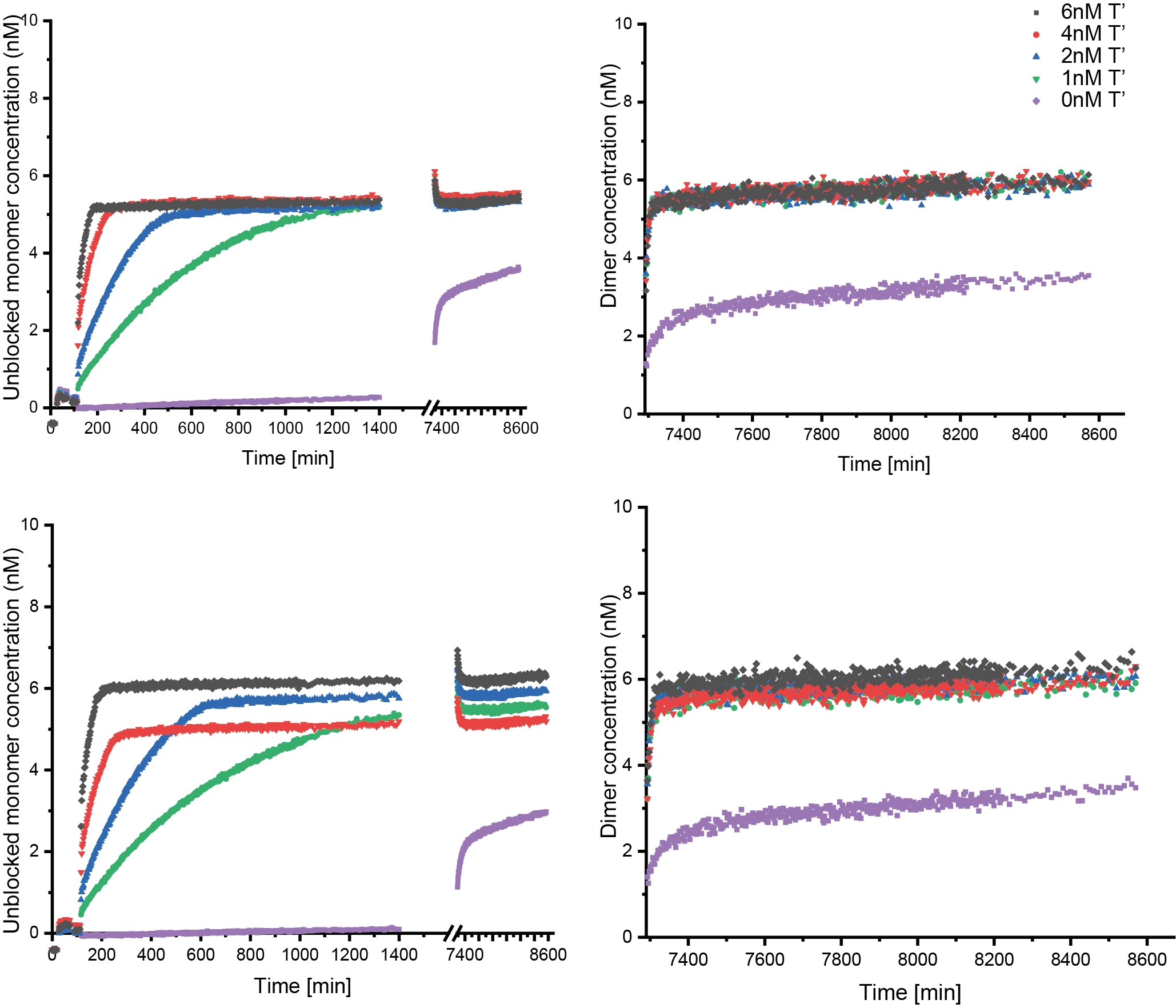

Figure S 50: M’_1_N dimer formation in presence (top row) and absence (bottom row) of P’_8_.

Reactions are triggered by adding 0-6 nM of *T’* into a solution containing 8 nM of *M’_1_L’*, 50 or 0 nM *P’_8_*, and 10 nM *N*. The progress of the reactions was followed by monitoring the Cy3 channel (left column). When all the experiments reached a plateau at almost similar level of fluorescence signal intensities, 20 nM of external reporter *RQ* was added to the reaction to measure the amount of *M’_1_N* formed by monitoring the AlexaFluor-647 fluorescence signal (right column). The data are processed as described in Supplementary note 15.

### Supplementary Figure 51: Dimer formation for *M’_2_N* in presence and absence of *P’_8_*

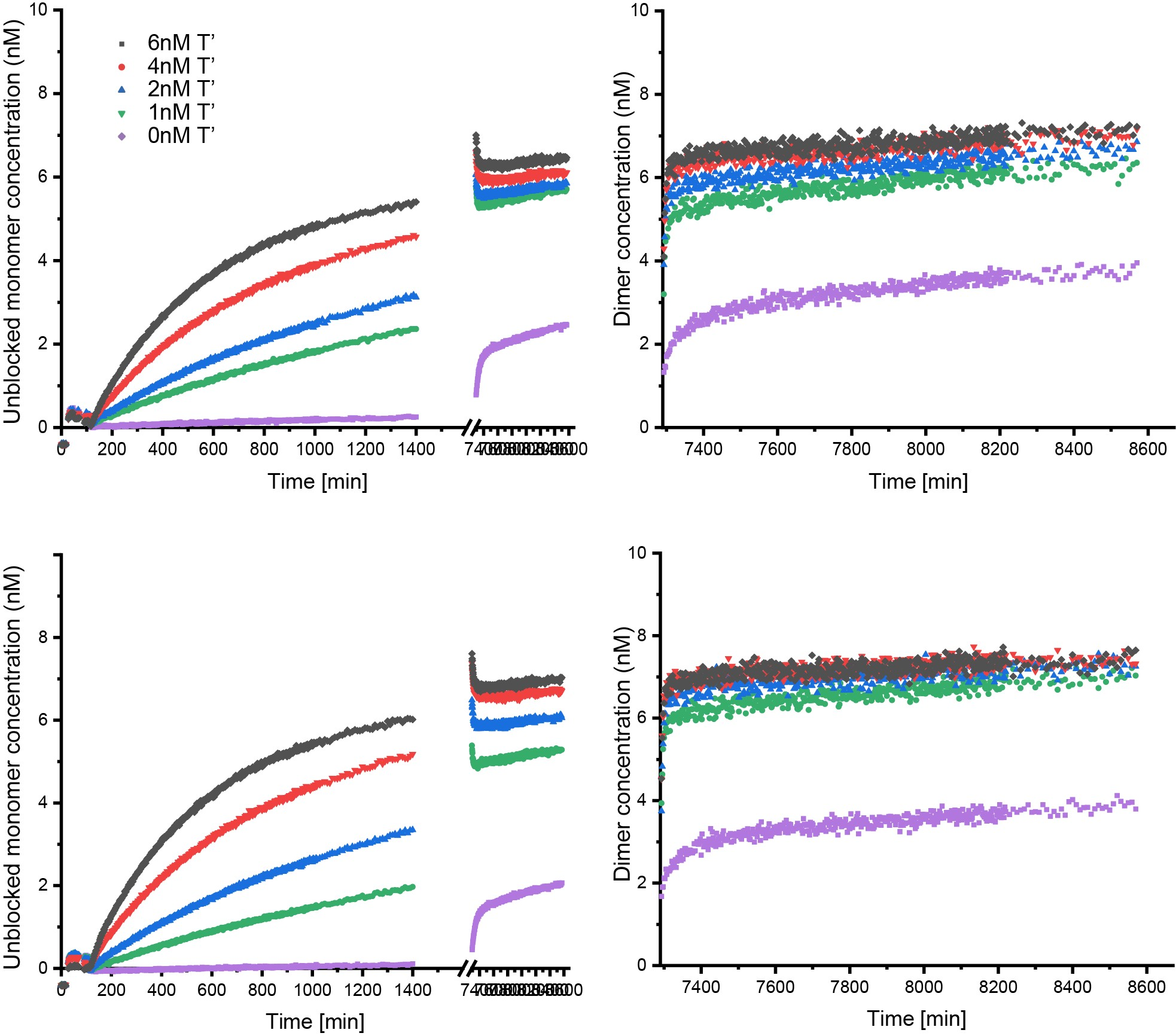

Figure S 51: M’_2_N dimer formation in presence (top row) and absence (bottom row) of P’_8_.

Reactions are triggered by adding 0-6 nM of *T’* into a solution containing 8 nM of *M’_2_L’*, 50 or 0 nM *P’_8_*, and 10 nM *N*. The progress of the reactions was followed by monitoring the Cy3 channel (left column). When all the experiments reached a plateau at almost similar level of fluorescence signal intensities, 20 nM of external reporter *RQ* was added to the reaction to measure the amount of *M’_2_N* formed by monitoring the AlexaFluor-647 fluorescence signal (right column). The data are processed as described in Supplementary note 15.

### Supplementary Figure 52: Dimer formation for *M’_3_N* in presence and absence of *P’_8_*

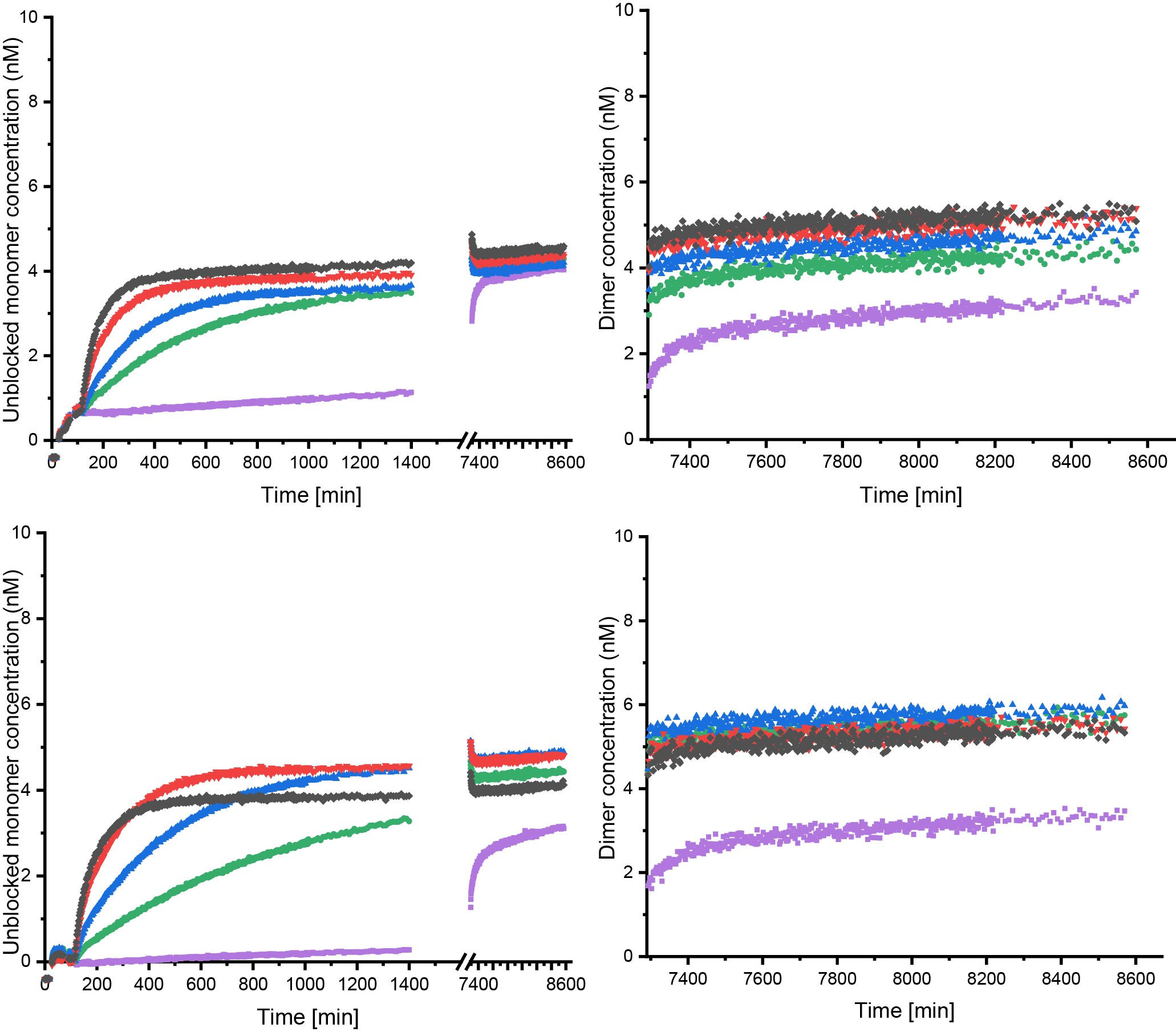

Figure S 52: M’_3_N dimer formation in presence (top row) and absence (bottom row) of P’_8_.

Reactions are triggered by adding 0-6 nM of *T’* into a solution containing 8 nM of *M’_3_L’*, 50 or 0 nM *P’_8_*, and 10 nM *N*. The progress of the reactions was followed by monitoring the Cy3 channel (left column). When all the experiments reached a plateau at almost similar level of fluorescence signal intensities, 20 nM of external reporter *RQ* was added to the reaction to measure the amount of *M’_3_N* formed by monitoring the AlexaFluor-647 fluorescence signal (right column). The data are processed as described in Supplementary note 15.

### Supplementary Figure 53: Dimer formation with and without proofreaders as predicted by ODE model

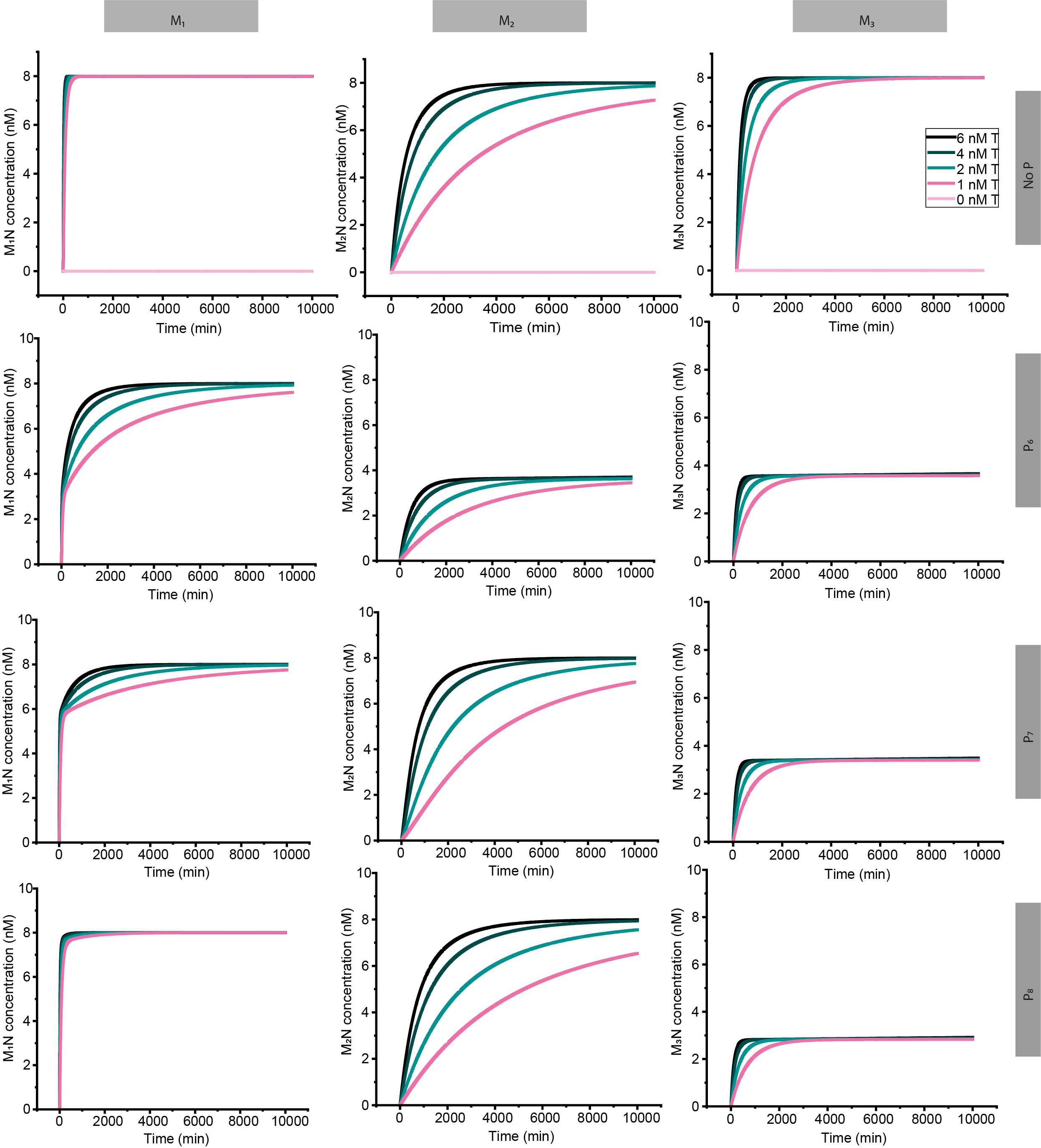

Figure S53: Prediction of dimerization rates with and without different proofreaders

The rates of dimerization with and without different proofreaders generated by the ODE model Eqs. (S51) -(S53) described in Supplementary note 14. The initial conditions at t=0 are [*M’L’*]=[*M’L’_total_*]=8nM, [*T’*]=[*T’_total_*] is varied from {6nM, 4nM, 2nM, 1nM, 0nM}, [*N*] = [*N_total_*] = 10nM, [*P’*] = [*P’_total_*] = 50nM. Parameters *k_1_, k_2_, k_3_, k_4_* have been derived from experimental data and values are shown in Tables S9 and S11. Parameters {*k_8_*, *k_9_*} were approximated by doing some preliminary experiments as {3.50E+05, 0} respectively. For *M’_1_* with proofreaders *P’_6_* and *P’_7_*, and *M’_2_* with proofreaders *P’_7_* and *P’_8_*, we see evidence of previously proofread molecules being rebound to the template and dimerised over long timescales, as discussed in Supplementary note 9. For all *M’_2_* and *M’_3_* traces, moderate concentrations of product are reached quickly because the initial proofreading reaction is only partially successful in out-competing dimerization for the parameters used here.

### Supplementary Figure 54: Dimer formation for *M’_3_N* in presence and absence of excess blocker strand

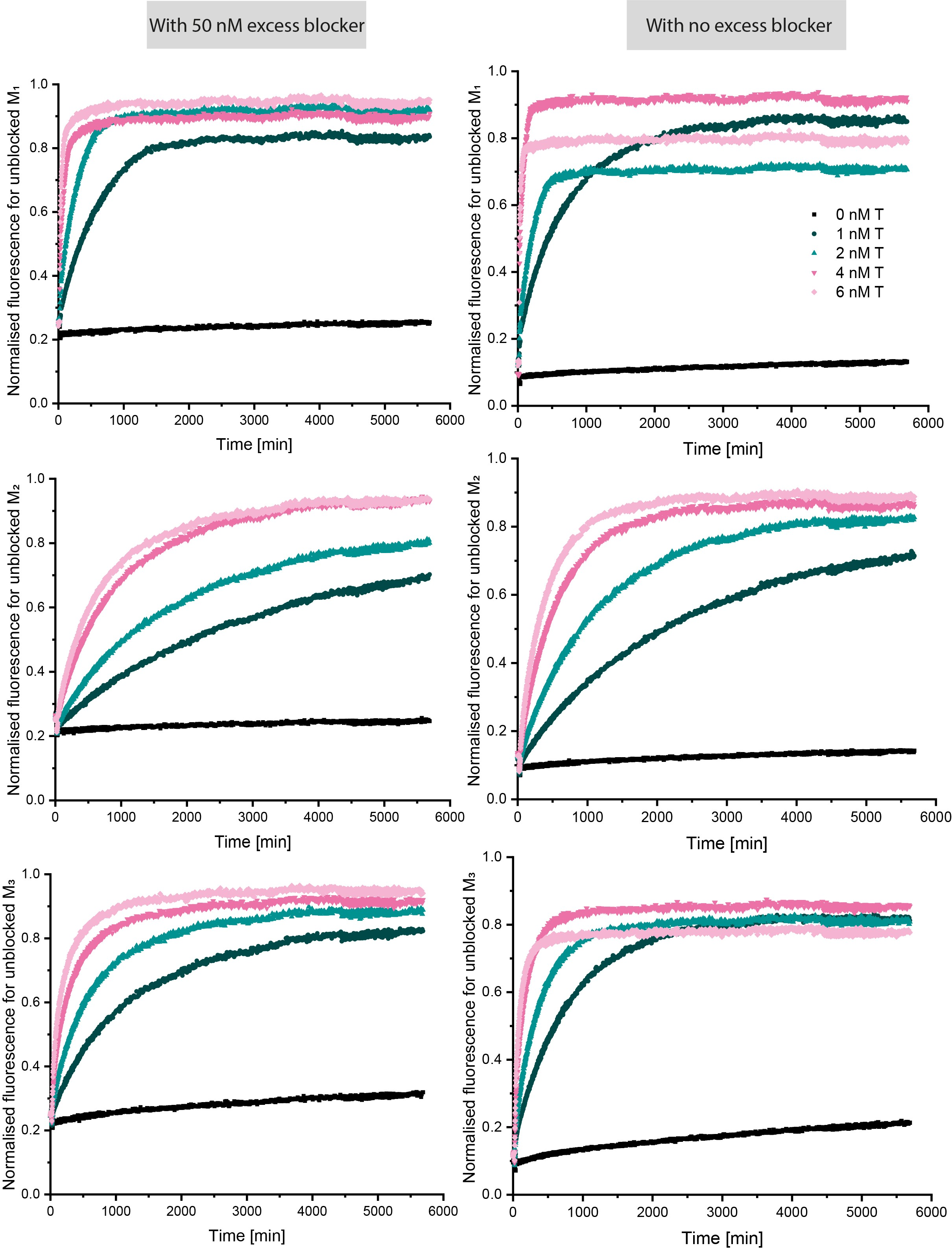

Figure S54: M’_3_N dimer formation in presence (left) and absence (right) of excess blocker

Normalised cy3 fluorescence data shows the extent of dimer formation for each monomer in presence and absence of excess blocker showing it’s fundamental difference from a proofreader. Reactions are triggered by adding 0-6 nM of *T’* into a solution containing 8 nM of *M’_3_L’*, 50 or 0 nM excess *L’*, and 10 nM *N*. The progress of the reactions was followed by monitoring the Cy3 channel. Fluorescence signals are normalised to the highest signal intensity for each set of experiments (a set of experiment here is a monomer with or without excess blocker triggered by injecting different concentration of the template *T’*) for each monomer. Since there is no alternative proofreading pathway, the increase in fluorescence implies template binding and subsequent incorporation into products *M’_i_N*. The limited difference in both the kinetics of product formation and its final yield show that the blockers do not achieve the enhanced discrimination of the proofreader.

### Supplementary Figure 55: Formation of a *M’_1_N* dimer from a mixture of *M’_1_*, *M’_2_* and *M’_3_*, in the presence and absence of *P’_6_*

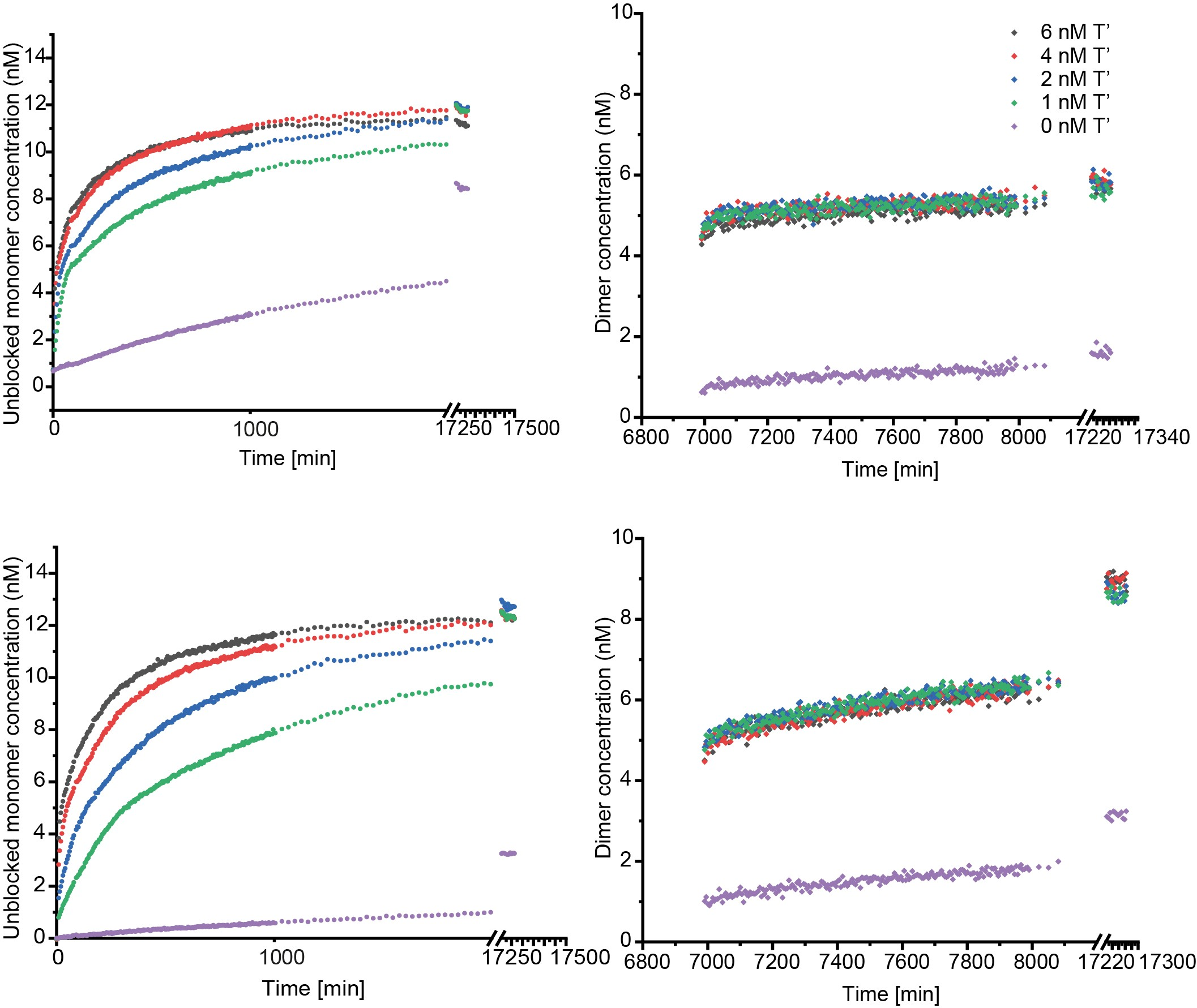

Figure S55: M’_1_N dimer formation from a mixture of M’ monomers in presence (top row) and absence (bottom row) of P’_6_

Reactions are triggered by adding 0-6 nM of *T’* into a solution containing 5 nM of each *M’_1_L’*, *M’_2_L’* and *M’_3_L’*, 50 or 0 nM *P’_6_*, and 15 nM *N*. The progress of the reactions was followed by monitoring the Cy3 channel (left column). When all the experiments reached a plateau at almost similar level of fluorescence signal intensities, 20 nM of external reporter *RQ* was added to the reaction to measure the concentration of *M’_1_N* formed by monitoring the AlexaFluor-647 fluorescence signal (right column). The reporter was not added at t=0 as it induces leak in dimerization reaction. The data are processed as outlined in Supplementary note 15.

### Supplementary Figure 56: Formation of a *M’_2_N* dimer from a mixture of *M’_1_*, *M’_2_* and *M’_3_*, in the presence and absence of *P’_6_*

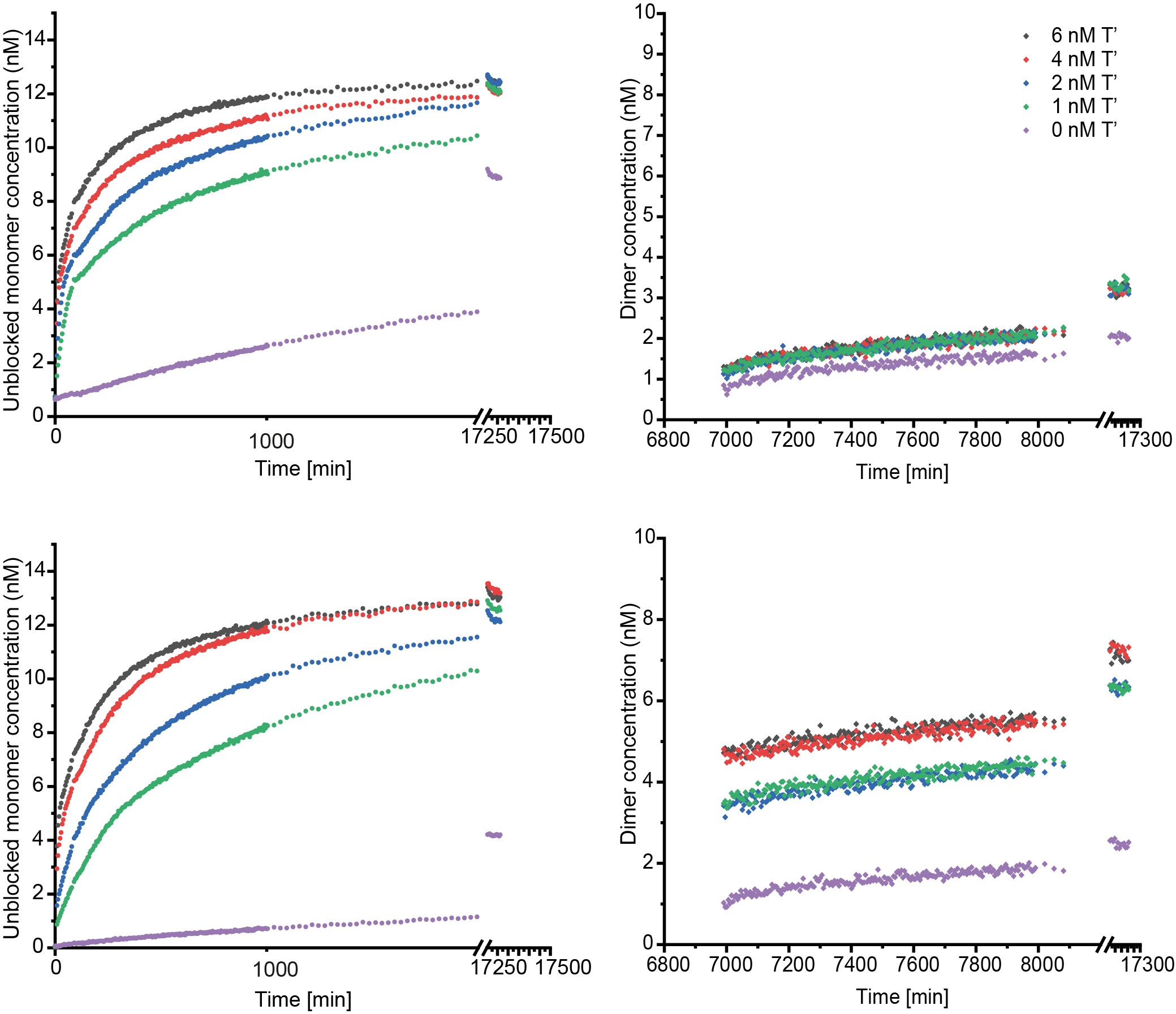

Figure S56: M’_2_N dimer formation from a mixture of M’ monomers in presence (top row) and absence (bottom row) of P’_6_

Reactions are triggered by adding 0-6 nM of *T’* into a solution containing 5 nM of each *M’_1_L’*, *M’_2_L’* and *M’_3_L’*, 50 or 0 nM *P’_6_*, and 10 nM *N*. The progress of the reactions was followed by monitoring the Cy3 channel (left column). When all the experiments reached a plateau at almost similar level of fluorescence signal intensities, 20 nM of external reporter *RQ* was added to the reaction to measure the amount of *M’_2_N* formed by monitoring the AlexaFluor-647 fluorescence signal (right column). The reporter was not added at t=0 as it induces leak in dimerization reaction. The data are processed as outlined in Supplementary note 15.

### Supplementary Figure 57: Formation of a *M’_3_N* dimer from a mixture of *M’_1_*, *M’_2_* and *M’_3_*, in the presence and absence of *P’_6_*

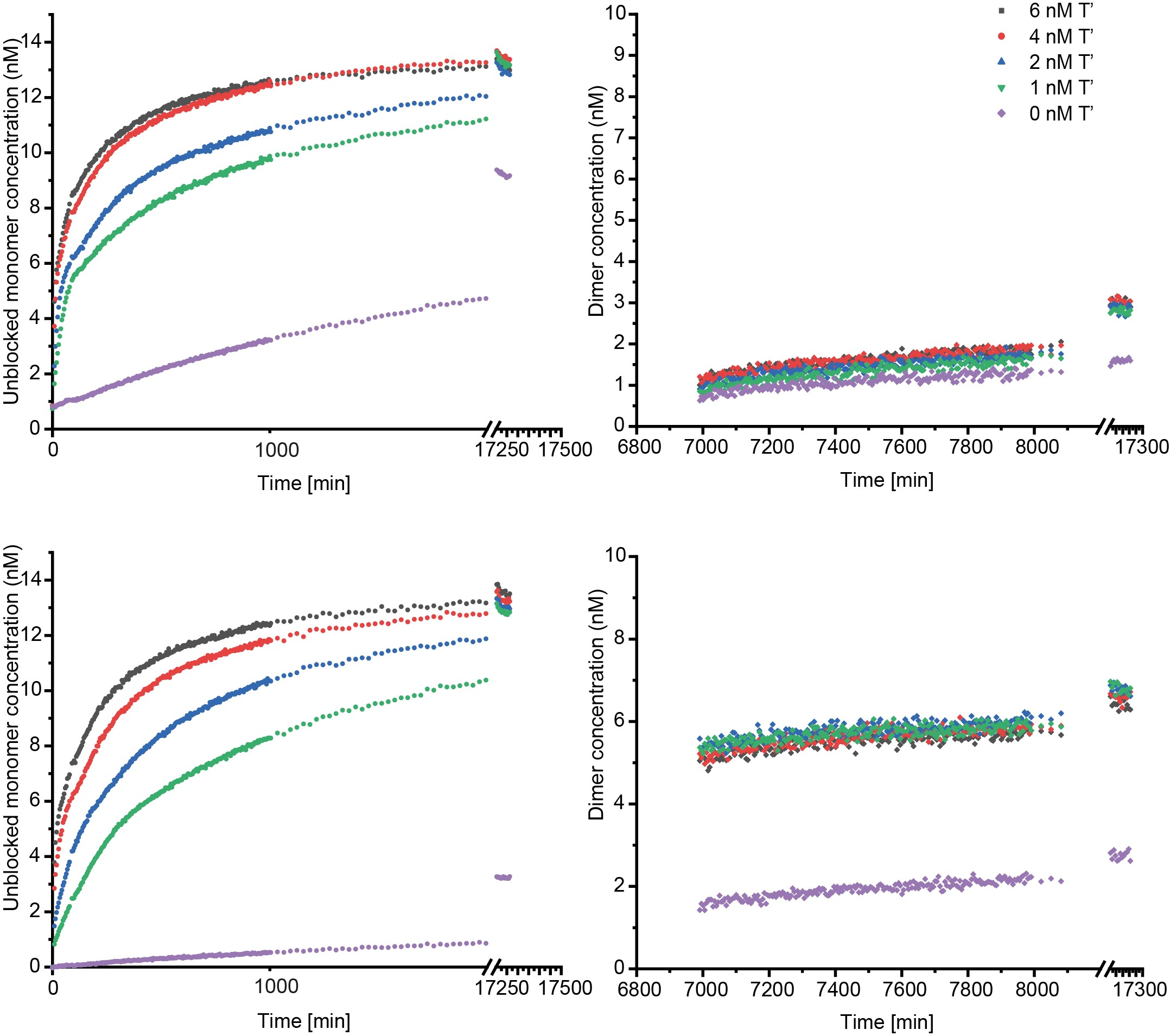

Figure S57: M’_3_N dimer formation in presence (top row) and absence of P’_6_ (bottom row) from a mixture of monomers

Reactions are triggered by adding 0-6 nM of *T’* into a solution containing 5 nM of each *M’_1_L’*, *M’_2_L’* and *M’_3_L’*, 50 or 0 nM *P’_6_*, and 10 nM *N*. The progress of the reactions was followed by monitoring the Cy3 channel (left column). When all the experiments reached a plateau at almost similar level of fluorescence signal intensities, 20 nM of external reporter *RQ* was added to the reaction to measure the amount of *M’_3_N* formed by monitoring the AlexaFluor-647 fluorescence signal (right column). The reporter was not added at t=0 as it induces leak in dimerization reaction. The data are processed as outlined in Supplementary note 15.

### Supplementary Figure 58: Formation of a *M’_1_N* dimer from a mixture of *M’_1_*, *M’_2_* and *M’_3_*, in the presence and absence of *P’_7_*

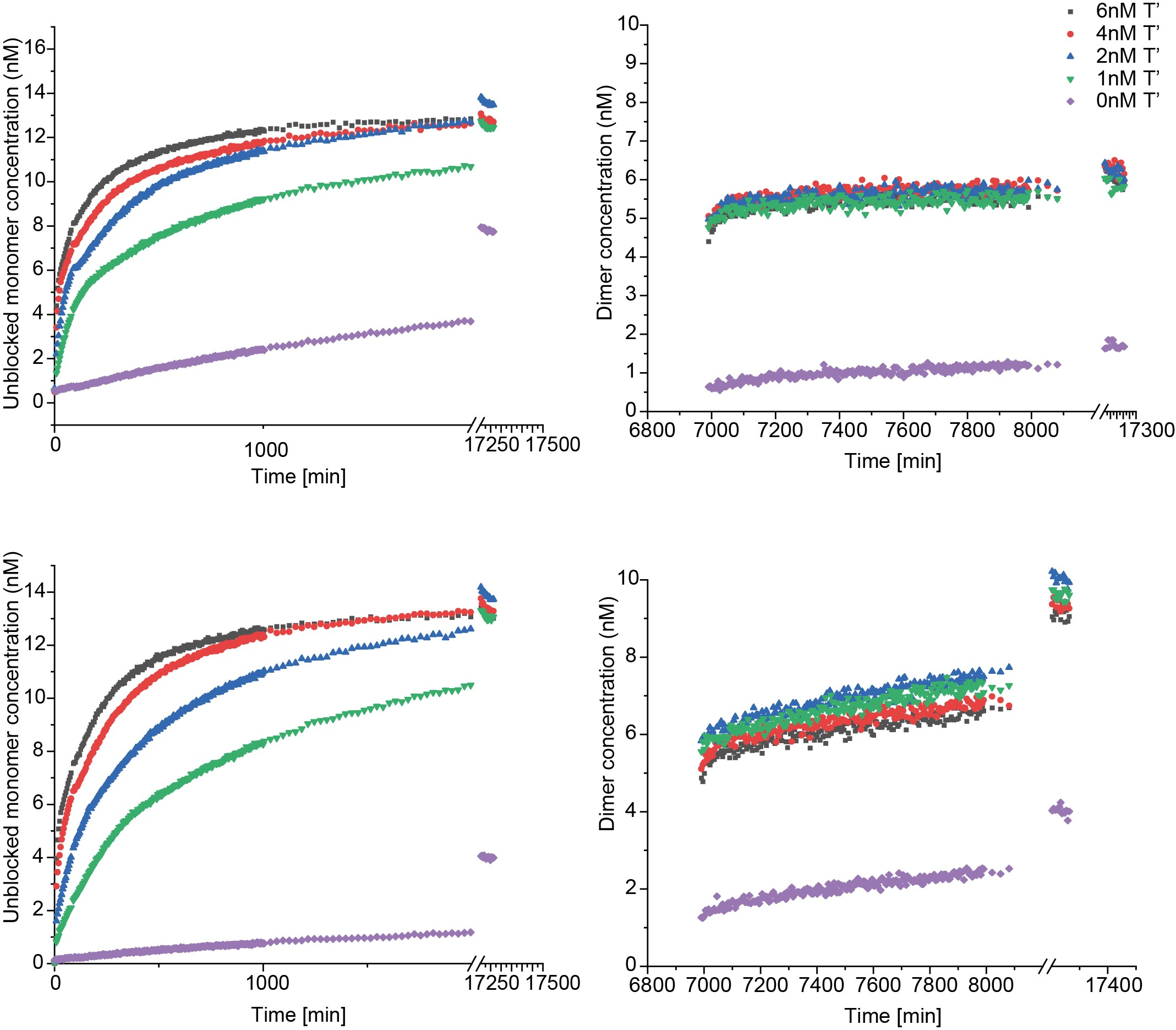

Figure S58: M’_1_N dimer formation from a mixture of M’ monomers in presence (top row) and absence (bottom row) of P’_7_

Reactions are triggered by adding 0-6 nM of *T’* into a solution containing 5 nM of each *M’_1_L’*, *M’_2_L’* and *M’_3_L’*, 50 or 0 nM *P’_7_*, and 10 nM *N*. The progress of the reactions was followed by monitoring the Cy3 channel (left column). When all the experiments reached a plateau at almost similar level of fluorescence signal intensities, 20 nM of external reporter RQ was added to the reaction to measure the amount of *M’_1_N* formed by monitoring the AlexaFluor-647 fluorescence signal (right column). The reporter was not added at t=0 as it induces leak in dimerization reaction. The data are processed as outlined in Supplementary note 15.

### Supplementary Figure 59: Formation of a *M’_2_N* dimer from a mixture of *M’_1_*, *M’_2_* and *M’_3_*, in the presence and absence of *P’_7_*

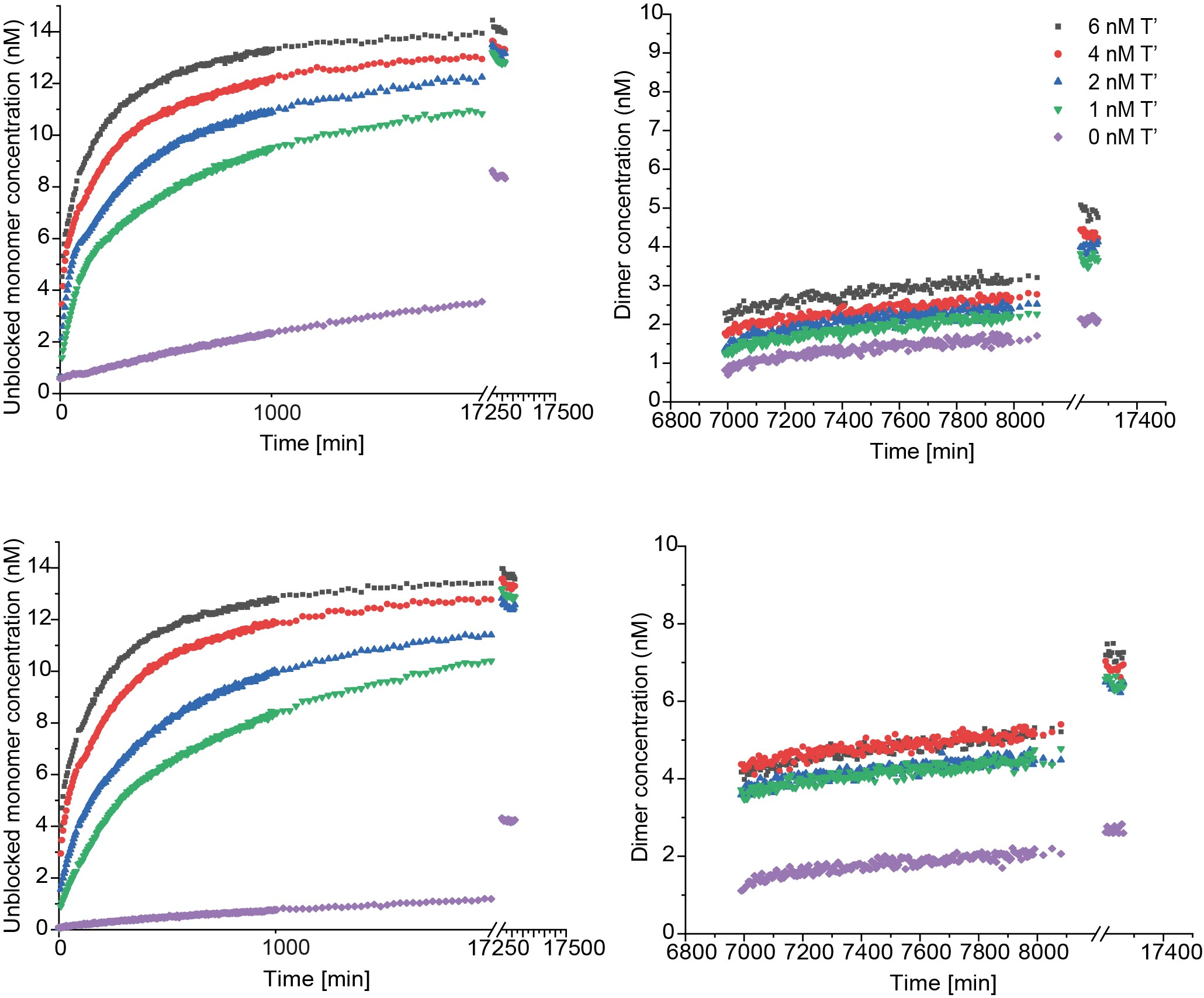

Figure S59: M’_2_N dimer formation from a mixture of M’ monomers in presence (top row) and absence (bottom row) of P’_7_

Reactions are triggered by adding 0-6 nM of *T’* into a solution containing 5 nM of each *M’_1_L’*, *M’_2_L’* and *M’_3_L’*, 50 or 0 nM *P’_7_*, and 10 nM *N*. The progress of the reactions was followed by monitoring the Cy3 channel (left column). When all the experiments reached a plateau at almost similar level of fluorescence signal intensities, 20 nM of external reporter *RQ* was added to the reaction to measure the amount of *M’_2_N* formed by monitoring the AlexaFluor-647 fluorescence signal (right column). The reporter was not added at t=0 as it induces leak in dimerization reaction. The data are processed as outlined in Supplementary note 15.

### Supplementary Figure 60: Formation of a *M’_3_N* dimer from a mixture of *M’_1_*, *M’_2_* and *M’_3_*, in the presence and absence of *P’_7_*

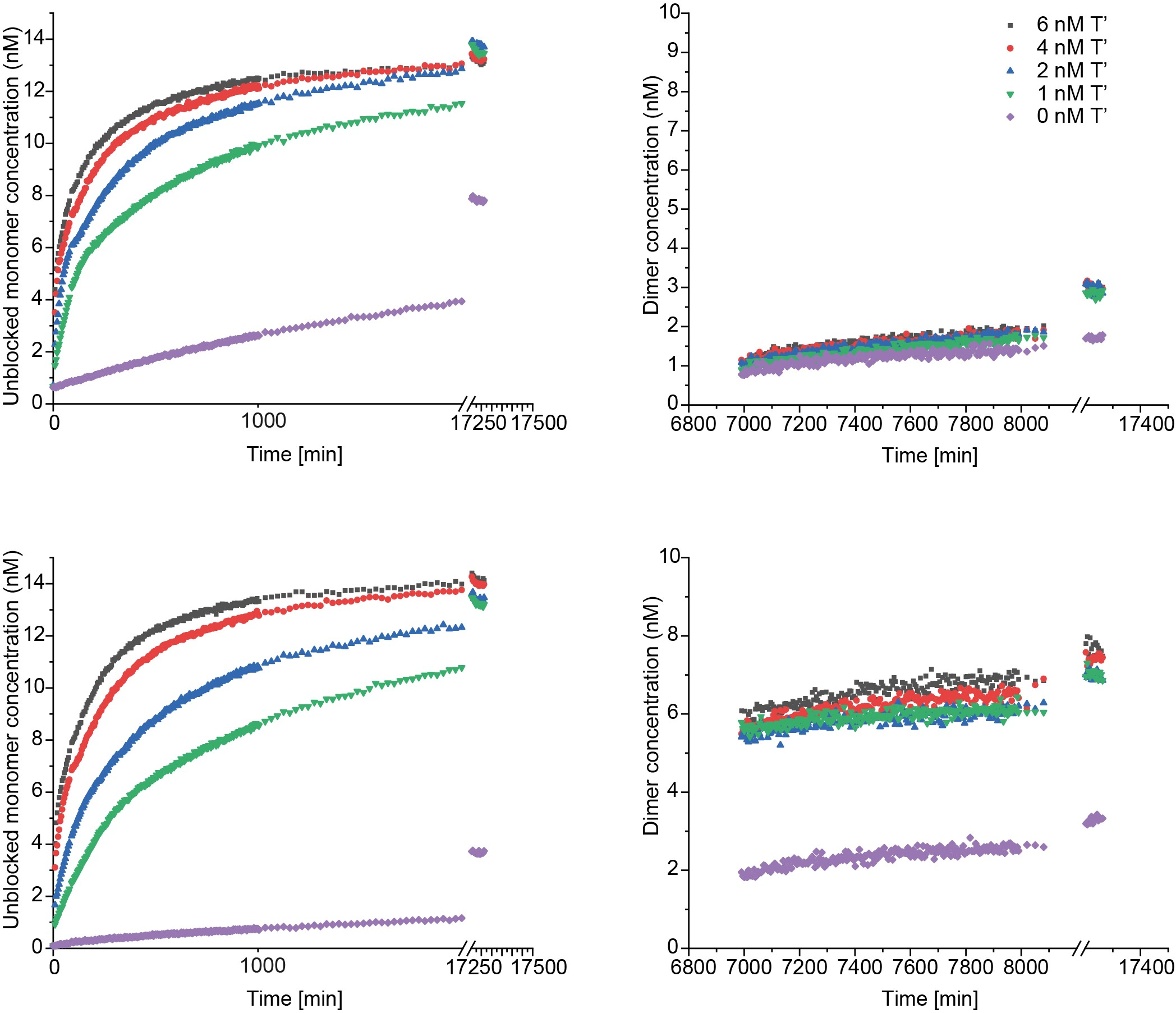

Figure S60: M’_3_N dimer formation from a mixture of M’ monomers in presence (top row) and absence (bottom row) of P’_7_

Reactions are triggered by adding 0-6 nM of *T’* into a solution containing 5 nM of each *M’_1_L’*, *M’_2_L’* and *M’_3_L’*, 50 or 0 nM *P’_7_*, and 10 nM *N*. The progress of the reactions was followed by monitoring the Cy3 channel (left column). When all the experiments reached a plateau at almost similar level of fluorescence signal intensities, 20 nM of external reporter *RQ* was added to the reaction to measure the amount of *M’_3_N* formed by monitoring the AlexaFluor-647 fluorescence signal (right column). The reporter was not added at t=0 as it induces leak in dimerization reaction. The data are processed as outlined in Supplementary note 15.

### Supplementary Figure 61: SNP discrimination with varying concentrations of proofreader molecule

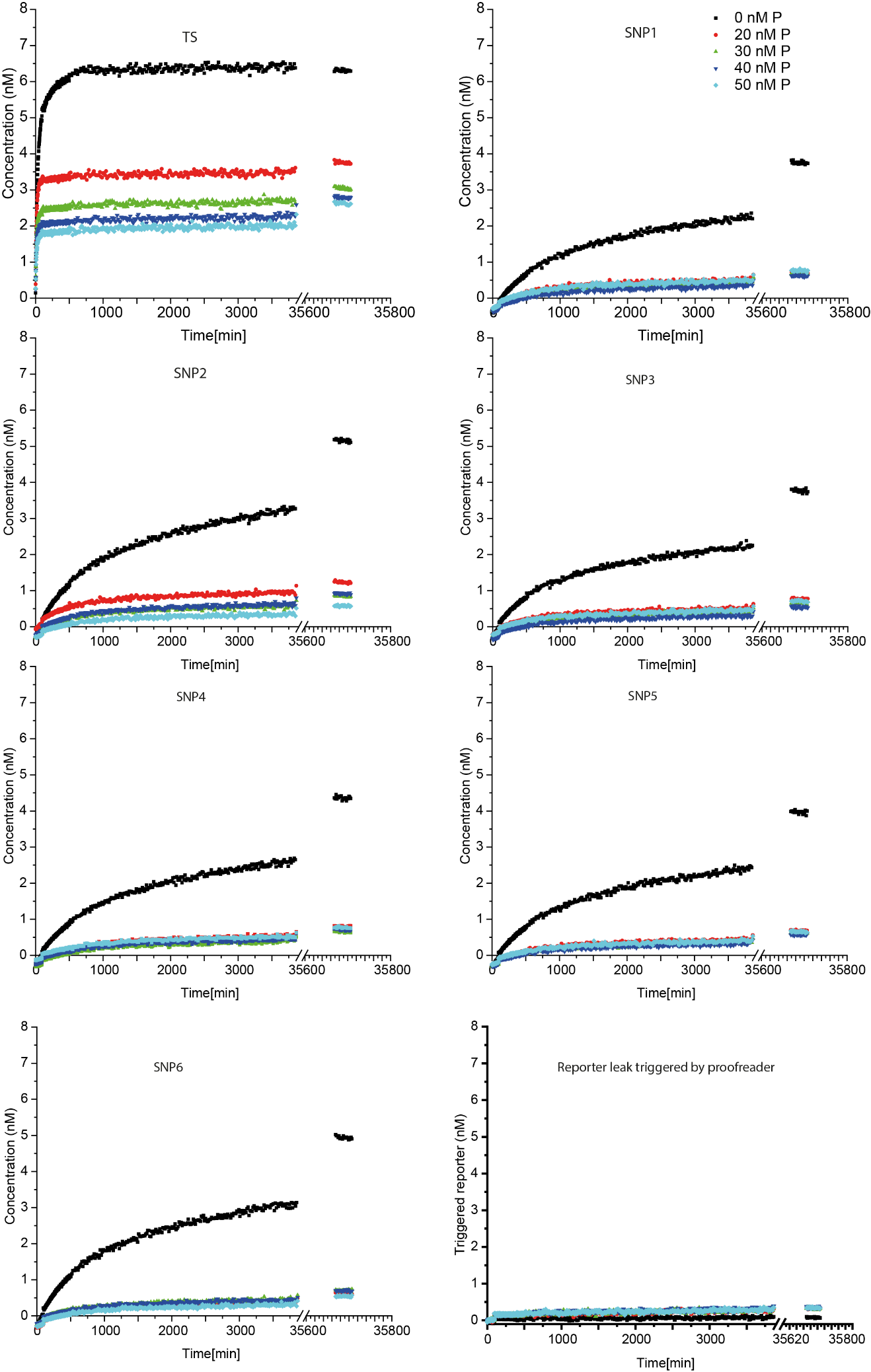

Figure S61: Concentration of candidate-probe complexes with different concentrations of proofreader.

10 nM *Probe-blocker* complex was added to a solution containing 8 nM candidate strand (*TS* or one of six different *SNPs*), 20 nM reporter complex, 20 nM sink complex, and 0-50 nM of proofreader strand. The desired signal is obtained by measuring the Cy3 fluorescence intensity for about 3 days. The experiments were measured again after four weeks to ensure that there was no triggering of the reporter even in the long term. The bottom right panel shows the direct triggering of the reporter by different concentrations of proofreader which are almost comparable to the SNP signals in presence of the proofreader. Raw data are processed as outlined in Supplementary note 16.

### Supplementary note 18: Specificity of dimerization process

The mean of final 100 measurements of dimer concentrations in Supplementary Figures S44-S52 are used to determine the concentration of *M’T’* complexes at the completion of reactions. The initial experimental conditions are: 8 nM *M’L’* complex, 10 nM *N*, 50 nM *P’_6­_*, and 2 nM *T’*.

The overall reaction is given as

M’L’ + T’ ⇄ M’T’ + L’;

M’T’ + N ⇄ M’N’ + T’;

M’T’ + P’ ⇄ M’P’ + T’.

[*M’N*] is inferred from AlexaFluor647 fluorescence signal of the external reporter *R_1_Q_1_*.

The discrimination factor D is calculated for a monomer variant *i* as

D_1:i_ = [*M’_1_N*]/[*M’_i_N*],

since the total concentration of *M’* monomers are the same in all experiments.

In the following table, we have listed the specificity of dimer formation for *M’_2_* and *M’_3_* in presence or absence of three different proofreaders where the reactions were triggered by different concentrations of *T’*.

Table S 12: Discrimination factors of monomers containing a single mismatch at the toehold domain in a dimerisation reaction

| **Proofreader** | **[T']** | **With Proofreader** | | | **Without proofreader** | | |
| --- | --- | --- | --- | --- | --- | --- | --- |
|  |  | **D_1:1_** | **D_1:2_** | **D_1:3_** | **D_1:1_** | **D_1:2_** | **D_1:3_** |
| ***P'_6_*** | **6nM** | 1.00 | 2.17 | 3.68 | 1.00 | 1.00 | 1.05 |
|  | **4nM** | 1.00 | 2.83 | 3.66 | 1.00 | 0.89 | 1.00 |
|  | **2nM** | 1.00 | 3.00 | 4.20 | 1.00 | 0.87 | 0.92 |
|  | **1nM** | 1.00 | 3.62 | 4.45 | 1.00 | 1.12 | 1.01 |
| ***P'_7_*** | **6nM** | 1.00 | 2.21 | 3.93 | 1.00 | 0.92 | 0.92 |
|  | **4nM** | 1.00 | 2.28 | 3.91 | 1.00 | 0.98 | 0.96 |
|  | **2nM** | 1.00 | 2.71 | 3.91 | 1.00 | 0.98 | 0.93 |
|  | **1nM** | 1.00 | 3.30 | 4.46 | 1.00 | 1.01 | 0.90 |
| ***P'_8_*** | **6nM** | 1.00 | 0.84 | 1.13 | 1.00 | 0.84 | 1.16 |
|  | **4nM** | 1.00 | 0.86 | 1.16 | 1.00 | 0.80 | 1.09 |
|  | **2nM** | 1.00 | 0.89 | 1.23 | 1.00 | 0.83 | 1.03 |
|  | **1nM** | 1.00 | 0.96 | 1.37 | 1.00 | 0.85 | 1.05 |

### Supplementary note 19: Specificity of SNP identification

The mean of final 20 data points shown in figure S61 are used as the final concentration of the probe-candidate-reporter complexes at the completion of reactions. Initial experimental conditions are: 8 nM *TS* or *SNP*, 10 nM Probe-Lock complex, 0-50 nM *P_snp_*, 20 nM of *Reporter* and *Sink* complexes.

The overall reaction is given as:

TS + Probe-Lock ⇄ TS-Probe + Lock;

TS-Probe + Reporter-Quencher ⇄ TS-Probe-Reporter + Quencher;

TS-Probe + P ⇄ Probe-P + TS and

SNP + Probe-Lock ⇄ SNP-Probe + Lock;

SNP-Probe + Reporter-Quencher ⇄ SNP-Probe-Reporter + Quencher;

SNP-Probe + P ⇄ Probe-P + SNP.

*TS-Probe-Reporter* or *SNP-Probe-Reporter* concentrations were inferred from the Cy3 signals of the complexes.

The discrimination factor D is calculated for an *SNP* as

D_TS_:_SNP_ = [*TS-Probe-Reporter*]/[*SNP-Probe-Reporter*],

since the total concentration of candidate strands are the same in all experiments.

The following table shows the discrimination factors for the SNPs without or with different concentrations of the proofreader.

Table S 13: Discrimination factors of candidate SNP strands containing a single mismatch at the branch migration domain

| **[*P_snp_*]** | **D_TS:SNP1_** | **D_TS:SNP2_** | **D_TS:SNP3_** | **D_TS:SNP4_** | **D_TS:SNP5_** | **D_TS:SNP6_** |
| --- | --- | --- | --- | --- | --- | --- |
| **0 nM** | 1.70 | 1.23 | 1.69 | 1.46 | 1.60 | 1.28 |
| **20 nM** | 5.94 | 3.22 | 5.46 | 5.23 | 6.29 | 6.89 |
| **30 nM** | 5.37 | 3.95 | 5.27 | 5.32 | 5.86 | 4.89 |
| **40 nM** | 5.14 | 3.29 | 6.08 | 4.39 | 5.69 | 4.52 |
| **50 nM** | 3.80 | 5.29 | 4.14 | 3.76 | 4.54 | 5.68 |

### Supplementary Figure 62: Reporter rates for *TS-Probe* and *SNP-Probe* complexes

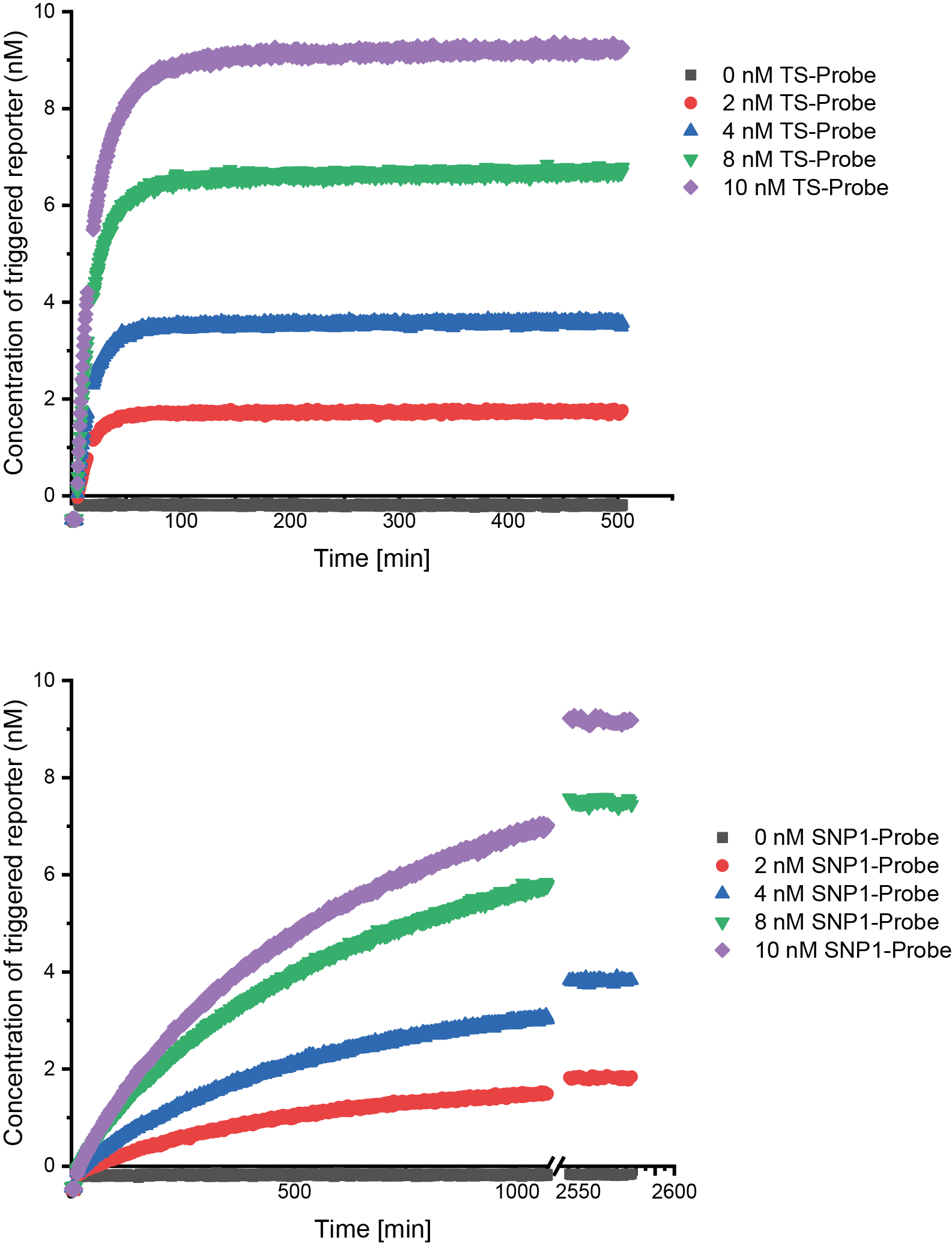

Figure S62: Reporter characterisation for TS-Probe and SNP-Probe complexes

0-10 nM of pre-annealed *TS-Probe or SNP-Probe* complexes were added to 20 nM of corresponding reporters, and fluorescence signals were measured in Cy3 channel. Cy3 reading showed the progress of the. Filled coloured symbols are experimentally obtained data. These data are the result of processing raw data as outlined in Supplementary note 16. Although the reporter rates are different, they are significantly faster than the reaction kinetics of the SNP identification as seen is Supplementary Fig. S61.

### Supplementary Figure 63: Initial reaction rate estimation for the SNP detection system

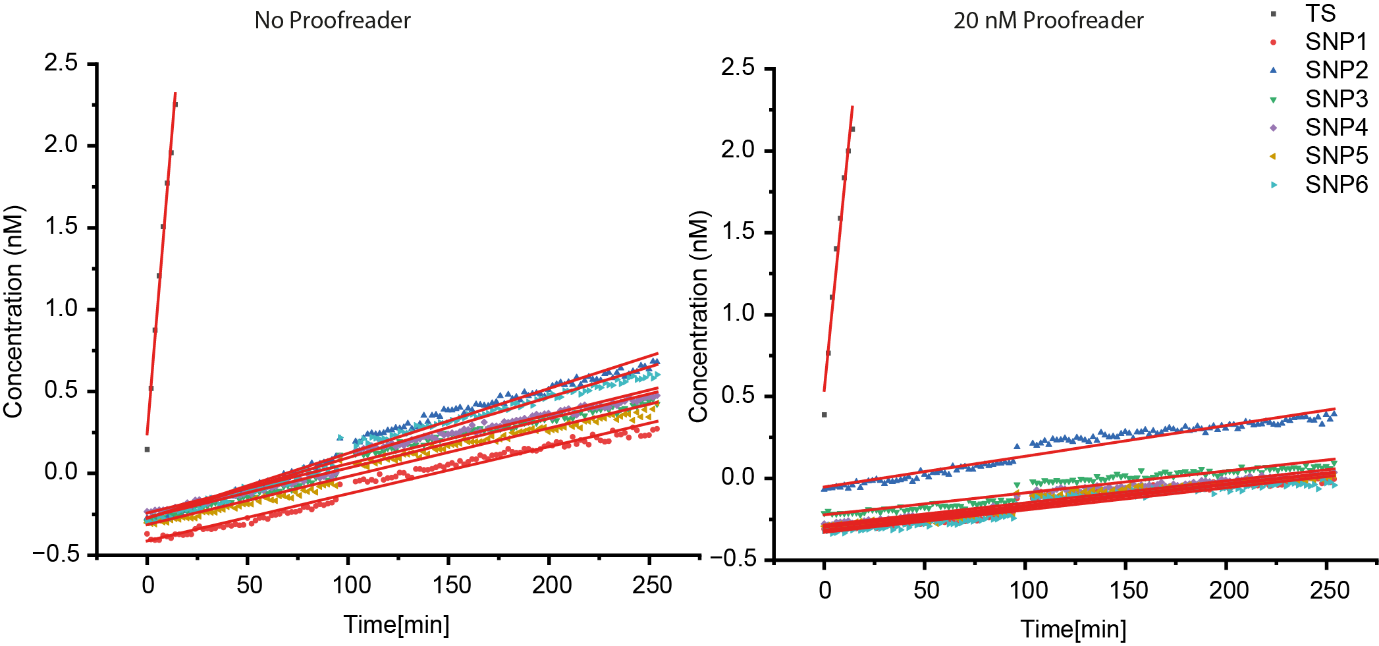

Figure S63: Estimates of initial reaction rates in SNP detection system with and without 20 nM proofreader.

For the *TS* strand, initial data of Figure S60 up to 15 minutes have been fit because of its linearity. For the SNPs, data up to 250 minutes have been used for the fit. The fit is generated by using Origin 2022 software’s linear fit option. The initial rates of the reactions are given in the table S12.

Table S 14: Initial rates of the SNP detection reaction with or without 20 nM Proofreader

| **Strand** | **Initial rate with proofreading (nM min^-1^)** | **Initial rate without proofreading**  **(nM min^-1^)** |
| --- | --- | --- |
| *TS* | 0.12354 ± 0.00831 | 0.14845 ± 0.00577 |
| *SNP1* | 0.00138 ± 3.68565E-5 | 0.00288 ± 3.98625E-5 |
| *SNP2* | 0.00188 ± 4.22206E-5 | 0.00396 ± 4.8246E-5 |
| *SNP3* | 0.00134 ± 3.69041E-5 | 0.00302 ± 4.62678E-5 |
| *SNP4* | 0.00135 ± 3.44479E-5 | 0.003 ± 4.10845E-5 |
| *SNP5* | 0.00133 ± 3.40065E-5 | 0.00294 ± 3.98922E-5 |
| *SNP_6_* | 0.00136 ± 3.61932E-5 | 0.0037 ± 4.54134E-5 |

### Supplementary Figure 64: Experiments with the TS and control-SNP with same reporter toehold

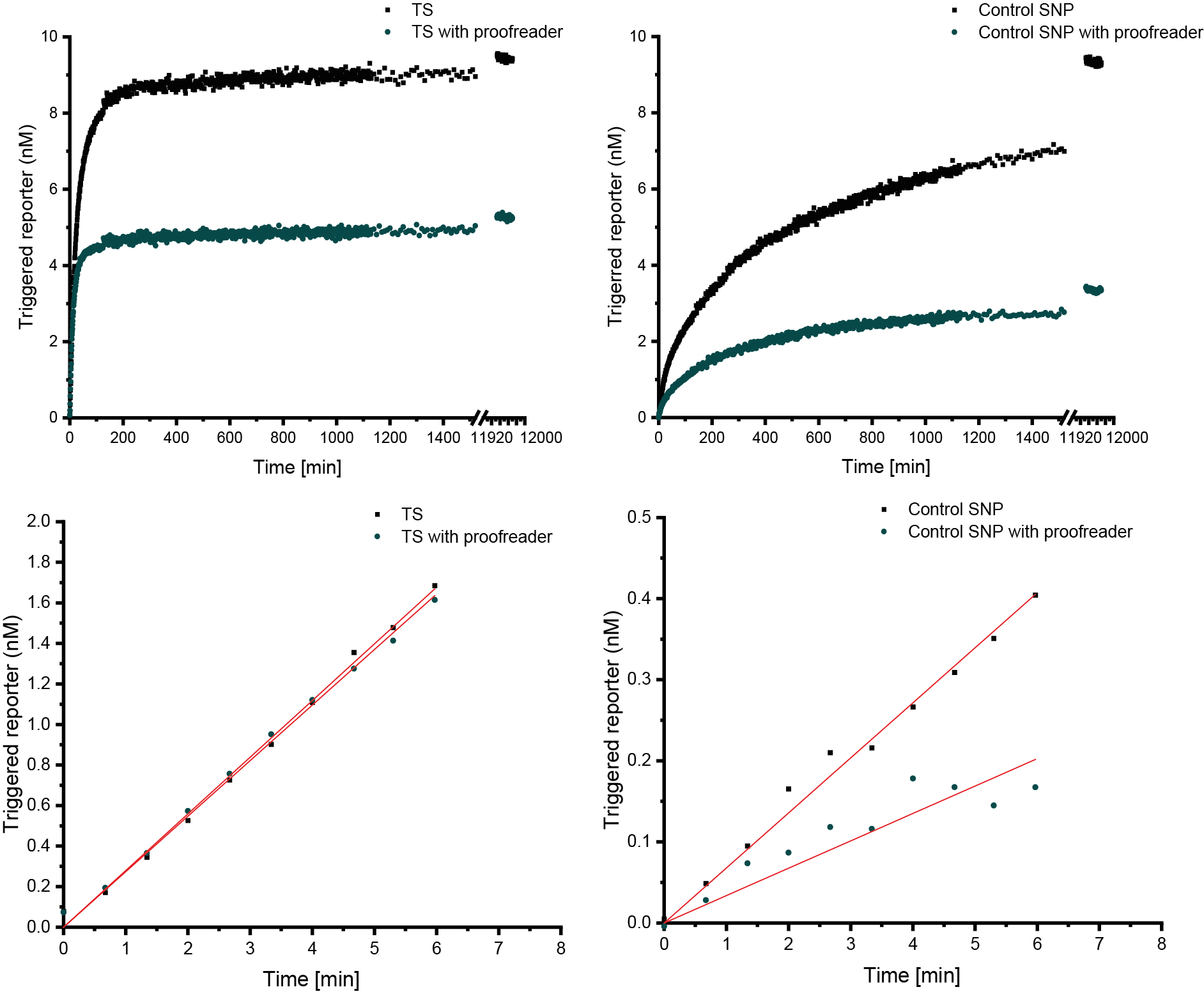

Figure S 64: Comparison of TS and control SNP reactions with same toehold

A new SNP strand “control SNP” is designed to where the reporter toehold has the same nucleotide sequence as WT, and the rest is same as SNP1. Although the control-SNP reaction rates are faster than the SNPs shown in Fig S61, the final levels attained by control-SNP is lower than TS in presence of 20 nM proofreader.

10 nM *Probe-blocker* complex was added to a solution containing 8 nM candidate strand (*TS* or Control-*SNP*), 20 nM reporter complex, 20 nM sink complex, and 0-50 nM of proofreader strand. The desired signal is obtained by measuring the Cy3 fluorescence intensity for about 3 days. The experiments were measured again after four weeks to ensure that there was no triggering of the reporter even in the long term. The bottom right panel shows the direct triggering of the reporter by different concentrations of proofreader which are almost comparable to the SNP signals in presence of the proofreader. Raw data are processed as outlined in Supplementary note 16.

### Sequences

Colour legends:

**Blue: Template toehold**

**Green: Proofreader toehold**

**Red: Displacement domain**

**Black: Spacer or reporter assist domain**

**Purple: Reporter quencher duplex domain**

**Light green: Duplex domain within reporter**

**Yellow: Handhold**

**Dirty yellow: Extended domain for dimer reporters**

Table S 15: DNA oligos used in characterising proofreading mechanism

| Name | 5' label | Sequence | 3' label |
| --- | --- | --- | --- |
| T |  | TCCATT TCTACTCCATCTCATCCATCCTACCTCAC |  |
| Lock |  | TCTACTCCATCTCATCCATTCTACCTCAC TCATC TTT | IowaBlack FQ |
| M_1_ | Cy3 | TTT GATGA GTGAGGTAGGATGGATGAGGTGGAGTAGA AATGGA TA |  |
| M_2_ | Cy3 | TTT GATGA GTGAGGTAGGATGGATGAGGTGGAGTAGA AATCGA AG |  |
| M_3_ | Cy3 | TTT GATGA GTGAGGTAGGATGGATGAGGTGGAGTAGA GATGGA GC |  |
| P_6_ |  | CAATCAACTAAACCTAAACG TCTACTCCATCTCATCCATCCTACCTCAC TCATC |  |
| P_7_ |  | CAATCAACTAAACCTAAACG CTACTCCATCTCATCCATCCTACCTCAC TCATC |  |
| P_8_ |  | CAATCAACTAAACCTAAACG TACTCCATCTCATCCATCCTACCTCAC TCATC |  |

Table S 16: Reporter strands used for monomer-proofreader waste complexes

| Name | 5' label | Sequence | 3' label |
| --- | --- | --- | --- |
| Rcomp_AF |  | GGAGGTGATATGAGGTGG CGTTTAGGTTTAGTTGATTG TTT | AlexaFluor-647 |
| Q_2_ | IowaBlack RQ | TTT CAATCAACTAAACCTAAAC |  |
| r_1-6_ |  | TA TCCATT CCACCTCATATCACCTCC |  |
| r_1-7_ |  | AT CCATTT CCACCTCATATCACCTCC |  |
| r_1-8_ |  | TC CATTTC CCACCTCATATCACCTCC |  |
| r_2-6_ |  | CT TCGATT CCACCTCATATCACCTCC |  |
| r_2-7_ |  | TT CGATTT CCACCTCATATCACCTCC |  |
| r_2-8_ |  | TC GATTTC CCACCTCATATCACCTCC |  |
| r_3-6_ |  | GC TCCATC CCACCTCATATCACCTCC |  |
| r_3-7_ |  | CT CCATCT CCACCTCATATCACCTCC |  |
| r_3-8_ |  | TC CATCTC CCACCTCATATCACCTCC |  |

Table S 17: DNA oligos used for the dimerization experiments

| Name | 5' label | Sequence | 3' label |
| --- | --- | --- | --- |
| T’ |  | TCCATT TCACTCAGTCCATCCCATCCACATCTTCA TT CTTATTCT |  |
| P’_6_ |  | CAATCAACTAAACCTAAACG TCACTCAGTCCATCCCATCCACATCTTCA CCATC |  |
| P’_7_ |  | CAATCAACTAAACCTAAACG CACTCAGTCCATCCCATCCACATCTTCA CCATC |  |
| P’_8_ |  | CAATCAACTAAACCTAAACG ACTCAGTCCATCCCATCCACATCTTCA CCATC |  |
| N |  | ACATACAGAT AGAATAAG TCACTCAGTCCATCGCATCCAGATCTTCA CC |  |
| M’_3_ | Cy3 | TTT GATGG TGAAGATCTGGATGCGATGGACTGAGTGA GATGGA GC |  |
| M’_2_ | Cy3 | TTT GATGG TGAAGATCTGGATGCGATGGACTGAGTGA AATCGA AG |  |
| M’_1_ | Cy3 | TTT GATGG TGAAGATCTGGATGCGATGGACTGAGTGA AATGGA TA |  |
| L’ |  | TCACTCACTCCATCCCATCCACATCTTC ACCATC TTT | IB-FQ |
| RcompP |  | GGAGGTGATATGAGGTGG CTTATTCT ATCTGTATGT TTT | AlexaFluor-647 |
| r_3_ |  | GCTCCATC CCACCTCATATCACCTCC |  |
| r_2_ |  | CTTCGATT CCACCTCATATCACCTCC |  |
| r_1_ |  | TATCCATT CCACCTCATATCACCTCC |  |
| Q1 | IB-RQ | TTT ACATACAGAT ACAATAAG |  |

Colour legends:

**Blue: Template toehold**

**Green: Proofreader toehold**

**Red: Displacement domain (single nucleotide mutations in black)**

**Dark blue: Reporter quencher duplex domain**

**Purple: Duplex domain within reporter or sink complex**

**Yellow: Flanking reporter domain**

**Light green: Flanking domain for sink complex**

Table S 18: DNA oligos used for SNP detection experiments

| Name | 5' label | Sequence | 3' label |
| --- | --- | --- | --- |
| TS |  | CGCCAC TCTACC CTCACCTCTAACCTCCACCTCCATACCTC |  |
| SNP3 |  | CATCAC TCTACC CTCACCTCTAACCTCCACCTCGATACCTC |  |
| SNP2 |  | CATCAC TCTACC CTCACCTCTAACCTGCACCTCCATACCTC |  |
| SNP1 |  | CATCAC TCTACC CTCACCTGTAACCTCCACCTCCATACCTC |  |
| SNP4 |  | CATCAC TCTACC CTCACCTTTAACCTCCACCTCCATACCTC |  |
| SNP5 |  | CATCAC TCTACC CTCACCTCTAACCACCACCTCCATACCTC |  |
| SNP6 |  | CATCAC TCTACC CTCACCTCTAACCTCCACCTCCTTACCTC |  |
| Control SNP |  | CGCCAC TCTACC CTCACCTGTAACCTCCACCTCCATACCTC |  |
| P_snp_ |  | CCATCAAA CTCACCTCTAACCTCCACCTCCATACCTC ACCTC |  |
| Probe |  | GAGGT GAGGTATGGAGGTGGAGGTTAGAGGTGAG GGTAGA ATAGAGACAATAGGACGG |  |
| Lock |  | CTCACCTCTAACCTCCACCTCCATACCTC ACCTC |  |
| r_2_ (for SNPs) |  | TGGGATGAAGTGAGTGAG GTGATG |  |
| r_1_ (for TS) |  | TGGGATGAAGTGAGTGAG GTGGCG |  |
| Rcomp | Cy3 | TTT CCGTCCTATTGTCTCTAT CTCACTCACTTCATCCCA |  |
| Q |  | TAGAGACAATAGGACGG TTT | IB-FQ |
| S1 |  | GTGGAGGTGAGCGGTGGA TTTGATGG |  |
| Scomp29 |  | CCGTCCTATTGTCTCTAT TCTACC TCCACCGCTCACCTCCAC |  |
| Sq |  | GTAGA ATAGAGACAATAGGACGG |  |
